# Biodynamics: A novel quasi-first principles theory on the fundamental mechanisms of cellular function/dysfunction and the pharmacological modulation thereof

**DOI:** 10.1101/384719

**Authors:** Gianluca Selvaggio, Robert A Pearlstein

## Abstract

Cellular function depends on heterogeneous <u>dynamic</u> intra-, inter-, and supramolecular structure-function relationships. However, the specific mechanisms by which cellular function is transduced from molecular systems, and by which cellular dysfunction arises from molecular dysfunction are poorly understood. We proposed previously that cellular function manifests as a molecular form of analog computing, in which specific time-dependent state transition fluxes within sets of molecular species (“molecular differential equations” (MDEs)) are sped and slowed in response to specific perturbations (inputs). In this work, we offer a theoretical treatment of the molecular mechanisms underlying cellular analog computing (which we refer to as “biodynamics”), focusing primarily on non-equilibrium (dynamic) intermolecular state transitions that serve as the principal means by which MDE systems are solved (the molecular equivalent of mathematical “integration”). Under these conditions, bound state occupancy is governed by *k_on_* and *k_off_*, together with the rates of binding partner buildup and decay. Achieving constant fractional occupancy over time depends on: 1) equivalence between k_on_ and the rate of binding site buildup); 2) equivalence between *k_off_* and the rate of binding site decay; and 3) free ligand concentration relative to *k_off_/k_0n_* (n · K_d_, where n is the fold increase in binding partner concentration needed to achieve a given fractional occupancy). Failure to satisfy these conditions results in fractional occupancy well below that corresponding to n · Kd. The implications of biodynamics for cellular function/dysfunction and drug discovery are discussed.

## INTRODUCTION

We proposed in our previous work [1] that time-dependent cellular function is derived from a 5 molecular form of analog computing [2,3], in which sets of coupled ordinary differential equations and their integral solutions are modeled physically by changes in the rates of buildup and decay) among the populations of biomolecular species to/from specific intra- and intermolecular states (the hardware and software are one and the same) (Fig 1). We referred to these constructs as “molecular differential equations” (MDEs) [1], the major forms of which include:

**Intramolecular:**

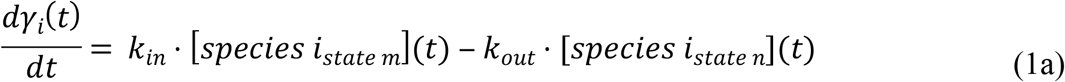

**Non-covalent Intramolecular:**

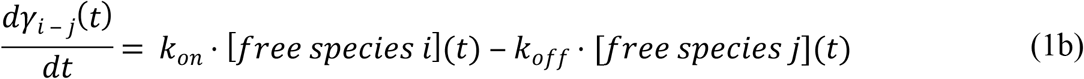

**Enzyme-substrate:**

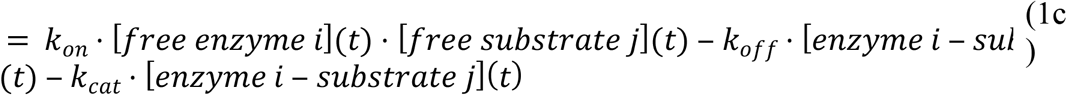

where *dγ_k_*(*t*)/*dt* represents the rates of non-equilibrium occupancy buildup and decay of state k from one or more predecessor to one or more successor states. State transition rates are governed by adjustable barriers originating from intra- or intermolecular interactions (which we referred to as “intrinsic rates”) and time-dependent changes in the concentrations or number densities of the participating species (which we referred to as “extrinsic rates”) [1]. Molecular populations “flow” over time in a transient (Markovian) fashion from one specific structural state to another (Fig 1) in response to production/degradation or translocation-driven changes in the levels of the participating species.

**Fig 1.**
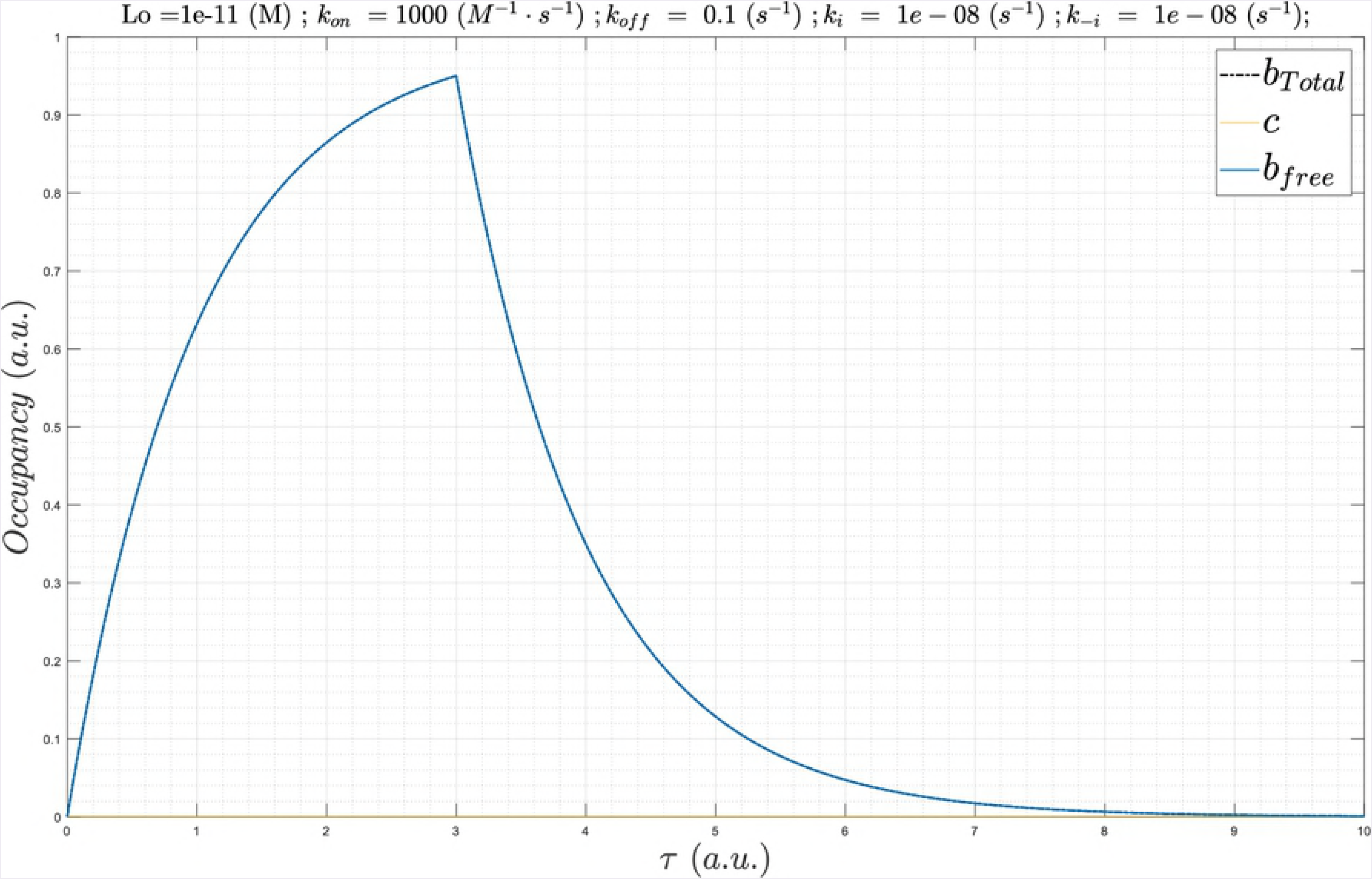
(A) Markovian state transition behavior exemplified for the human cardiac ether-a-go-go related gene (hERG) potassium channel between closed (C1, C2, and C3), open (O) and inactivated states (I) underlying the state probability curves in (B) [4]. The rate constants are labeled with Greek letters. (B) Molecular populations of hERG “flow” from one specific state to another based on intrinsic voltage-dependent rates of entry and exit. Multiple fluxes occur in parallel between specific states, speeding and slowing in response to dynamic perturbations. For example, the open (orange) and inactive (blue) states are fed by the closed state (magenta); the open and closed states are fed by the inactive state; and the inactive and closed states are fed by the open state. The time-dependent output of the overall system (i.e. the membrane potential in this case) is solved in cardiomyocytes via integration of the full complement of sodium, potassium, and calcium ion channel MDEs (which can be simulated using the O’Hara-Rudy model of the human cardiac AP [5]).

MDEs are “solved” by cells in aggregate for the corresponding time-dependent state occupancies of the participating species {γ_k_(t), k = 1, n} (e.g. dynamic ion channel states and currents, dynamic enzyme activation states, etc.), together with higher order (convergent) properties that we refer to as Γ_a_(t) (Fig 2). {γ_k_(t), k = 1, n} and Γ_a_(t) manifest as transient stimulus-response-driven “action potential-like” waveforms, in which the intrinsic or extrinsic rates of one or more MDEs are sped or slowed in response to Γ_a_(t). [1].

**Fig 2.**
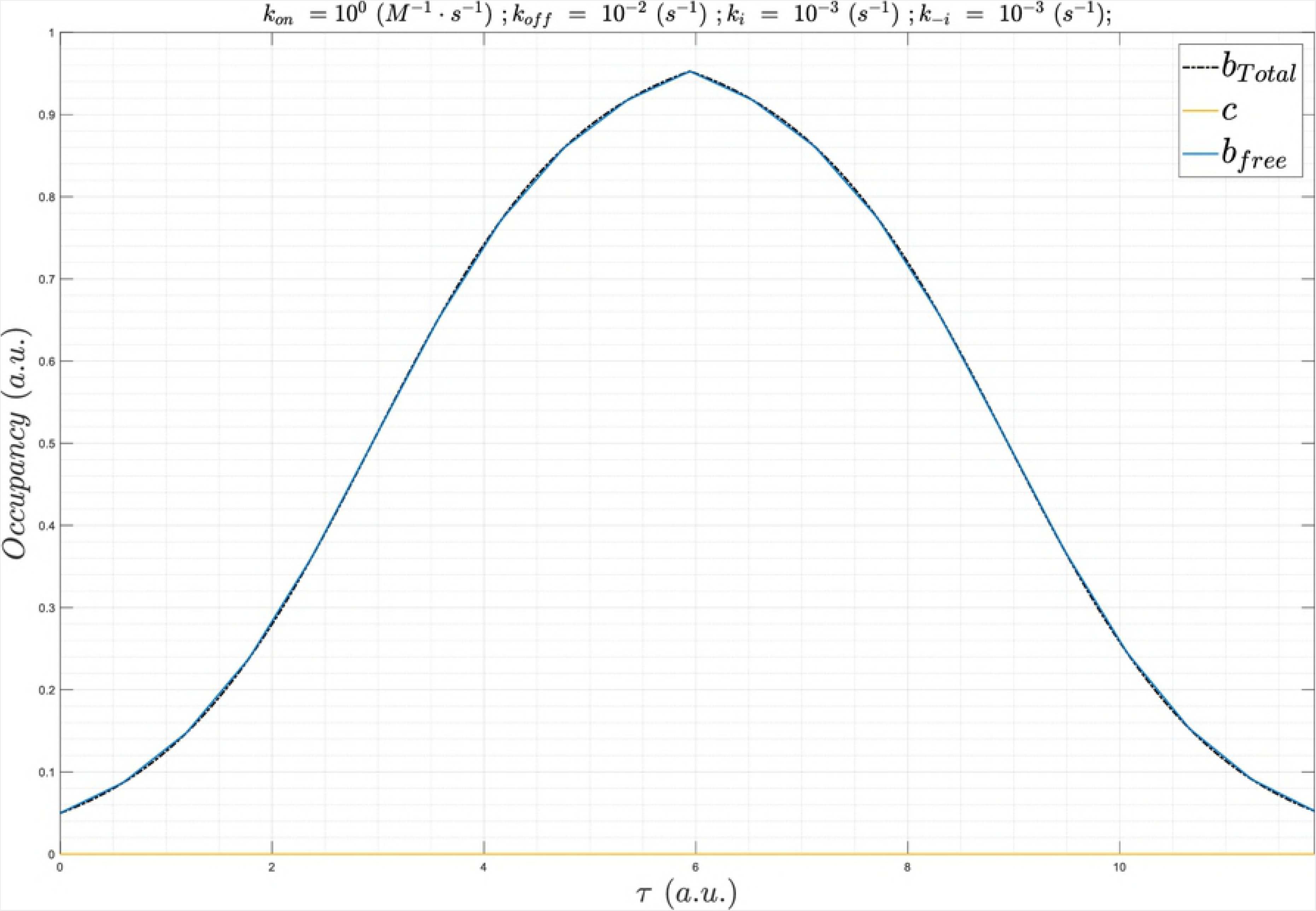
Cellular analog computing is based on the transduction of molecular state transitions into cellular function. MDEs {dγ_k_(t)/dt, k = 1, n} are “solved” in aggregate for the corresponding time-dependent state occupancies of the participating species {γ_k_(t), k = 1, n} (e.g. dynamic intra- and intermolecular states), together with higher order properties that we refer to as Γ_a_(t). The underlying MDEs accelerate or decelerate recursively in response to Γ_a_(t) according to their specific response mechanisms.

Building on our previous work [1] and that of others [2,6–11], we have developed a comprehensive multi-scale (atomistic ➔ molecular systems) first principles theory on the basic mechanisms of cellular function that we refer to as “biodynamics” (Fig 3).

**Fig 3.**
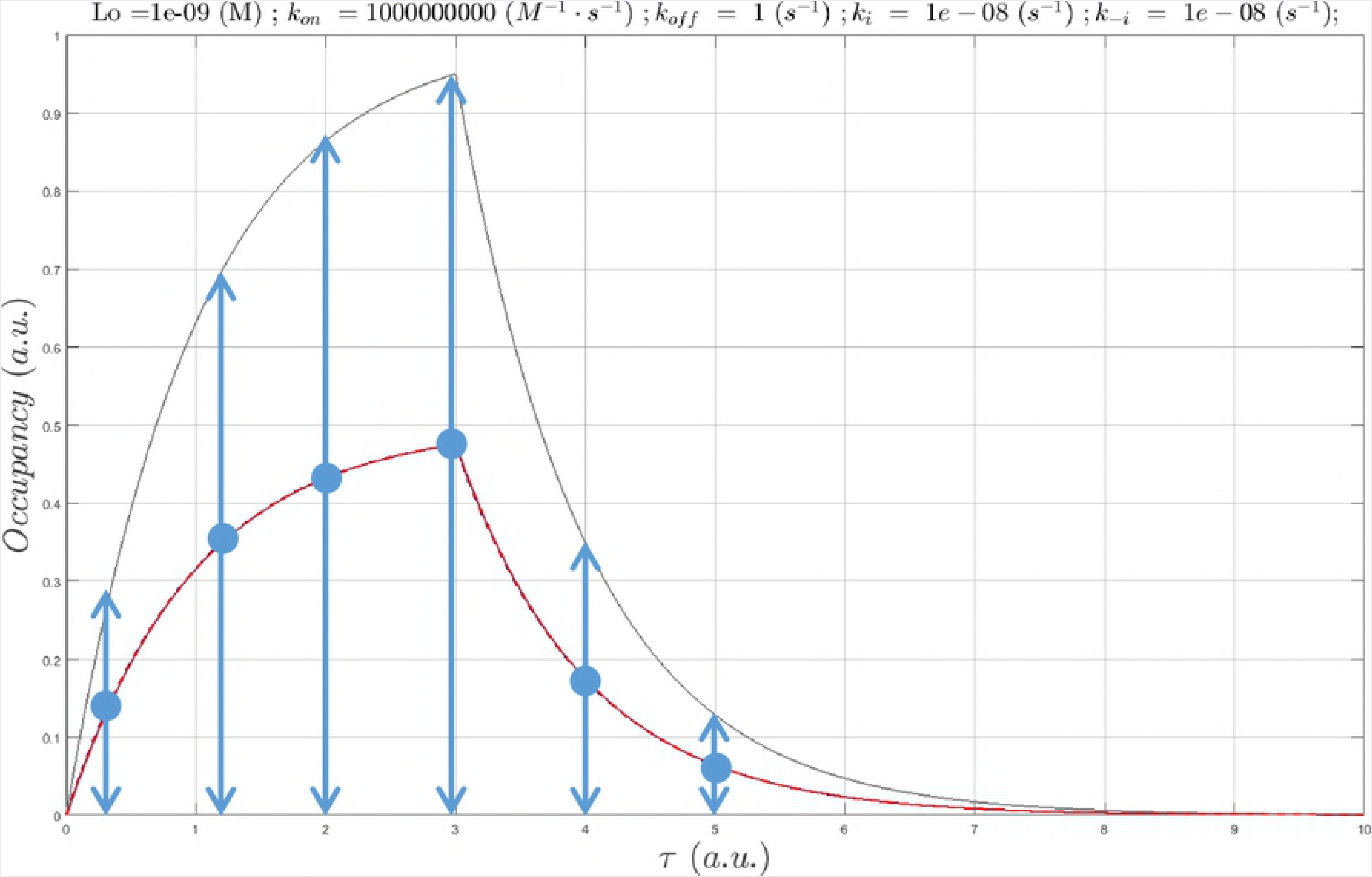
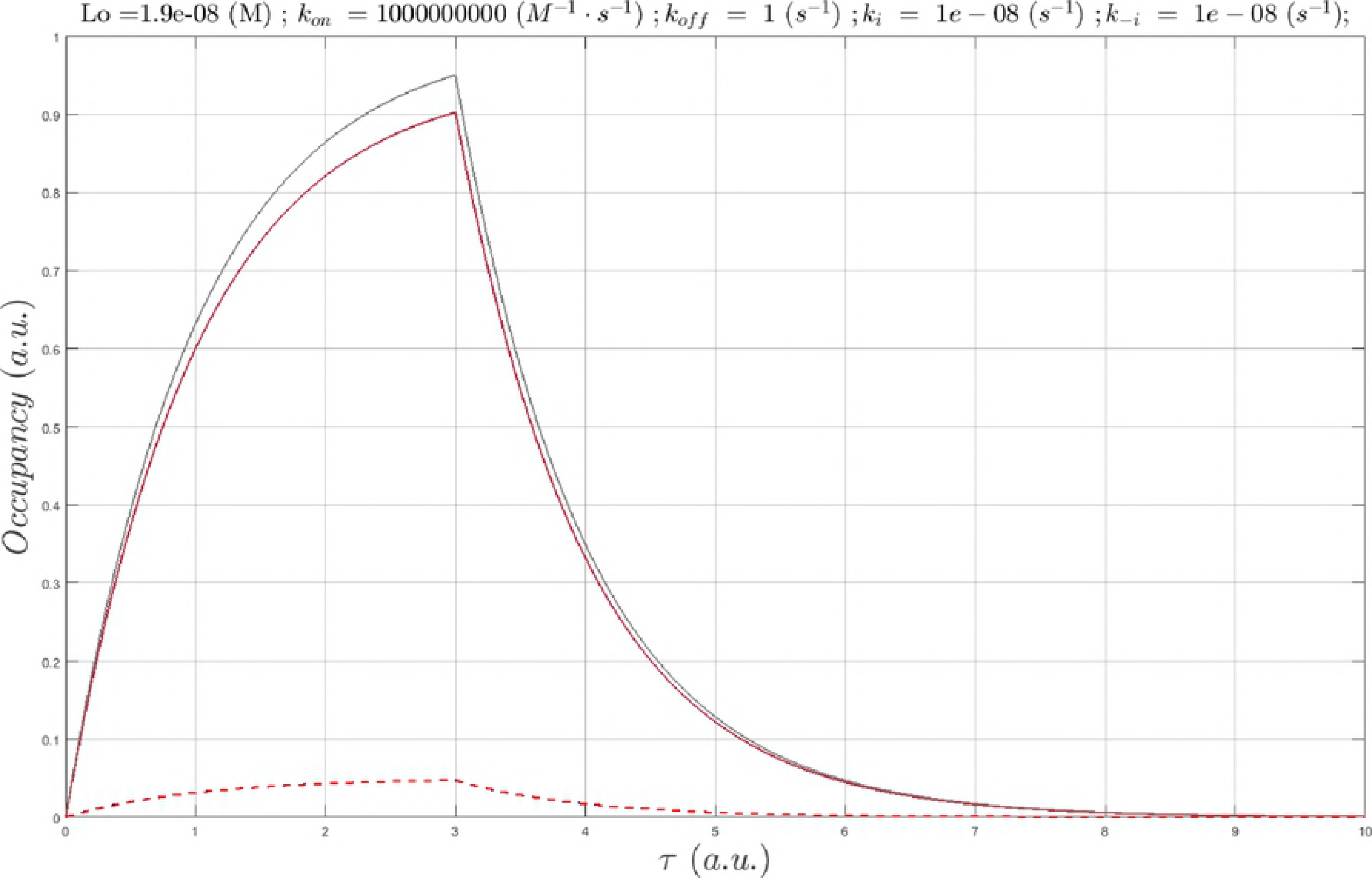
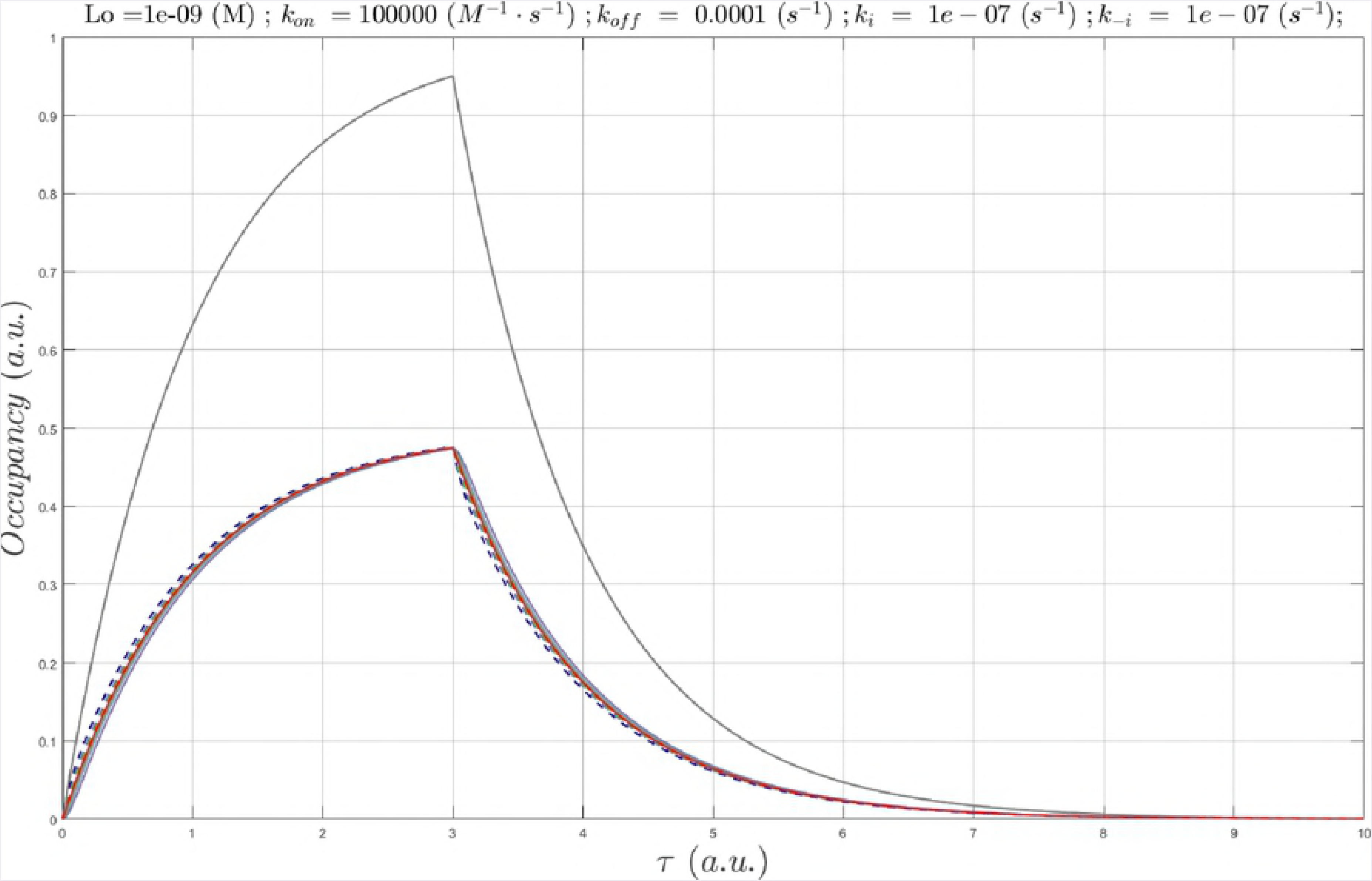
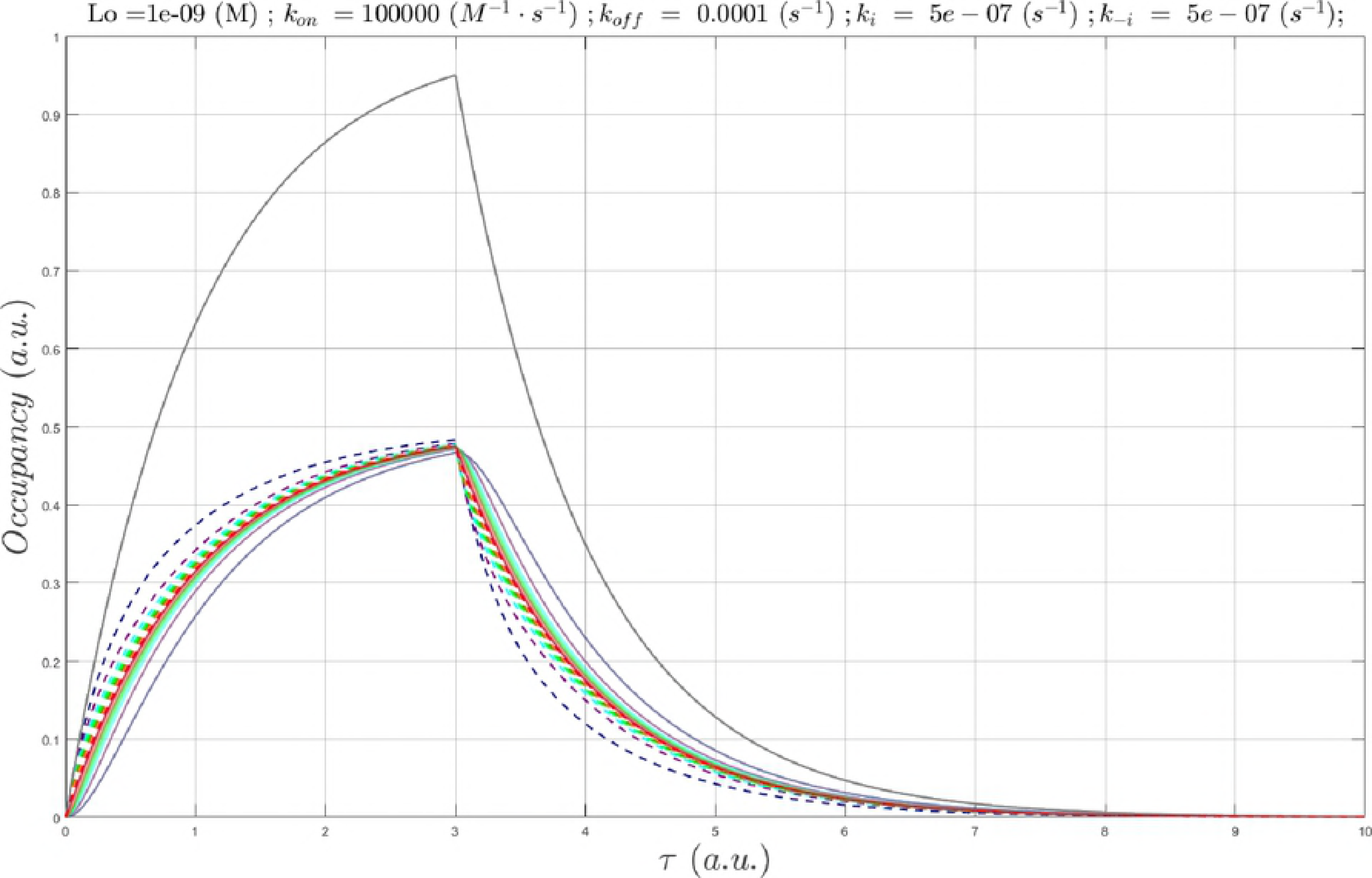
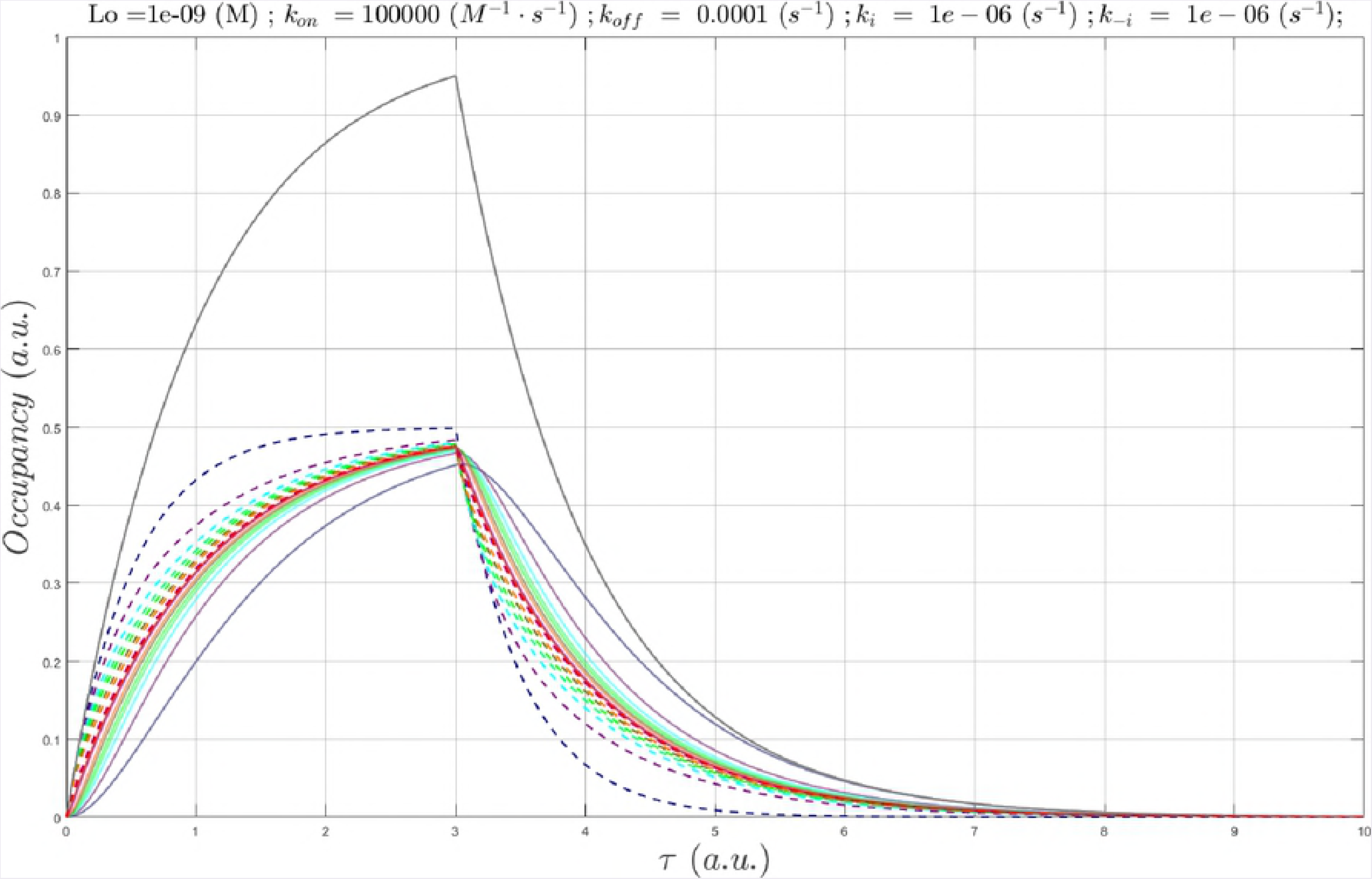
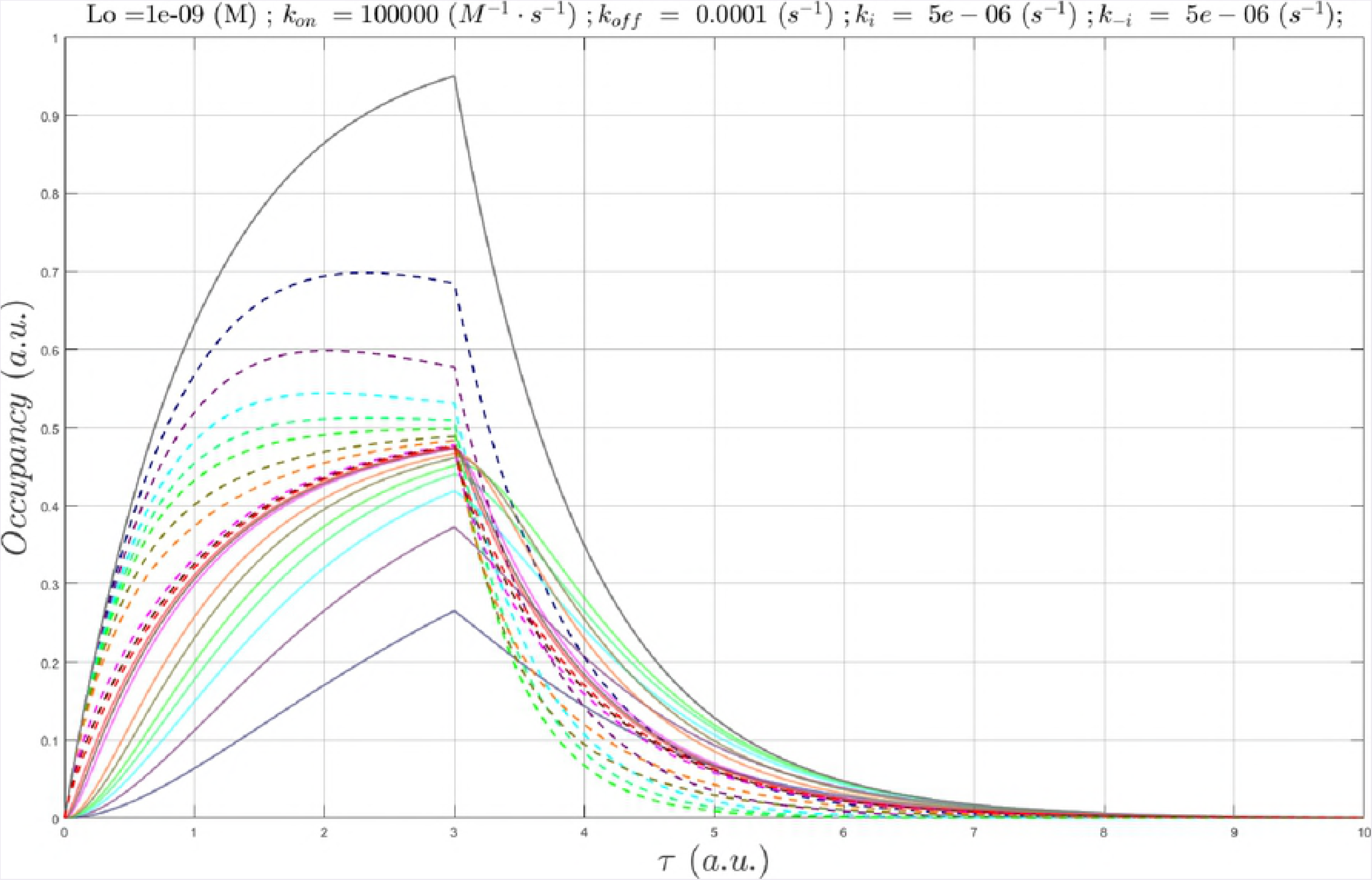
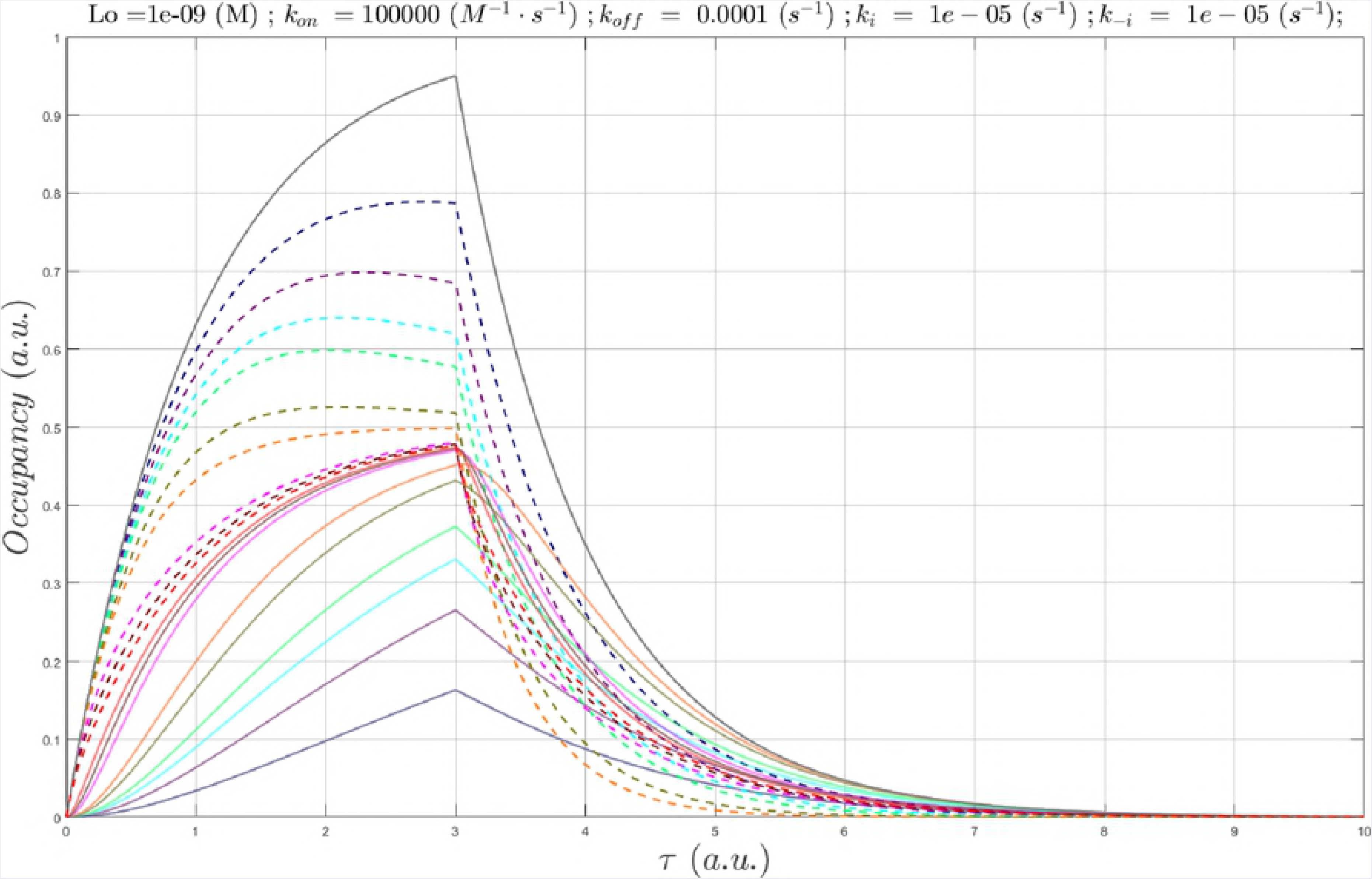
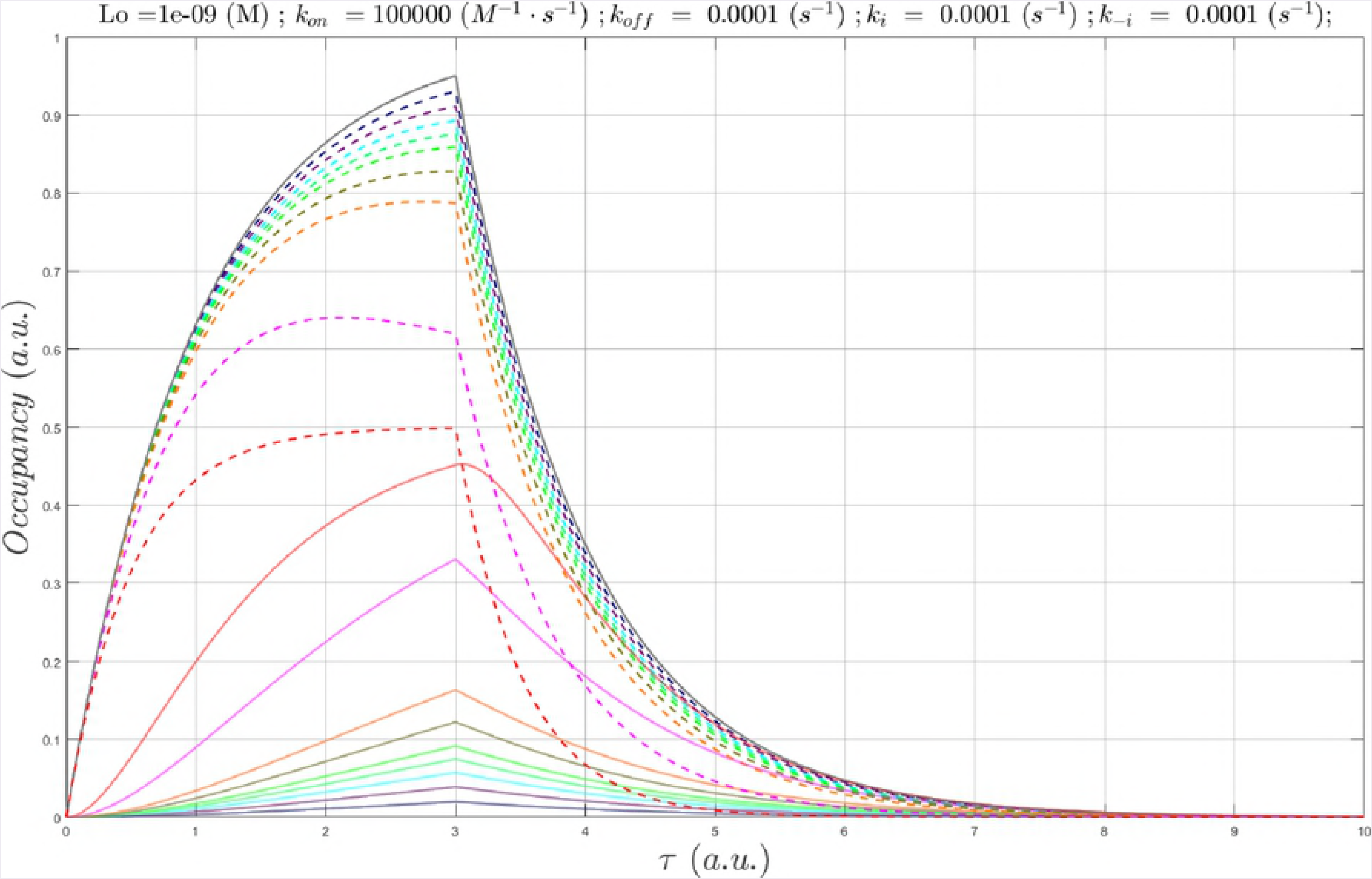
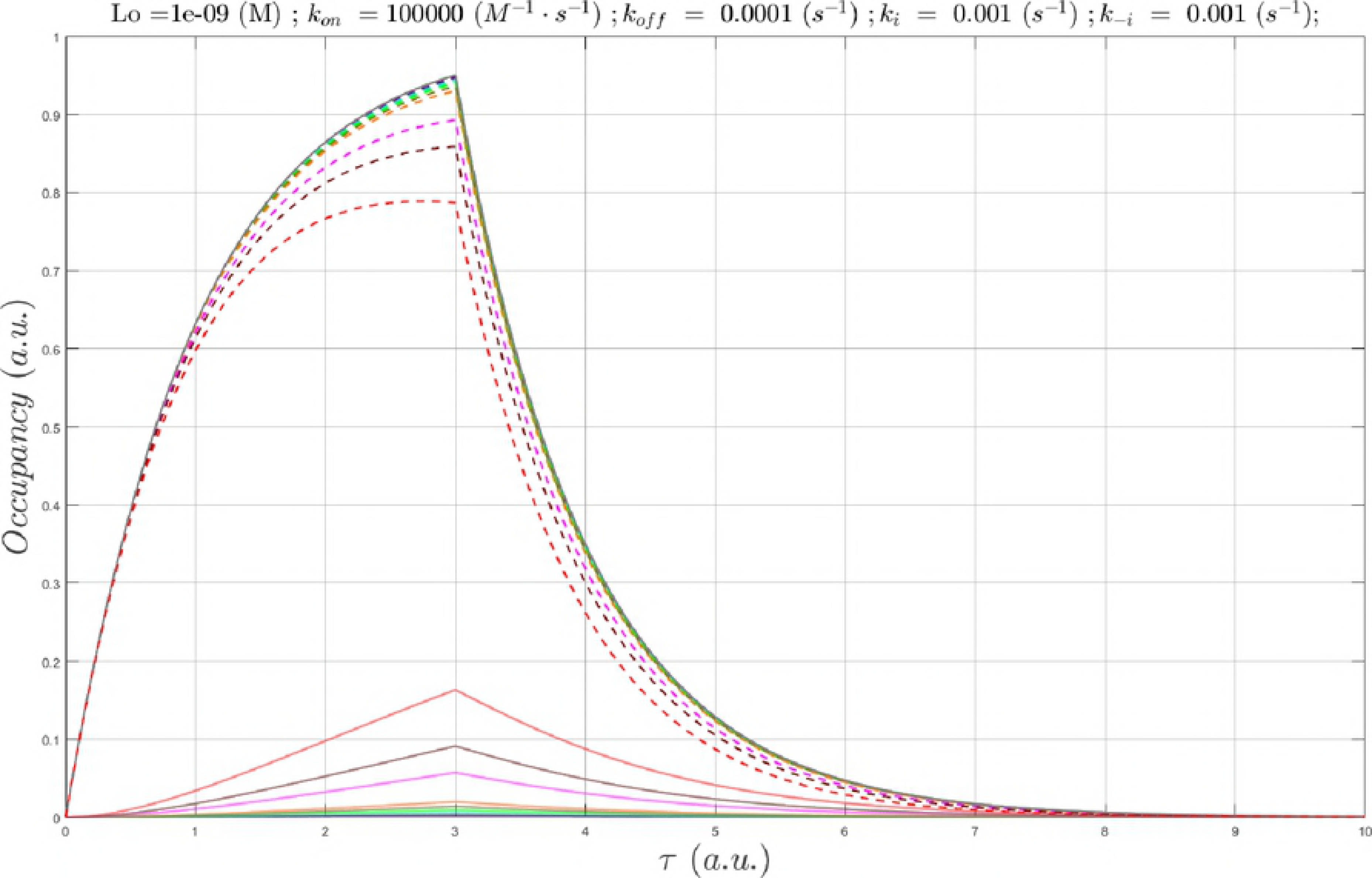
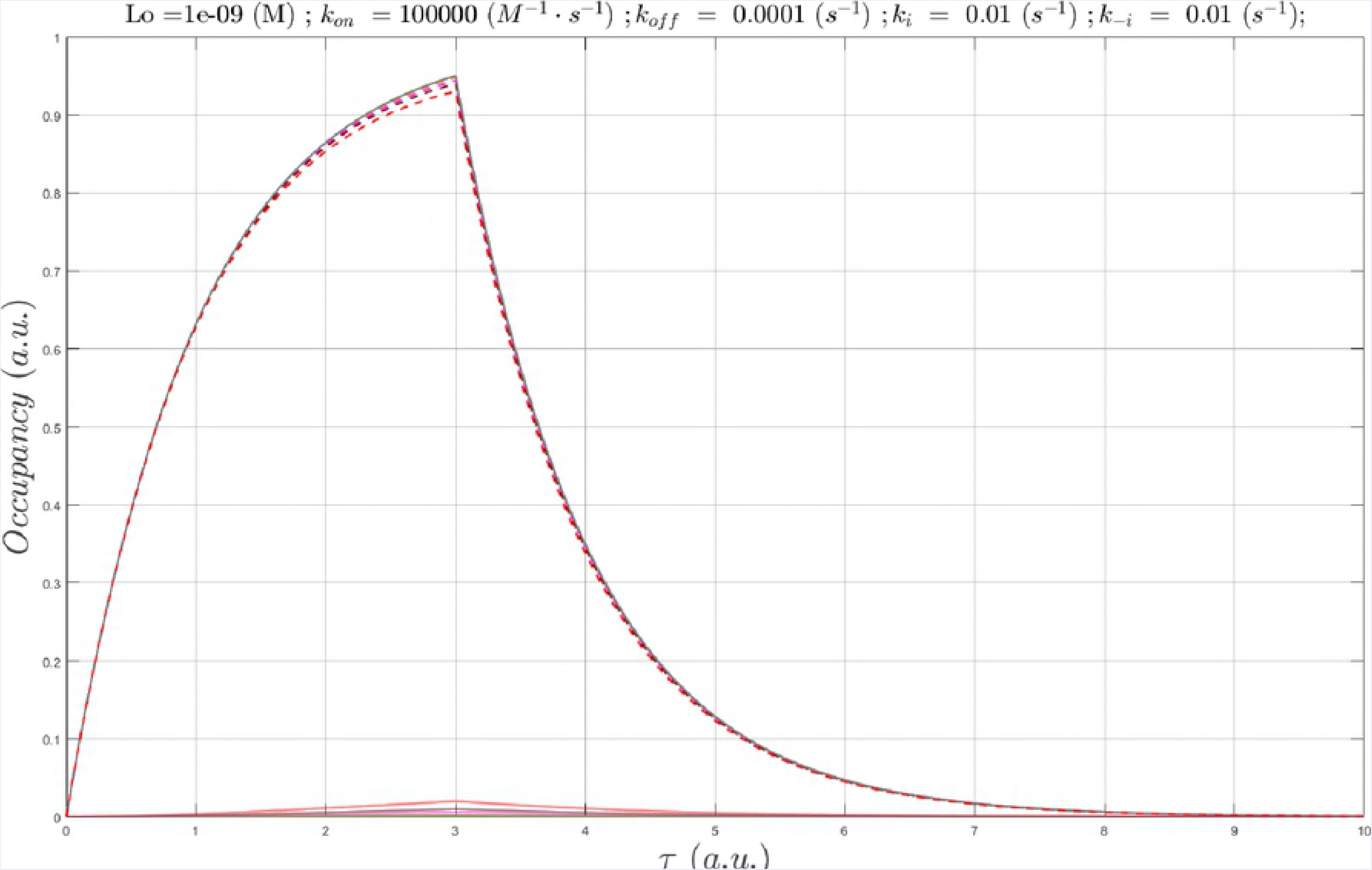
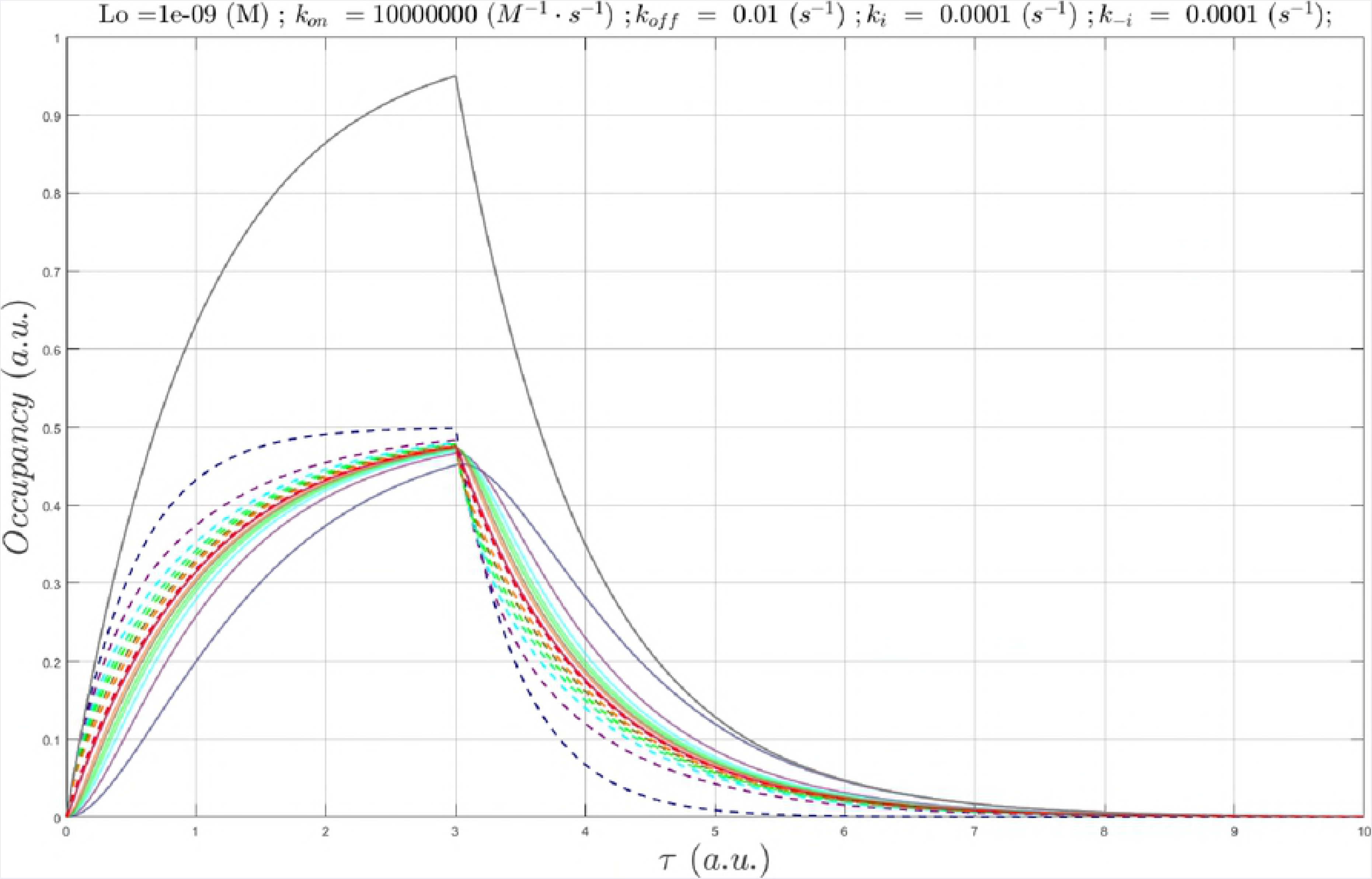
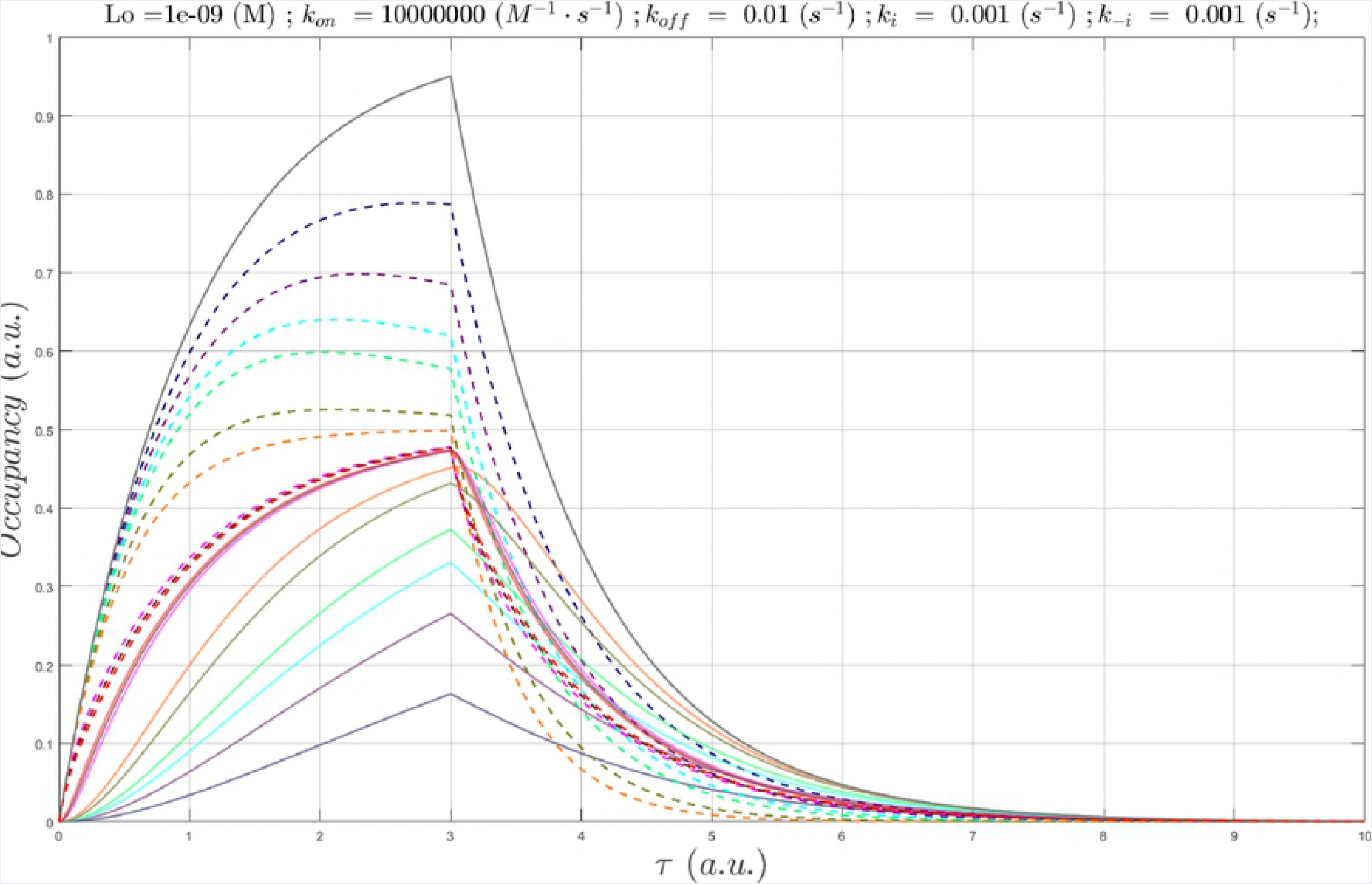
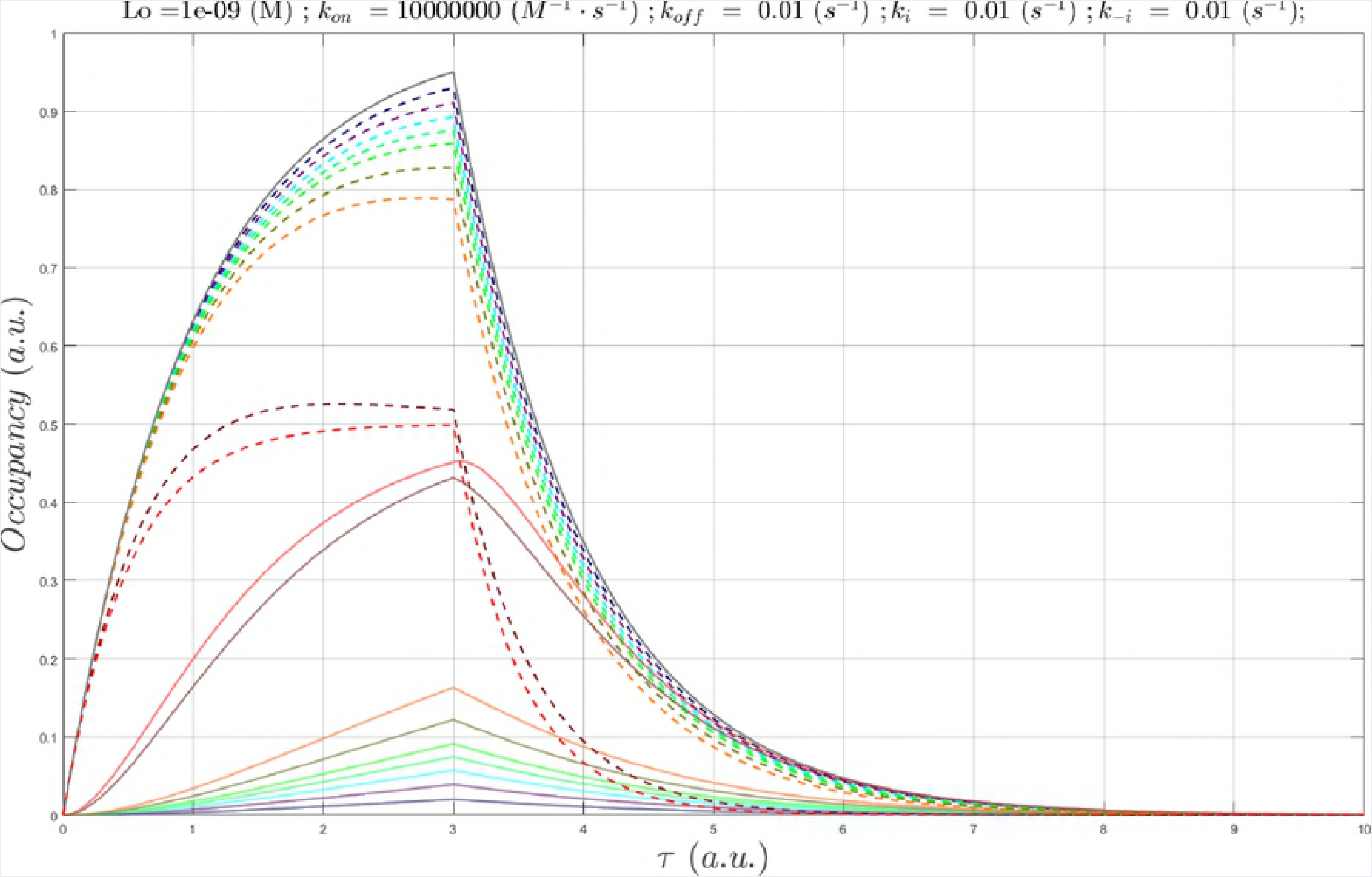
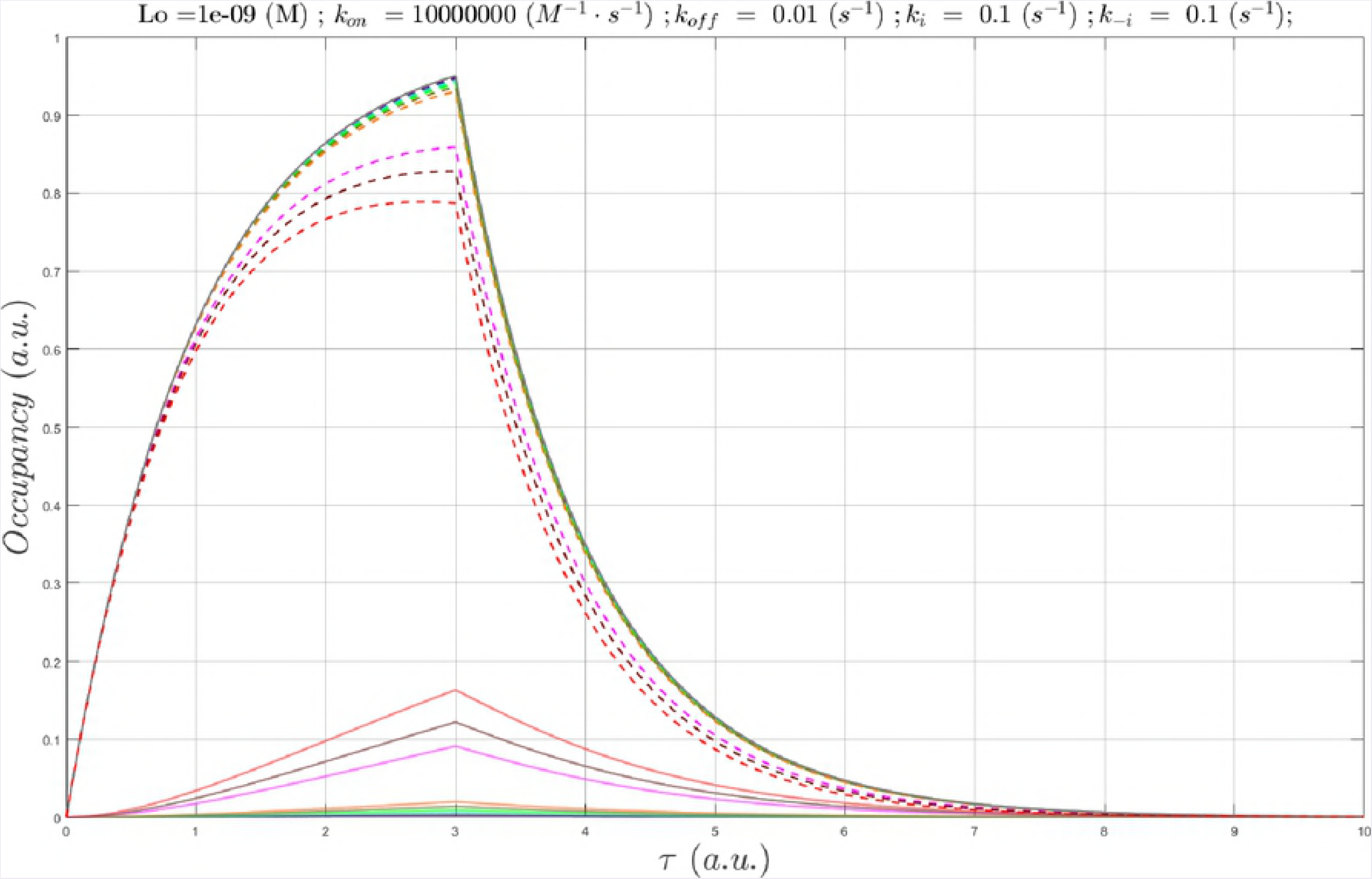
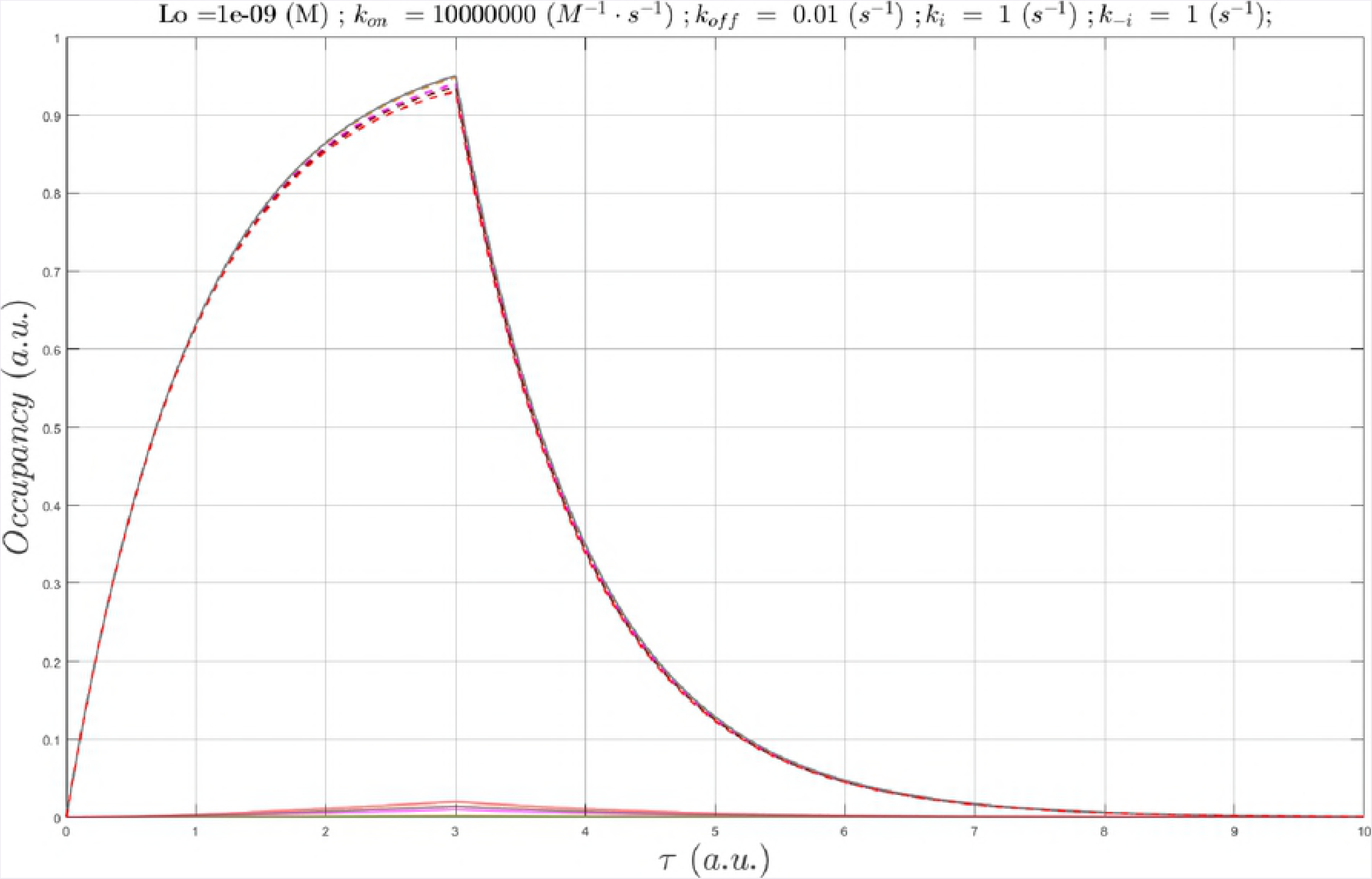
The six branches of biodynamics theory, which span the intra-/peri-cellular (micro-cellular) and macro-cellular levels (see text). Such processes can be simulated using multiscale modeling approaches, ranging from atomistic molecular dynamics-based [1] to “atom-less” time-dependent ODE-based simulations (“MDE mimetics”) [1].

Our theory encompasses the major mechanisms by which:

1. MDEs are generated, powered, solved (integrated), and transduced into function at the micro- and macro-cellular levels.
2. Cellular dysfunction results from molecular dysfunction and alterations in the corresponding MDEs and Γ_a_(t).
3. Cellular dysfunction can be mitigated by exogenously applied MDEs, consisting of drug-target occupancy.

In our previous work, we described the putative mechanisms by which free energy is transduced into non-equilibrium conditions and kinetic barriers (i.e. which we refer to here as “energy dynamics”) [1,4,12–15], and exemplified cellular computing by way of the cardiac AP [4] and a generic MAP kinase-phosphatase system [1]. Here, we focus on binding as a key “integrator” of MDE systems, and in particular, the specific implications of non-equilibrium conditions on occupancy of the bound states of both endogenous molecules and drugs (i.e. the interplay between the “molecular dynamics” and “binding dynamics” branches of our theory).

### Biodynamics both promotes and exploits non-equilibrium conditions

Rapid, transient perturbation-induced responses underlying cellular function depend on open systems, in which mass (chemical precursors, degradation products, and other substances) and energy are exchanged to/from the extracellular volume. Such responses are dampened by reversibility in closed systems exhibiting mass and energy conservation, noting that some species may undergo slow rates of buildup and decay when the overall response rate is slow. In the classical equilibrium view, covalent or non-covalent states i and j are populated according to their free energy differences ({ΔGÿ}), such that the total free energy of the system is minimized. Dynamic state transitions of the participating species necessarily depend on perturbation-re-equilibration under this scenario, which is subject to the following limitations:

1. Perturbation-induced re-equilibration (i.e. equilibrium 1 ➔ equilibrium 2) is time-consuming, and therefore, poorly suited for rapid biological responses.
2. Optimization of complex equilibria is cumbersome and offers extremely limited evolutionary adaptability.

Molecular systems are displaced from equilibrium (analogous to strain) when the participating species are continuously produced and degraded or translocated [16–21] (analogous to stress).

The resulting strain energy is proportional to the stress, analogous to that of a spring stretched away from its equilibrium state. Reversibility (e.g. A + B ⇋ AB) is circumvented by “tensioned” operation, in which △*G* is maintained permanently above zero via chemical or physical entry and exit of the species to/from the system over time (sources and sinks) (Fig 4). The potential energy of a system under such conditions (△*G_noneq_*) is given by:

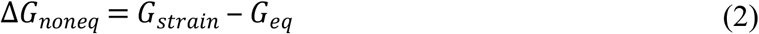

where *G_strain_* is the strain energy, and *G_eq_* is the equilibrium free energy. Strained systems relax via mass action upon release of the corresponding stress, wherein △G_noneq_➔ 0, and the participating species transition toward their equilibrium state distributions (i.e. [species A_state i_]/[species A_state j_] = K_eq_). Reversibility is conveyed indirectly through concurrent (but out-of-I phase) buildup and decay processes (e.g. A + B → AB; AB + C → A + B + C), which we refer to as “dynamic counter-balancing” (“Ying-Yang”) (Figs 2 and 5). Examples include kinase-versus phosphatase-mediated phospho-transfer and inward versus outward ion channel currents.

**Fig 4.**
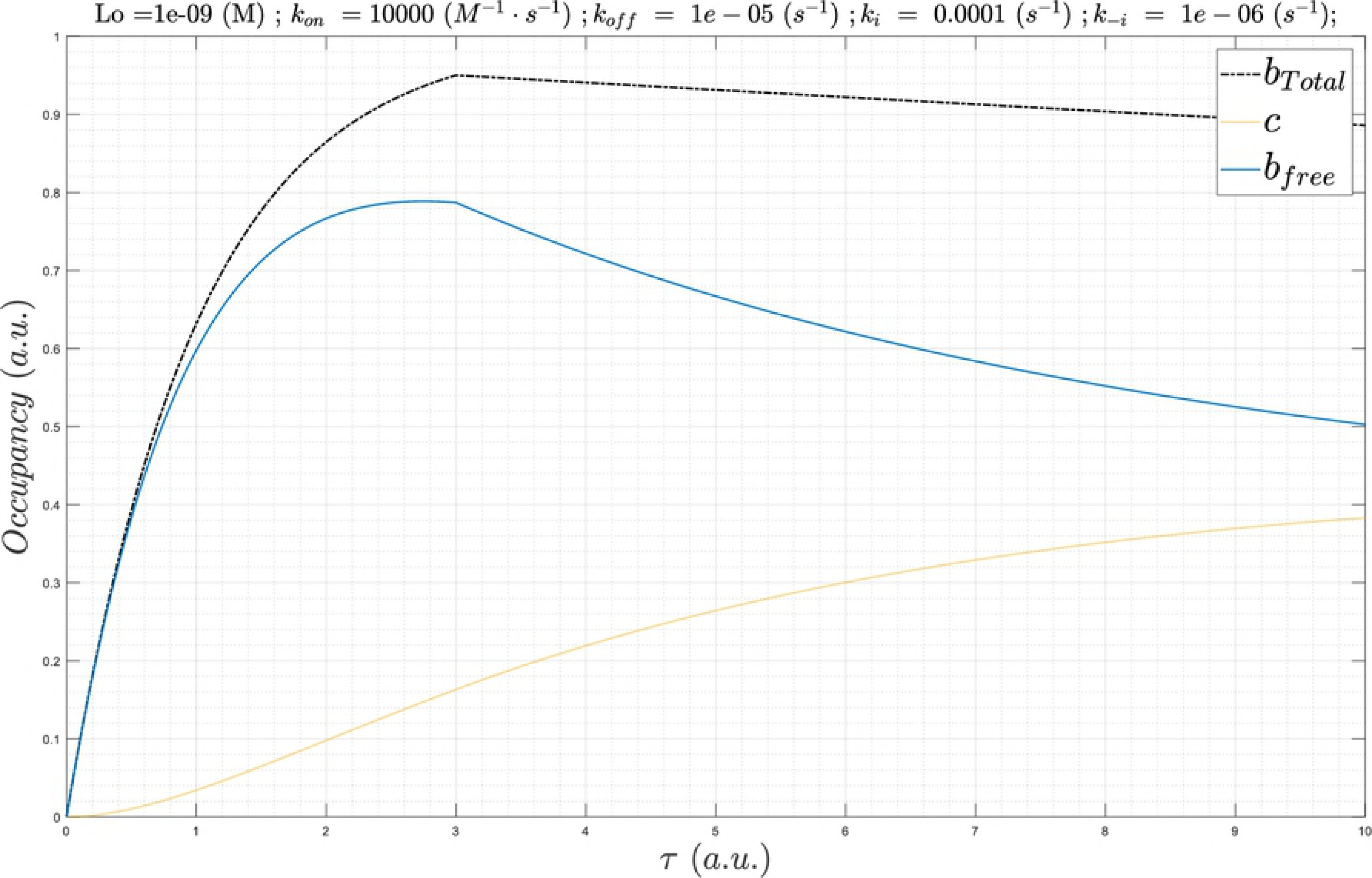
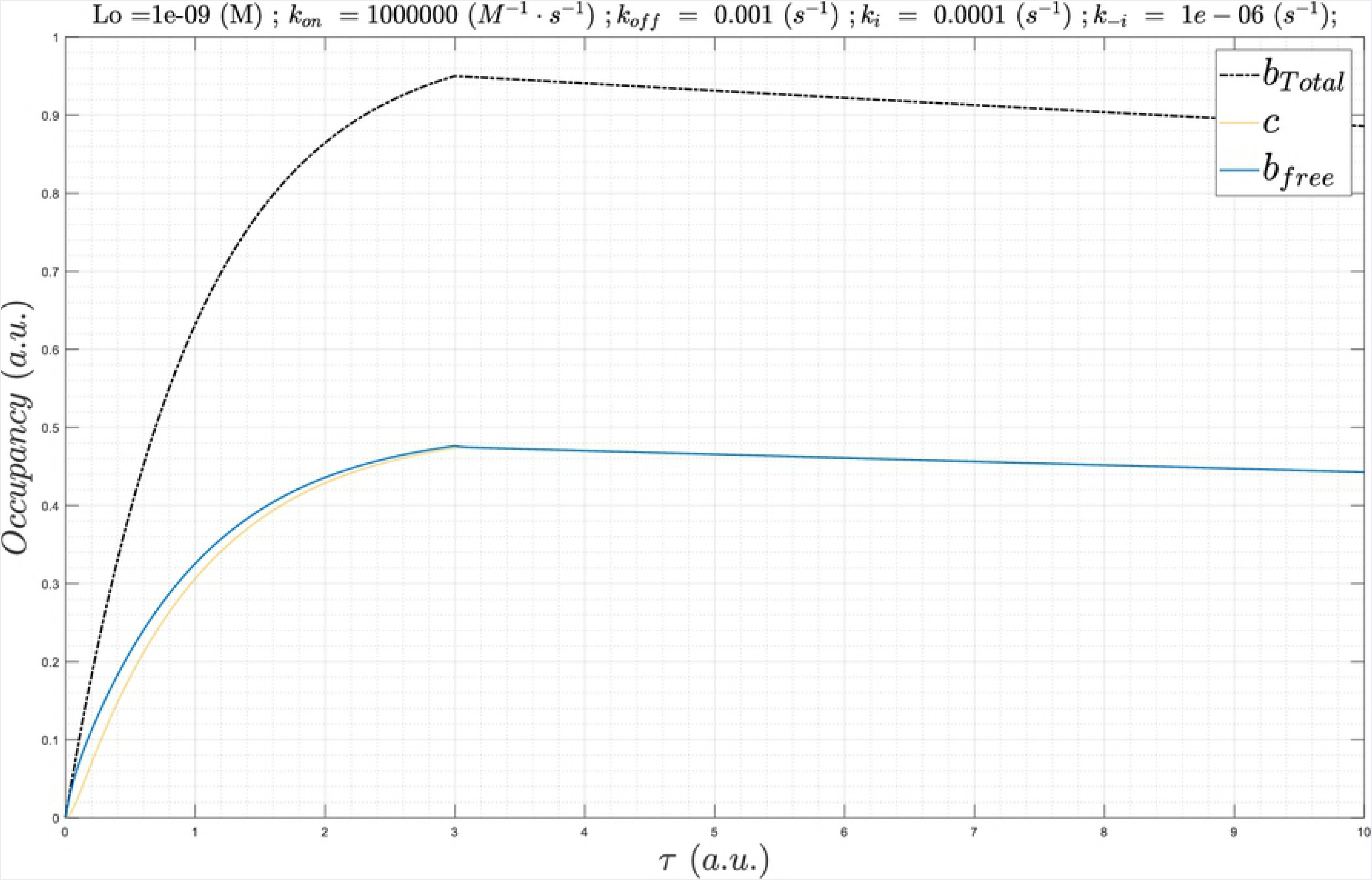
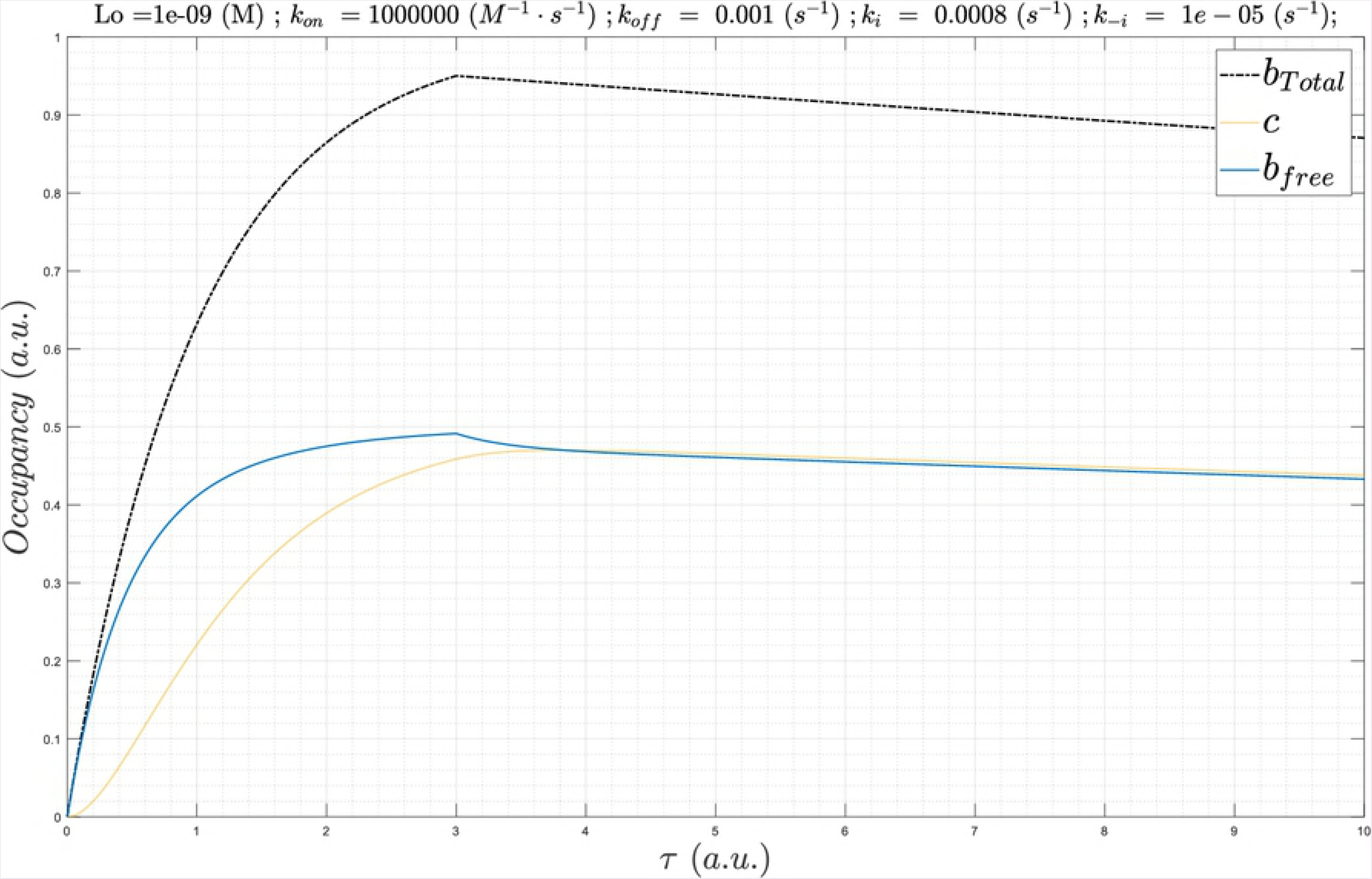
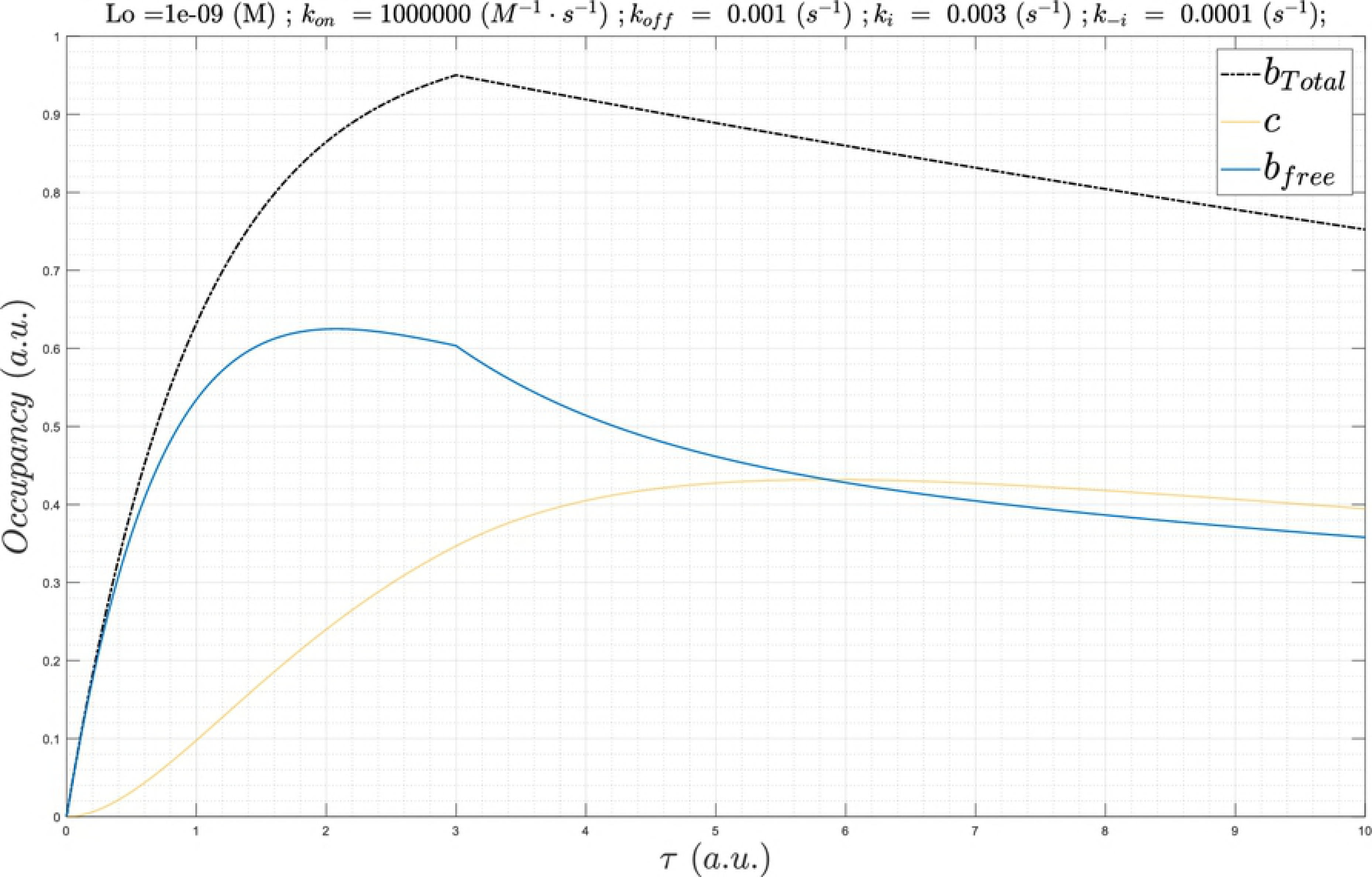
Molecular species and states build and decay <u>transiently</u> (i.e. sources ➔ sinks), thereby generating non-equilibrium conditions. Such conditions promote unidirectional fluxes on which cellular analog computing is based.

**Fig 5.**
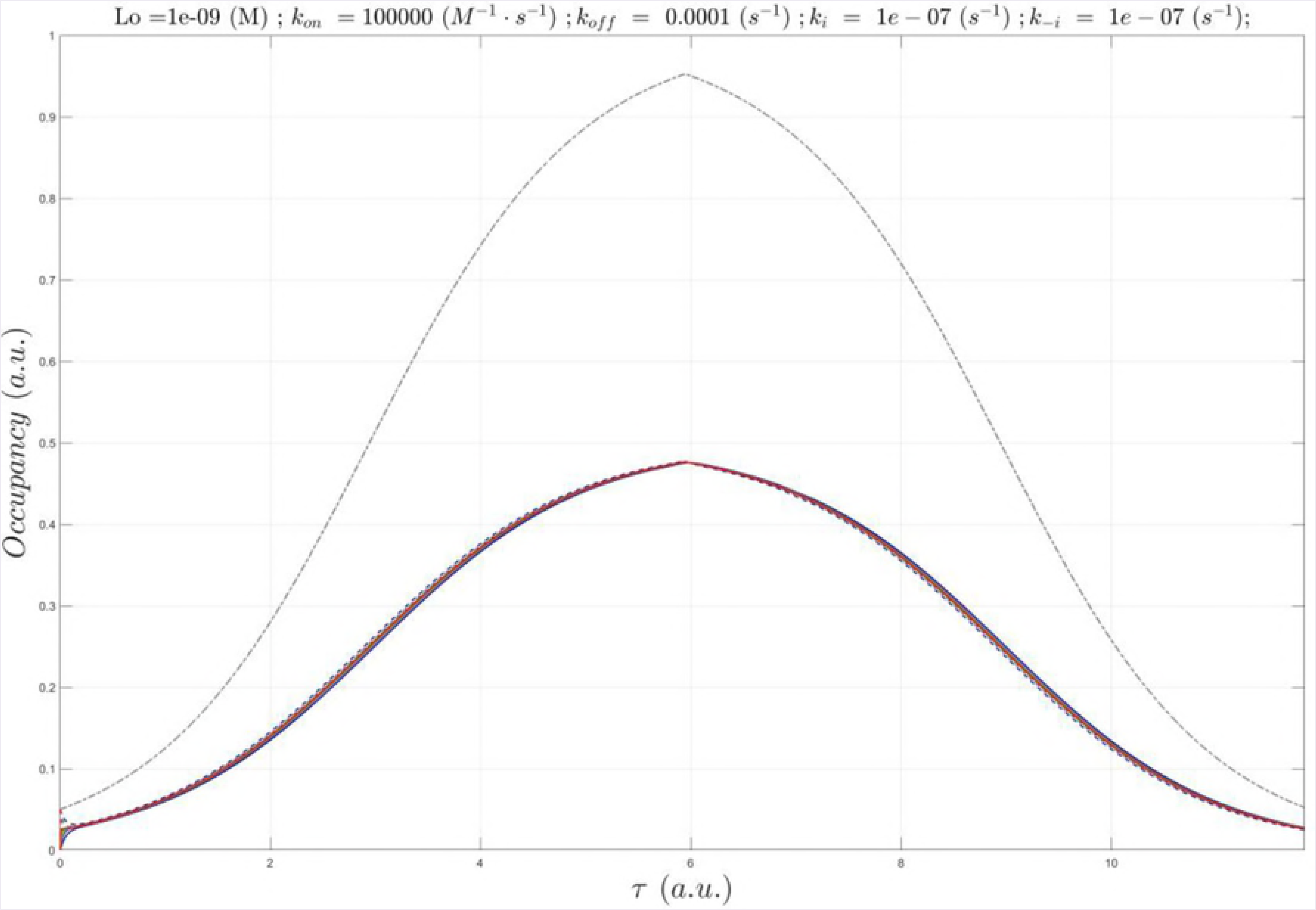
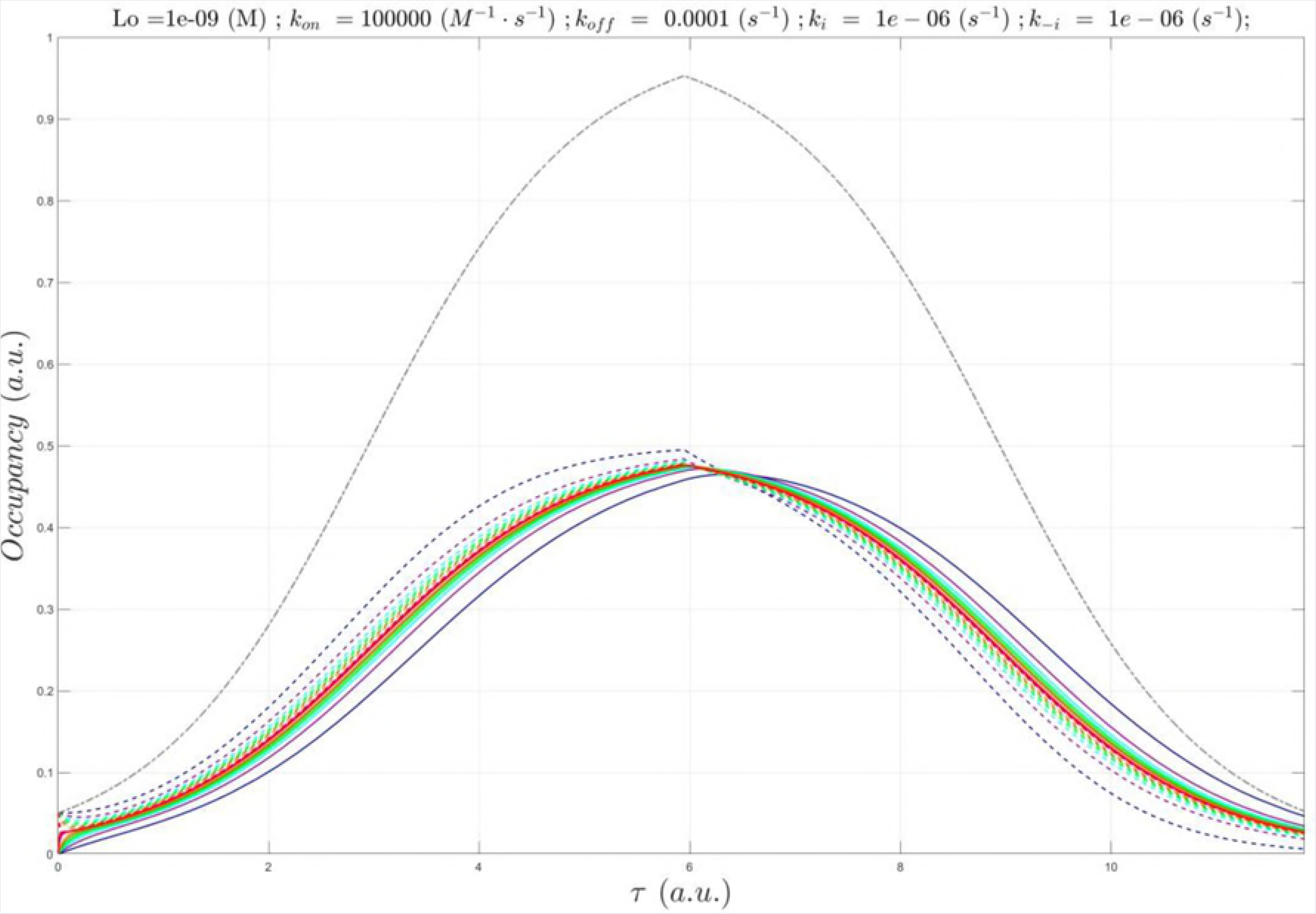
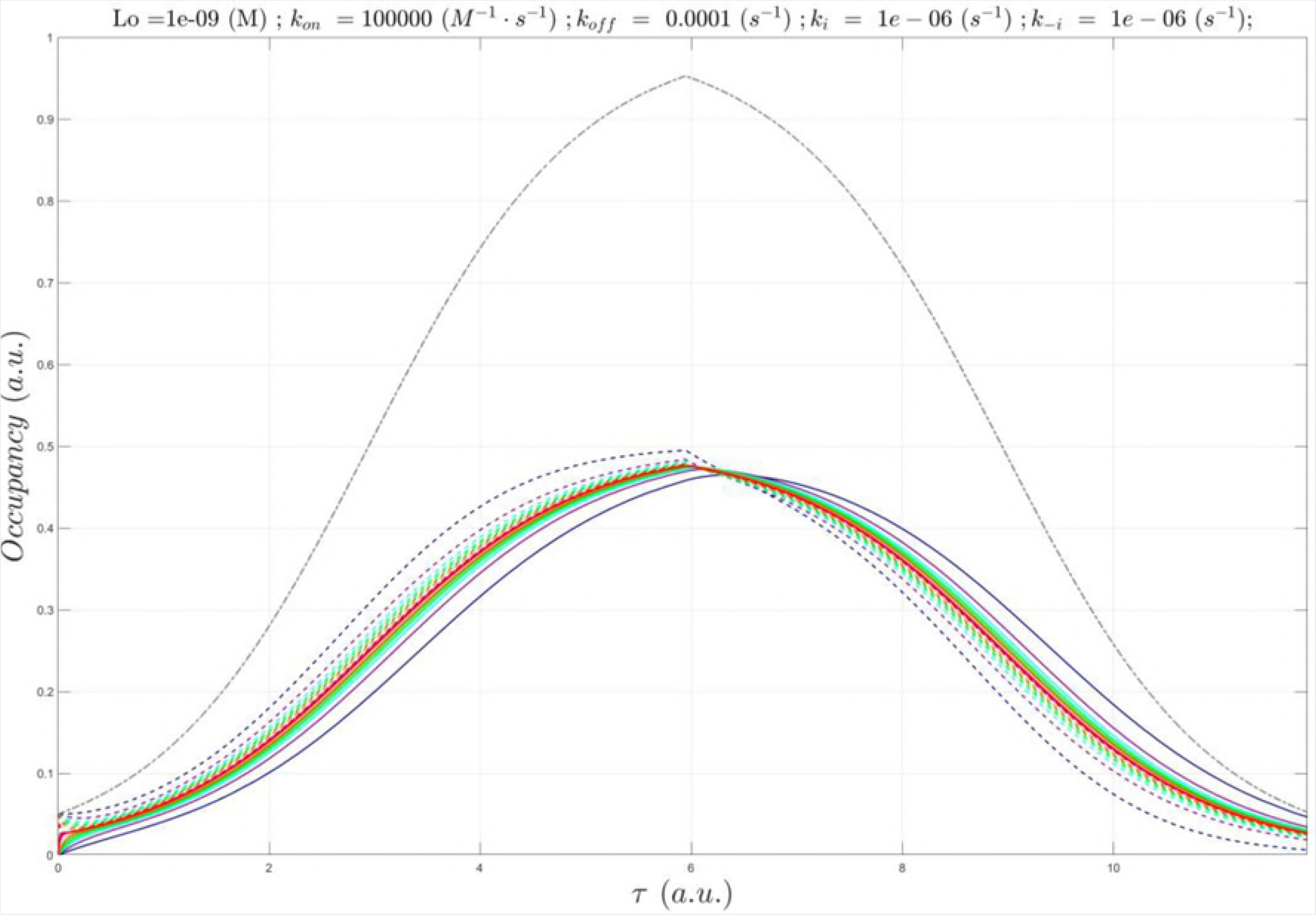
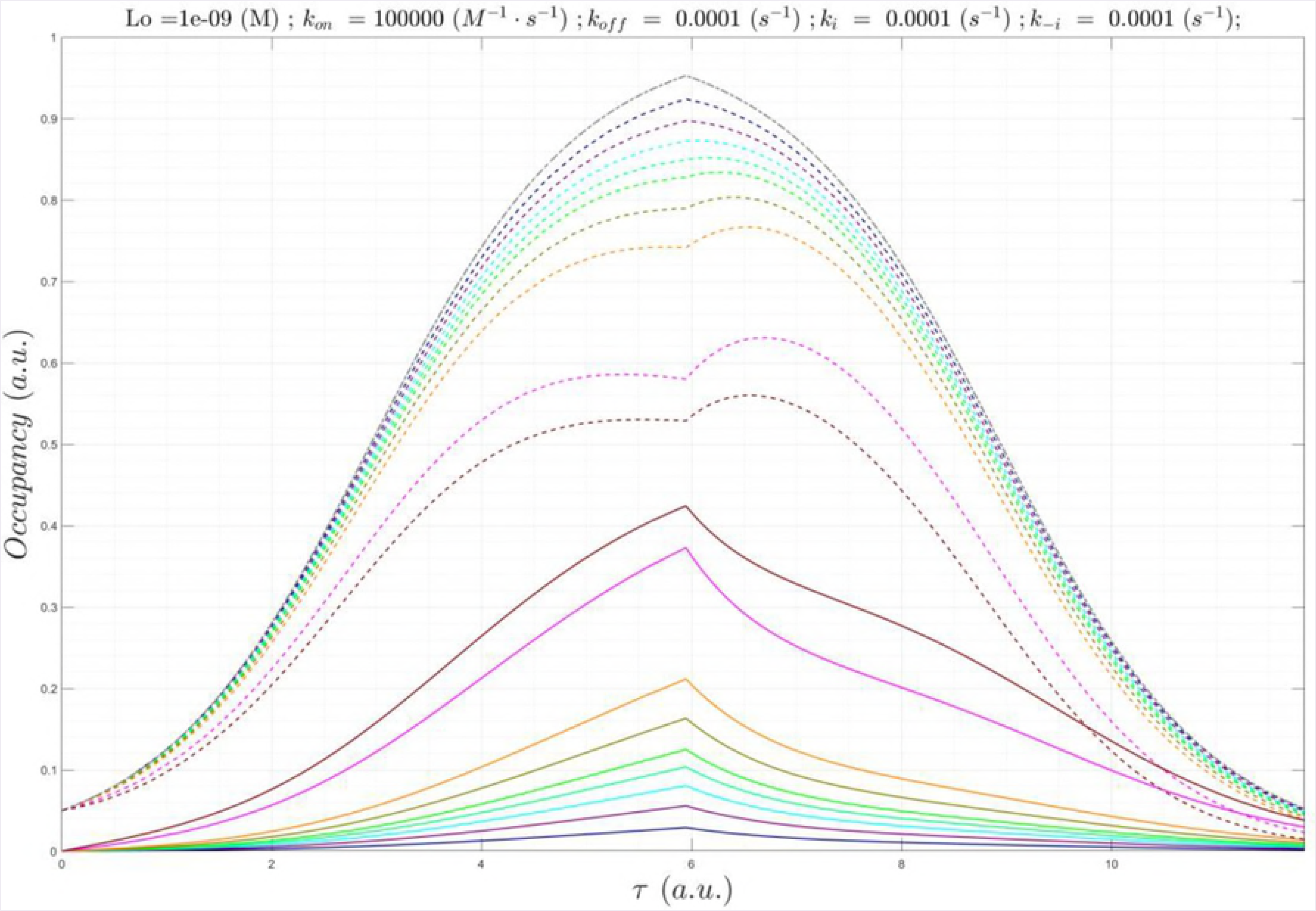
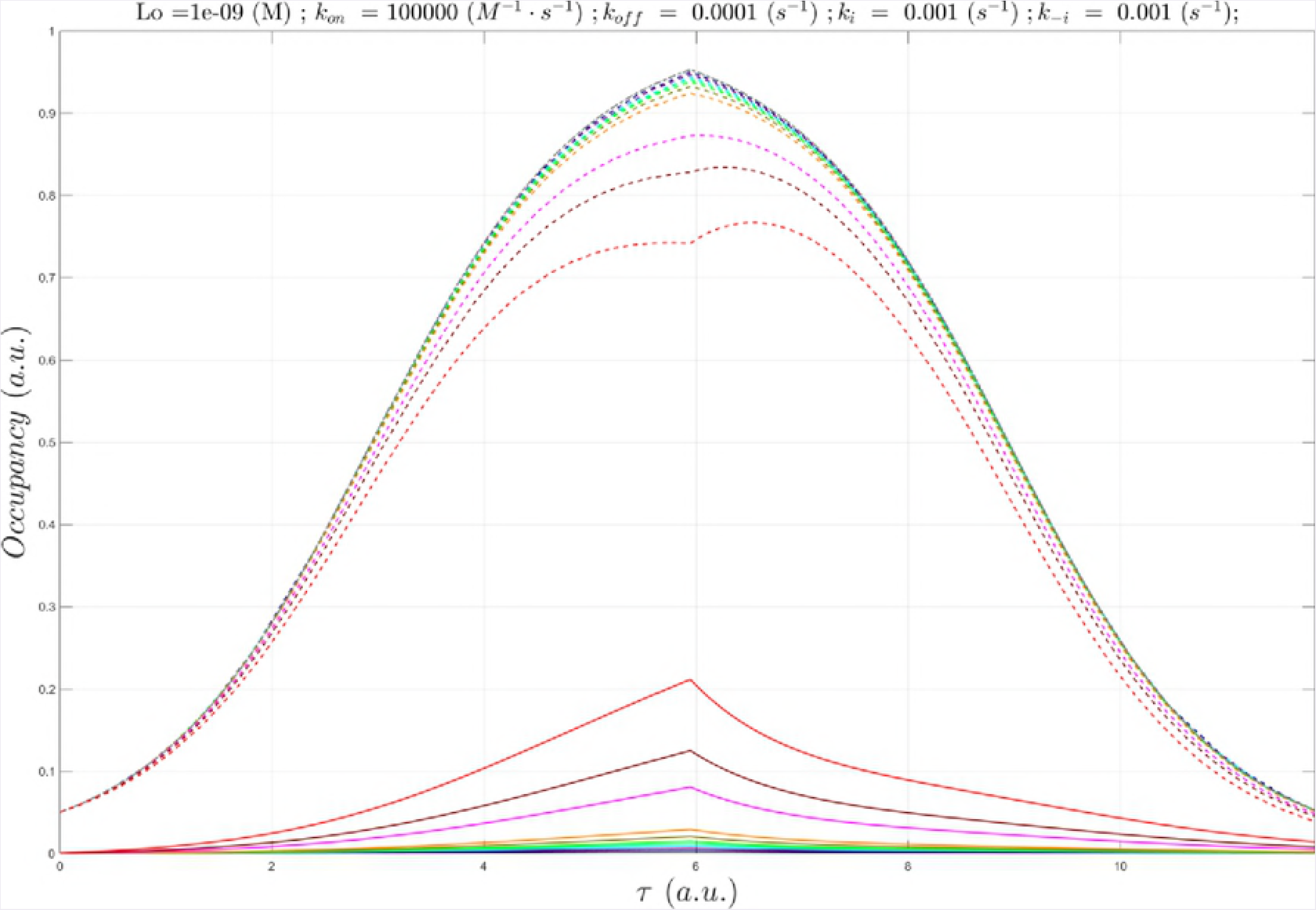
Buildup and decay of a Yin (typically an enzyme), together with its substrate, product (and substrates thereof), Yang across their accessible states, as depicted by a plumbing system consisting of pipes, vessels, and valves. A given function-generating system is constituted from an interdependent collection of m such systems. Each system is maintained at non-equilibrium (analogous to “pressurization”) via continuous inputs and outputs of molecular species (“fluxes”). The dynamic fluid levels within the (n · m) member vessels of the overall system, which define its instantaneous state, are solved by integration of the m MDE collections ({dγ_k_(t)/dt = rate of entry to state k – rate of exit from state k}_i_, where k = 1, 2, 3, …, n, and i = 1, 2, 3, …, m). The vessels may be connected in series or parallel. The flow rates to/from each vessel depend on the input and output valve settings (representing both the intrinsic and extrinsic free energy barriers in this simplified example), which are governed by the overall state of the system (constituting a feedback loop). (B) Dynamic counter-balancing prevents runaway exponential growth and decay of molecular species and state levels. The rates of buildup and decay consist of the net difference in Yin and Yang rates over time. The Yin-Yang balance is tipped toward the Yin during the buildup phase (left side of the blue curve) and toward the Yang during the decay phase (right side of the blue curve). The absolute rates of species or state level changes are irrelevant, except in cases where runaway behavior is functionally desirable (e.g. caspase activation). The overall buildup-decay cycle length is denoted as Λ, noting that long lived species or states likely result from slow decay rather than slow buildup (which would otherwise dampen the maximum level). Fast buildup has important implications for bound state occupancy, which are discussed later in the text.

**Fig 6.**
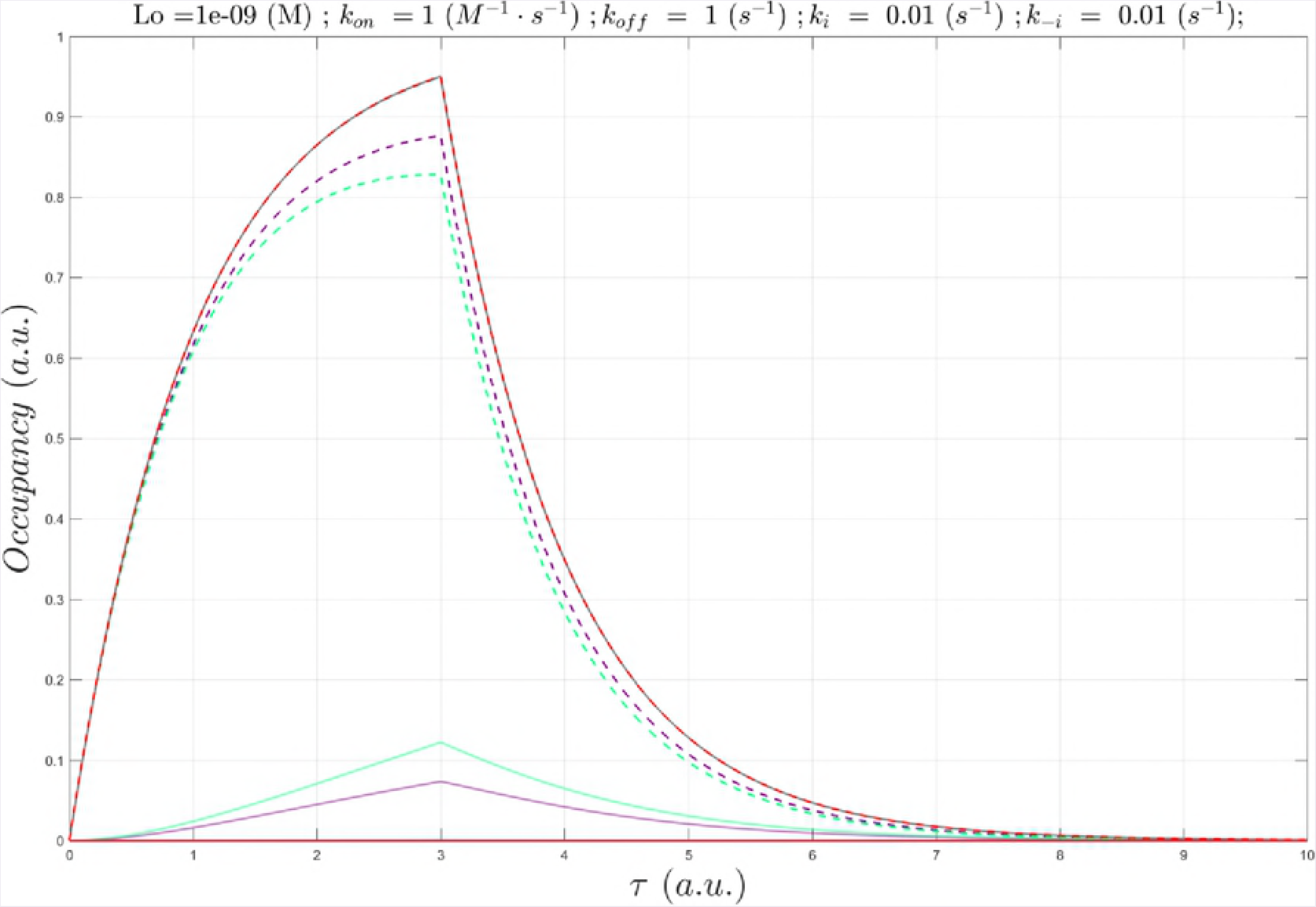
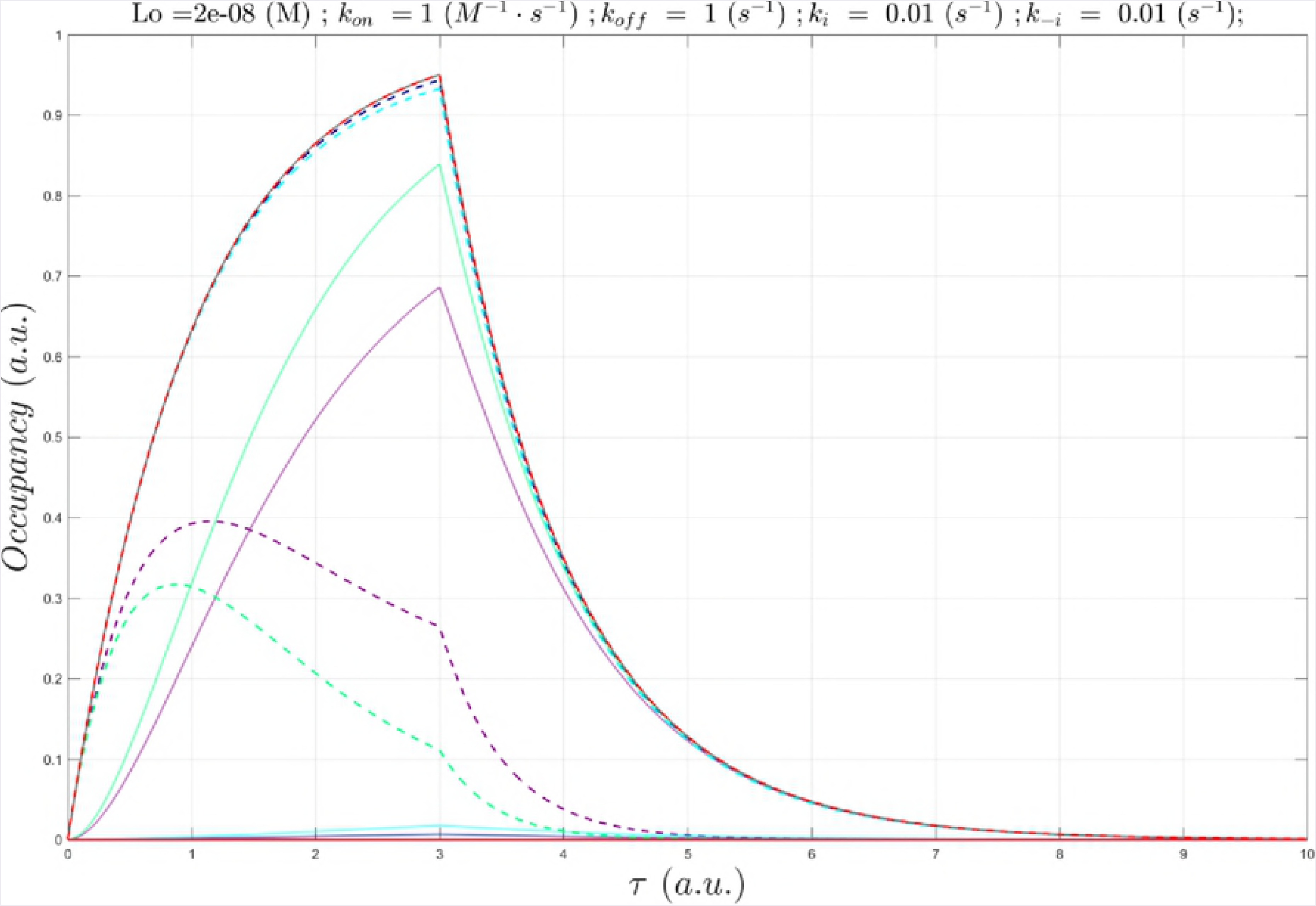
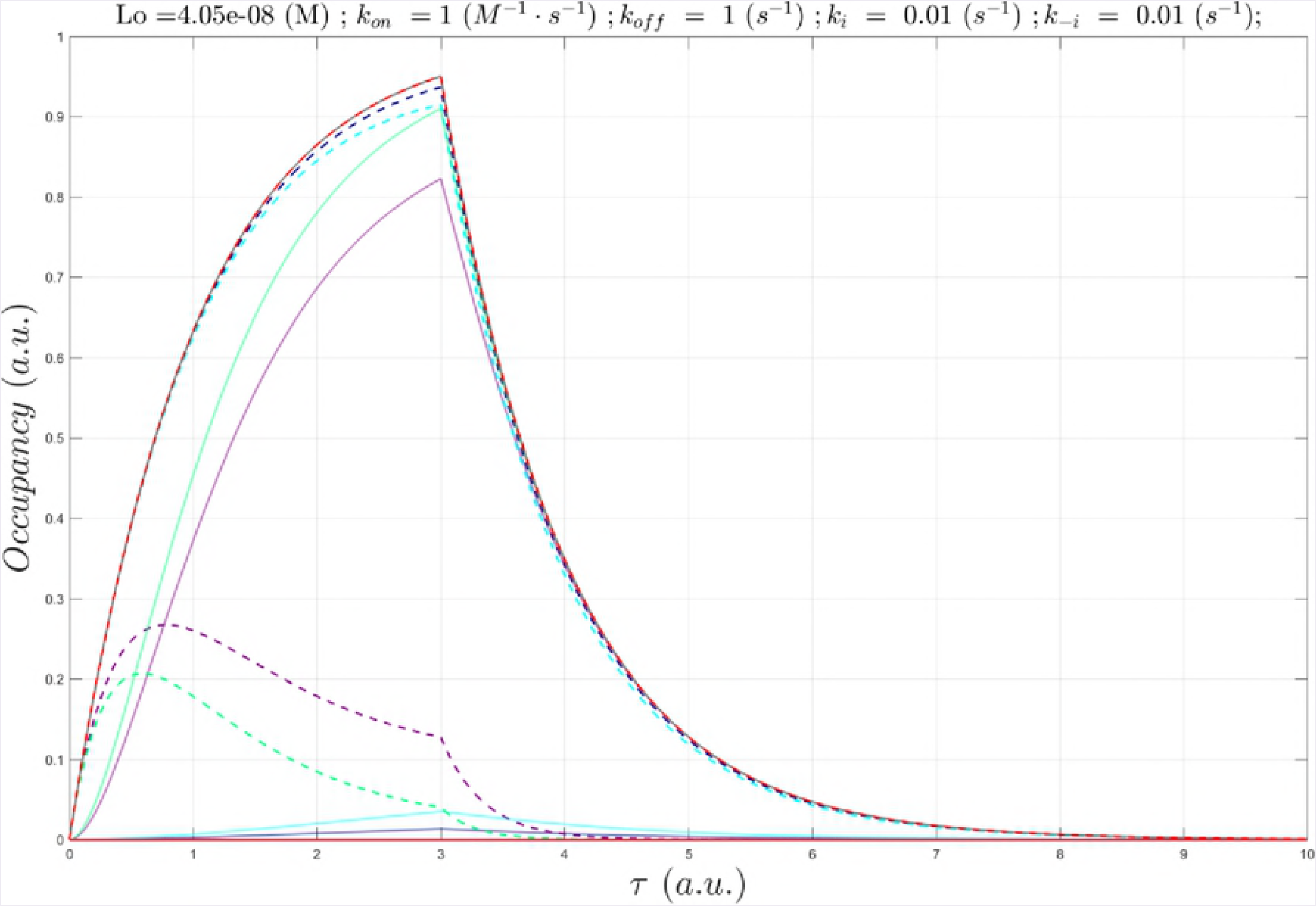
The buildup and decay phases of *B_total_*(*τ*) simulated using equations 6b and 6d (scenario 1). The maximum *B_total_*(*τ*) is normalized to 1.0, and always occurs at the normalized time *τ* = 3. The curve adopts a canonical “saw-tooth” morphology.

**Fig 7.**
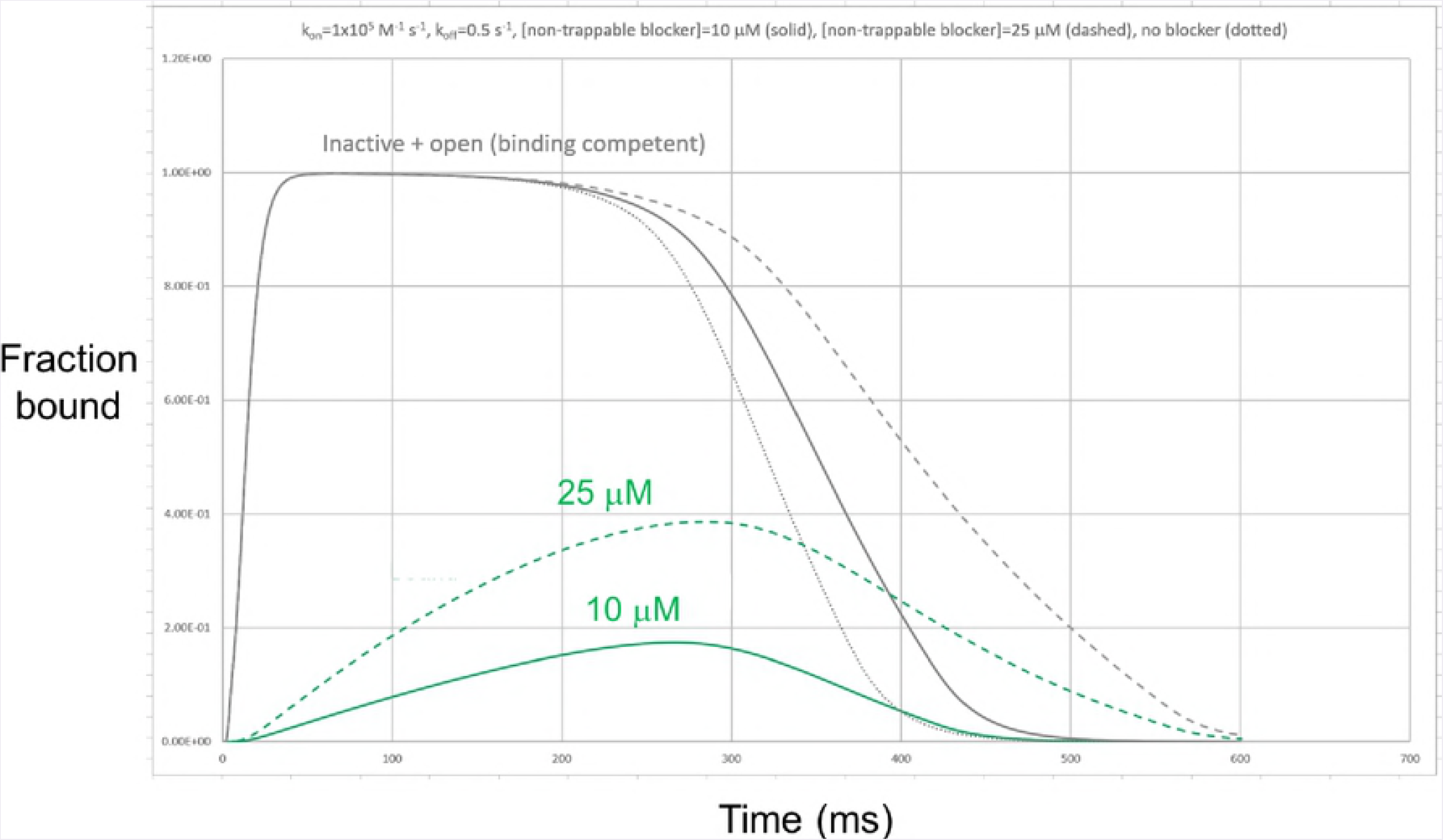
The buildup and decay phases of *B_total_*(*τ*) simulated using equations 6c and 6e (scenario 2). The maximum *B_total_*(*τ*) is normalized to 1.0, and always occurs at *τ* = 6. The morphology approximately resembles that of the state transition curves in Figs 1B and 2A, keeping in mind that the plot is normalized in all cases to a uniform width of *τ* = 12 at all Λ.

The Yin-Yang architecture serves to:

1. Prevent over- and undershooting desired γ_k_(t) due to the <u>exponential behavior</u> of molecular processes (noting that the integral solutions of MDEs are exponential functions). Yins and Yangs are necessarily maintained in an out-of-phase relationship by way of “clocking” (e.g. a phosphatase whose activation depends on a species whose buildup lags behind its target kinase).
2. Restore the initial conditions of the system.

The widespread practice of measuring biochemical (and even cellular and *in vivo*) effects as a function of free ligand concentration, rather than occupancy per se, reflects the extent to which the equilibrium assumption is relied upon throughout the biological sciences. However, our results suggest that time-independent equilibrium metrics of state occupancy are poor approximations to actual occupancy under cellular conditions *in vivo* when the rates of binding partner buildup and decay are fast.

## MATERIALS AND METHODS

We derived two complementary analytical expressions capturing the simultaneous exponential buildup and decay of the binding site (molecular dynamics) and bound state (binding dynamics) based on a simplified second order non-competitive interaction scheme:

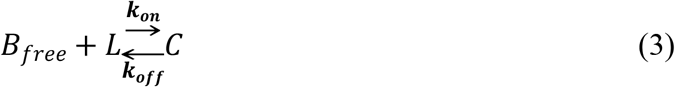

where *k_on_* is the association rate constant, *k_off_* is the dissociation rate constant, and *B_free_*, *C*, and *L* are the free binding site, free ligand, and bound state concentrations, respectively. The rate of change of *C* (the mathematical equivalent of a binding MDE) is given by:

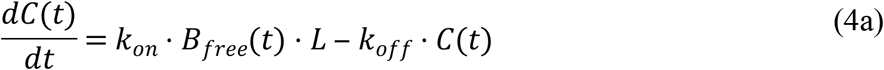

Solution of this equation is facilitated by the following assumptions:

1. Molecular response is proportional solely to dynamic fractional occupancy, which in some cases, may depend additionally on the binding mode (e.g. agonist, partial agonist, inverse agonist, and antagonist in the case of receptors).
2. The total binding site level (*B_total_*(*t*)) is conserved instantaneously, wherein:

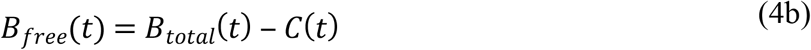

Substituting *Bf_ree_(t*) in equation 4a with equation 4b leads to:

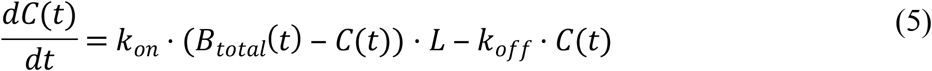
3. That cellular systems operate in the transient regime, in which state population levels build and decay cyclically over time (although quasi-equilibrium operation is possible in some cases).
4. Buildup of *B_total_*(*t*) consists of the separate synthesis of the biomolecule (typically a protein) and generation of its binding-competent structural state. We further assume that state transitions occur on a timescale ≥ the rate of synthesis of the binding site-containing species.
5. Both free and ligand-bound binding sites are lost during degradation, whereas, in practice, different rates of degradation of the bound and unbound states are possible.
6. *B_total_*(*t*) builds and decays <u>exponentially</u> via two distinct processes:

A. Production and degradation of the binding site-containing species, or inter-compartmental transfer to/from the site of action, whichever is slower.
B. Formation and loss of the binding competent structural state.
7. That *B_total_*(*t*) builds to the equilibrium level (*B*_∞_) at t = ∞.
8. That buildup and decay can be treated as separate sequential processes. Although *B_total_*(*t*) builds and decays simultaneously, <u>net</u> buildup occurs prior to *B∞*, followed by <u>net</u> decay. Possible forms of the buildup term include exponential growth (equation 6a), positive exponential decay (equation 6b), or logistic growth (equation 6c). Possible forms of the decay term include exponential (equation 6d) or logistic (equation 6e) decay (multiphasic exponential growth and decay behaviors are also conceivable):

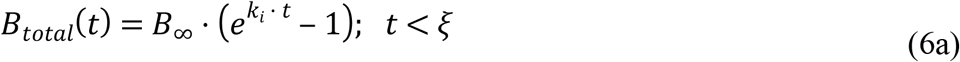

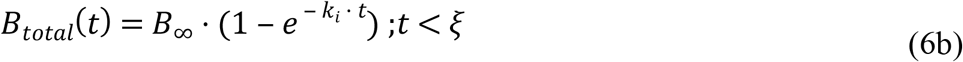

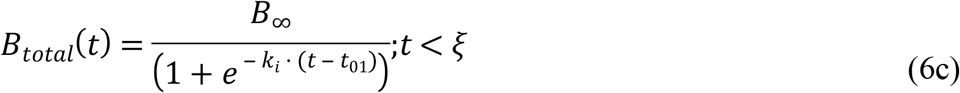

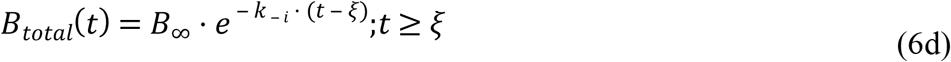

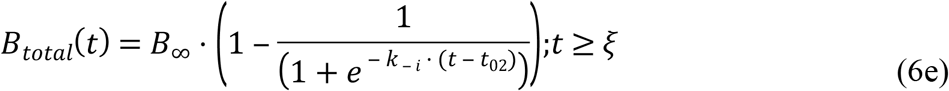

where *B*_∞_ is *B_total_* at t = ∞, k_i_ and k_-i_ are the buildup and decay rate constants, respectively, ξ is a characteristic time at which the function switches from net buildup to net decay, and *t*_01_ and *t*_02_ are the times at which the logistic curves reach 50% of their dynamic range during buildup and decay, respectively. We chose to approximate the buildup and decay phases of the binding site-containing species using equations 6b (based on [22]) and 6d, and the buildup and decay phases of the binding site using equations 6c and 6e, which roughly approximates the state probability curves shown in Figs 1B and 2A.
9. *k_-i_* defines the lower limit of *k_off_* (i.e. *k_off_* is “hijacked” by the rate of binding site decay), necessitating the correction of *k_off_* and *K¿* in cases where *k_off_ < k_-i_*

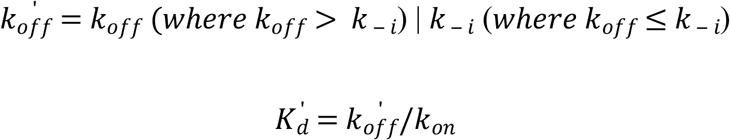
10. *L* is fixed at a constant concentration (*L*_o_). In practice, *L* is expected to build and decay dynamically, which serves to further constrain the buildup of the bound state.

### Scenario 1: putative production and degradation of binding site-containing species

Substituting equations 6b and 6d into equation 5 leads to:

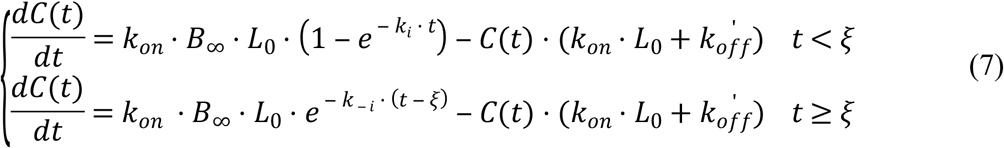

where ξ is the time required to reach equilibrium binding (conventionally referred to as the “settling time”). The net flux switches from buildup to decay at this time. ξ is defined conventionally as the time to reach 0.95 · *B*_∞_ (which we refer to as *B_max_*), given that *B*_∞_ is only asymptotically reached over long time periods [22], whereas *B_max_* is reached over finite times. Solving for *t_settUng_* leads to:

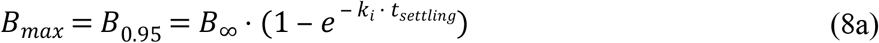

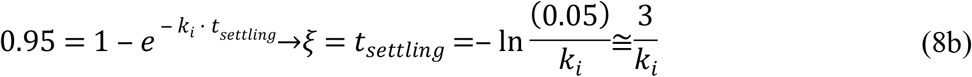

Equation 7 can be recast in dimensionless form to facilitate parameter-independent occupancy comparison, reduce the dimensionality of the system, and allow grouping of the rate terms:

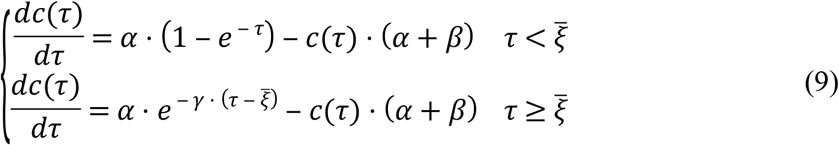

where 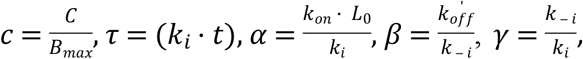, and ξ̄ = 3. Solving equation 9 for *c(τ*) leads to:

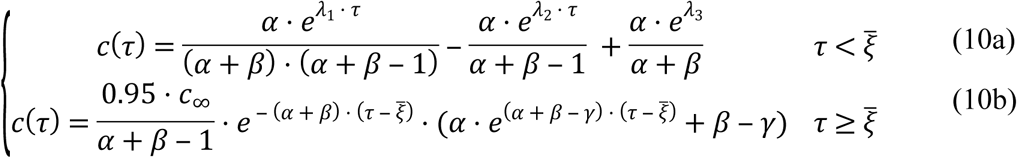

where *λ*_1_ =− (α + β), *λ*_2_ =− 1, and *λ*_3_ = 0. The behavior of equation 10a as a function of *α* and *β* can be described as follows:
***α* + *β* > 0 (satisfied when *α*≥*β*) ➔** stable conditions. *α* = *β* ≥ *k_on_* · *L*_0_ = *k_off_* and cancellation of *k_i_* Under these conditions, equation 10a reduces to:

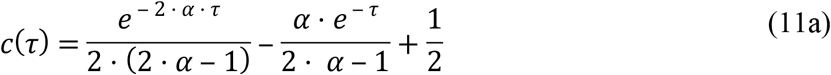

the first term of which undergoes rapid decay at *α*≳ 10 (translating to *k_onSS_* = *k_on_* ≳ 10 · *k_i_*/ *L*_0_, where 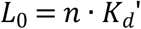), resulting in:

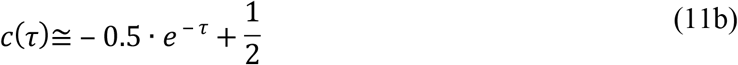

<u>Constant</u> fractional occupancy exists at all time points of equation 11b, which we define as the steady-state occupancy (SSO) profile. Conversely, <u>variable</u> time-dependent fractional occupancy occurs at *α* < 10, which we define as the non-steady-state occupancy (nSSO) profile. Unlike the equilibrium case, an asymmetric relationship exists between *k_on_* and *L*_0_ in governing dynamic occupancy, which follows from the requirement that *α* = *β* ≳ 10. The nSSO profile prevails at all non-saturating *L_0_* in the absence of *α* = *β* ≳ 10. For example:

1. *α* = *β* =1 at ***L*_0_ = 1×10^-9^ M**, *k_on_* = 1×10^5^ M^-1^ s^-1^, *k_off_* = 1×10^-4^ s^-1^, and *k_i_*; = 1×10^-4^ s^-1^ (noting that only *α* = *β* is satisfied in this case).
2. *α* = 10 and *β* = 1 at ***L*_0_ = 1×10^-8^ M**, *k_on_* = 1×10^5^ M^-1^ s^-1^, *k_off_* = 1×10^-4^ s^-1^, and *k_i_* = 1×10^-4^ s^-1^ (noting that only *α* ≳ 10 is satisfied in this case).

*α* + *β* ≫1 ➔ rapid decay of the first term in equation 10a, which at infinite time, reduces to the equilibrium case:

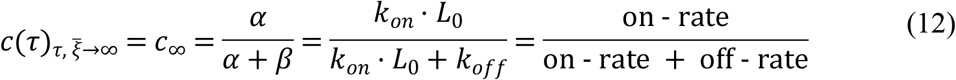

We assume that species or state population levels ≥ 50% of *B_max_* make significant contributions to cellular function, translating to a “functional” buildup + decay time window:

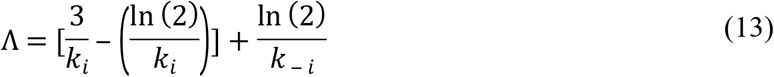

where *k_i_* and *k_-i_* likely range between ms^-1^ (e.g. voltage-gated ion channels) to hr^-1^.

### Scenario 2: putative buildup and decay of the binding competent state

Substituting equations 6c and 6e into equation 5 leads to:

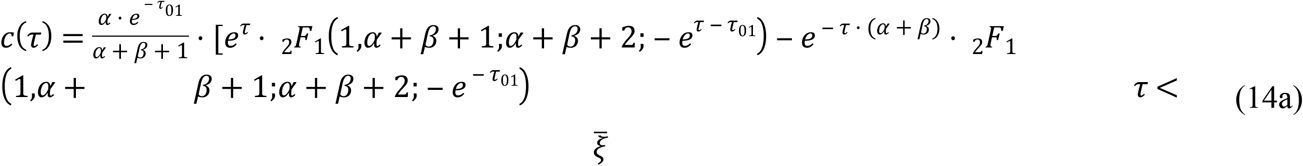

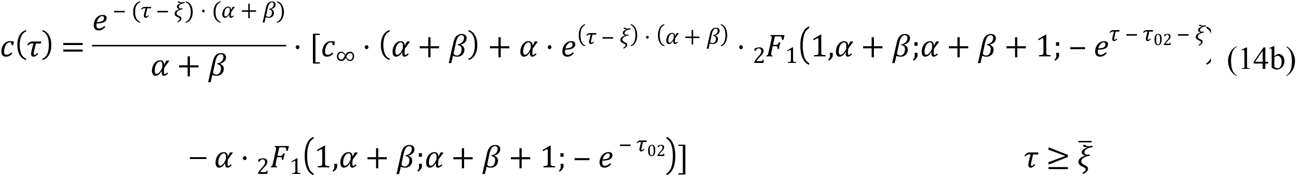

where _2_*F*_1_(*a,b;c;x*) denotes the hypergeometric function. Unlike equation 6b, equation 6c contains a lower asymptote, which approaches zero along the negative time axis. We set *B_total_*(*τ* = 0) to 5% of its full maximum value, and then calculated *τ*_01_, *τ*_02_, and 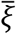 (the normalized settling time corresponding to *B_max_* =95% of the final value), as follows:

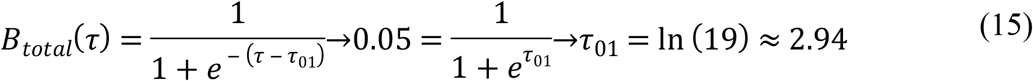

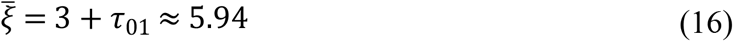

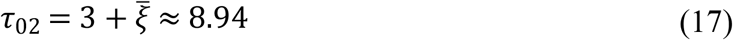

We solved equation 14 both analytically and numerically using Wolfram Alpha (Wolfram Research, Champaign, IL, 2017) and MATLAB (Version 9.2a, The MathWorks, Inc., Natick, MA 2017), respectively. As for scenario 1, we assume that species levels or state populations ≥ 50% of *B_max_* make significant contributions to cellular function, which leads to the following expression for Λ:

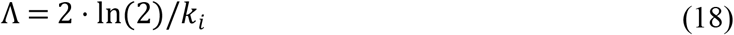

### Comparison of equilibrium versus non-equilibrium occupancy

Static <u>non-competitive</u>, <u>non-cooperative</u> occupancy (γ) under equilibrium conditions can be described by the following form of the Hill equation [23]:

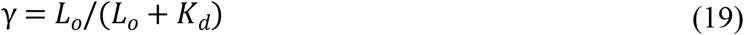

where *L*_0_ = n · *K_d_*. Fractional occupancy under the non-equilibrium SSO profile (*c*(*τ*)) is constant over time, and is also described by equation 19. However, *K_d_* is non-equivalent under equilibrium versus non-equilibrium conditions:

#### Equilibrium

*K_d_* = *k_off_*/*k_0n_* ➔ absolute occupancy = *γ* · *B*_0_, where *B*_0_ is the fixed binding site concentration.

#### Non-equilibrium SSO

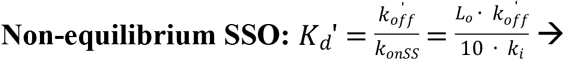 absolute occupancy = *γ* · *B_total_*(*t*). Achieving the SSO profile depends on in-step buildup and decay with, *B_total_*which in turn, depends on *k_on_*≥*K_onSS_*

#### Non-equilibrium nSSO

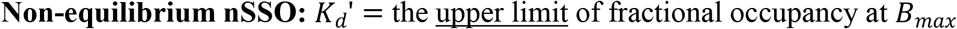. nSSO occurs at all *k_on_*≥*K_onSS_*, irrespective of 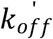 and 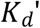 The quasi-steady-state occupancy profile is achieved at near saturating *L*_0_ levels (referred to hereinafter as qSSO).

We assume the most biologically and pharmacologically desirable profile consists of SSO (i.e. where the intrinsic and extrinsic rates are tuned), which gives rise to constant fractional occupancy (*B_free_*(*t*)/*B_total_*(*t*)) over time. However, the other profiles may suffice in cases where cellular function or pharmacological/toxic effects are driven by peak, rather than sustained occupancy (e.g. ion channel blockade vis-à-vis pro-arrhythmia [4]), which necessarily coincides with *C_max_*.

### Characterization of dynamic binding site occupancy under conditions of time-dependent binding site availability

We used equations 10 and 14 to explore the effect of decreasing Λ (and, in particular, speeding *k_i_*) on achieving the SSO versus nSSO profile as a function of:

**Λ**: 83,000 hr (equilibrium) to 300 ms (the approximate timescale of ion channel transitions), sampled as noted.

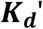 (for reference): fixed at 1 nM, except where otherwise noted.

***k_on_***: 1×10^9^ (diffusion limit) to 1×10^3^ M^-1^ s^-1^, sampled as noted.

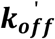. 1×10^0^ to 1×10^-6^ s^-1^, sampled as noted.

*L*_0_: sampled at 1 nM (i.e. 50% occupancy at equilibrium, equating to 0.95 · 50% in our model), 19 nM (95% occupancy at equilibrium, equating to 0.95 · 95% in our model), as well as other concentrations (as noted).

### Key limitations of our approach

Our generalized analytical occupancy relationships avoid assumptions about the specific dynamic mechanisms driving binding site availability. However, such simplified relationships are subject to certain limitations compared with Markov-based simulation models tailored to specific cellular systems (e.g. as was used in our previous work [1,4]), including the potential for underestimated sensitivity of dynamic occupancy to decreasing *Λ* resulting from: 1) the use of <u>fixed</u> free ligand concentration (where ligand and binding site buildup may or may not occur in phase); and 2) neglect of competition with endogenous molecules that may also build and decay over time.

## RESULTS

Whereas static occupancy can be quantified using the Hill, Michaelis-Menten, and similar algebraic expressions, quantification of dynamic occupancy necessitates more complex time-dependent differential equation-based models. However, detailed knowledge of the participating species, parameters (e.g. rate constants), and initial conditions (e.g. states and species levels) is needed to fully implement such models. We set about in this work to characterize the relationships between intrinsic and extrinsic rates and SSO versus nSSO profiles using a set of closed form dimensionless relationships that we derived for this purpose (described in MATERIALS AND METHODS).

### Scenario 1

In this scenario, binding site lifetimes are assumed to depend on the rates of production and degradation of binding site-containing species versus formation and decay of the binding-competent state (i.e. transition to the binding competent state fully mirrors the buildup and decay of the species). 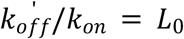 was maintained in all cases, except as otherwise noted. We characterized the sensitivity of *c*(*τ*) to *k_on_* and *k_off_* as a function of decreasing *Λ* based on the following criteria:

1. The non-equilibrium threshold of *Λ* at which occupancy becomes kinetically-versus *K_d^-^_* driven (i.e. the equilibrium regime).
2. The approximate *k_on_* and 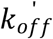 required to achieve 50% and 95% occupancy under SSO conditions (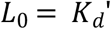 and 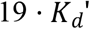 respectively).
3. The fold-increase in *L_0_* required to approach the qSSO profile.

The results are summarized below and in Table 1.

**Table 1.** Approximate *k_onSS_* values needed to achieve the SSO profile as a function of *k_i_*, calculated from 10 · *k_i_*/*L*_0_, where *L_0_* = *K_d_* and *k_off_* ≥ *k_− i_* (noting that *k_off_* is not hijacked under these conditions).

| $\Lambda$ | $k_i$<br>($\text{s}^{-1}$) | *Buildup*<br>$t_{1/2}$ | $k_{-i}$<br>($\text{s}^{-1}$) | $K_d = k_{off}$<br>/$k_{on}$<br>($\text{nM}$) | $k_{off} =$<br>$K_d \cdot k_{on}$<br>($\text{s}^{-1}$) | $\sim k_{onSS}$<br>($\text{M}^{-1} \text{ s}^{-1}$) |
| --- | --- | --- | --- | --- | --- | --- |
| 83,333 hr | $1 \times 10^{-8}$ | 19,254 hr | $1 \times 10^{-8}$ | 1 | - | - |
| 833 hr | $1 \times 10^{-6}$ | 192.5 hr | $1 \times 10^{-6}$ | 1 | $1 \times 10^{-5}$ | $1 \times 10^4$ |
| 83 | $1 \times 10^{-5}$ | 19.25 hr | $1 \times 10^{-5}$ | 1 | $1 \times 10^{-4}$ | $1 \times 10^5$ |
| 8.33 hr | $1 \times 10^{-4}$ | 1.9 hr | $1 \times 10^{-4}$ | 1 | $1 \times 10^{-3}$ | $1 \times 10^6$ |
| 50 min | $1 \times 10^{-3}$ | 11.6 min | $1 \times 10^{-3}$ | 1 | $1 \times 10^{-2}$ | $1 \times 10^7$ |
| 5 min | $1 \times 10^{-2}$ | 1.16 min | $1 \times 10^{-2}$ | 1 | $1 \times 10^{-1}$ | $1 \times 10^8$ |
| 30 s | $1 \times 10^{-1}$ | 6.9 s | $1 \times 10^{-1}$ | 1 | $1 \times 10^0$ | $1 \times 10^{9\$}$ |
| 3 s | $1 \times 10^0$ | 0.69 s | $1 \times 10^0$ | 1 | $1 \times 10^1$ | $1 \times 10^{10\$}$ |
| 300 ms | $1 \times 10^1$ | 69 ms | $1 \times 10^1$ | 1 | $1 \times 10^2$ | $1 \times 10^{11\$}$ |
| 199 hr | $1 \times 10^{-4}$ | 1.9 hr | $1 \times 10^{-6}$ | 1 | $1 \times 10^{-3}$ | $1 \times 10^6$ |
| 20 hr | $8 \times 10^{-4}$ | 14.4 min | $1 \times 10^{-5}$ | 1 | $1 \times 10^{-2}$ | $1 \times 10^7$ |
| 2.1 hr | $3 \times 10^{-3}$ | 3.9 min | $1 \times 10^{-4}$ | 1 | $1 \times 10^{-1}$ | $1 \times 10^8$ |

§Exceeds the diffusion limit (further gains in $c(\tau)$ are necessarily driven by $L_0$ ).

#### The equilibrium regime

The equilibrium regime extends from Λ > 83,333 hr (*k_i_* = *k_− i_* = 1×10^-8^ s^-1^), at which *c*(*τ*) is fully independent of absolute *k_off_* and *k_on_* (constrained in our study to *k_off_*/*k_on_* = *K_d_*), converging in all cases to the SSO profile (Fig 8). Under these conditions, the bound fraction is 50% or 95% (*L_0_* = 1 nM or 19 nM, respectively) at all instants of time, and *c*(*τ*) exhibits the signature “saw-tooth” morphology of *B_total_*(*τ*). The quasi-equilibrium regime resides between 833 < Λ < 83,333 hr (Fig 9A).

**Fig 8.**
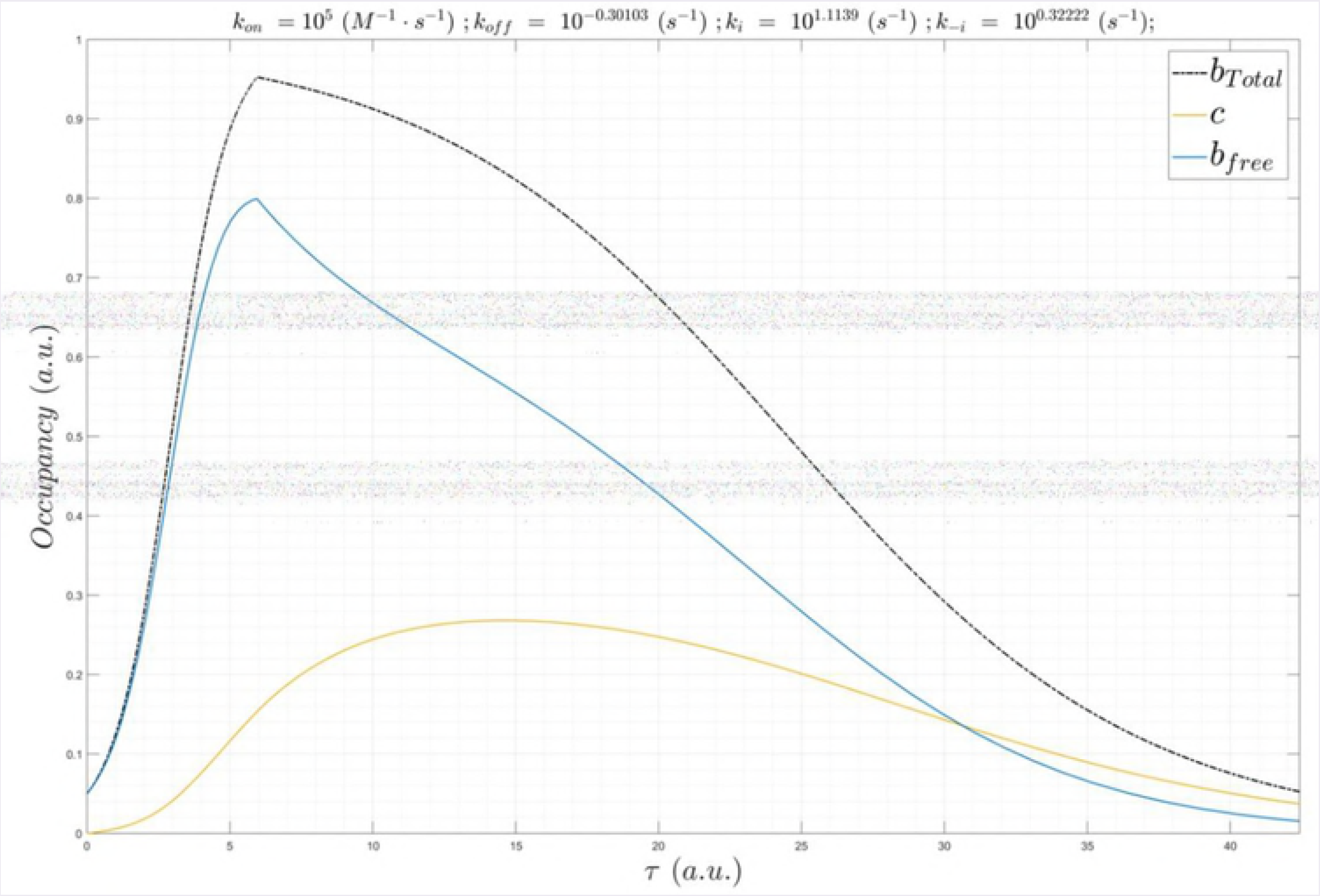
(A) Plot of *B_total_*(*τ*) (gray), *c*(*τ*) (red), and *B_free_*(*τ*) (obscured by *c*(*τ*)). *k_i_* = *k _− i_* = 1×10^-8^ s^-1^, Λ = 83,333 hr, and *L_0_* = *K_d_* = 1 nM, which yields 50% SSO at all times (denoted by equal lengths of the vertical arrows above and below their respective centroids), irrespective of *k_on_* and *k_off_* (holding *K_d_* constant at 1 nM). (B) Same as A, except with *L*_0_ = 19 · *K_d_*, which yields the expected SSO = 95%.

**Fig 9.**
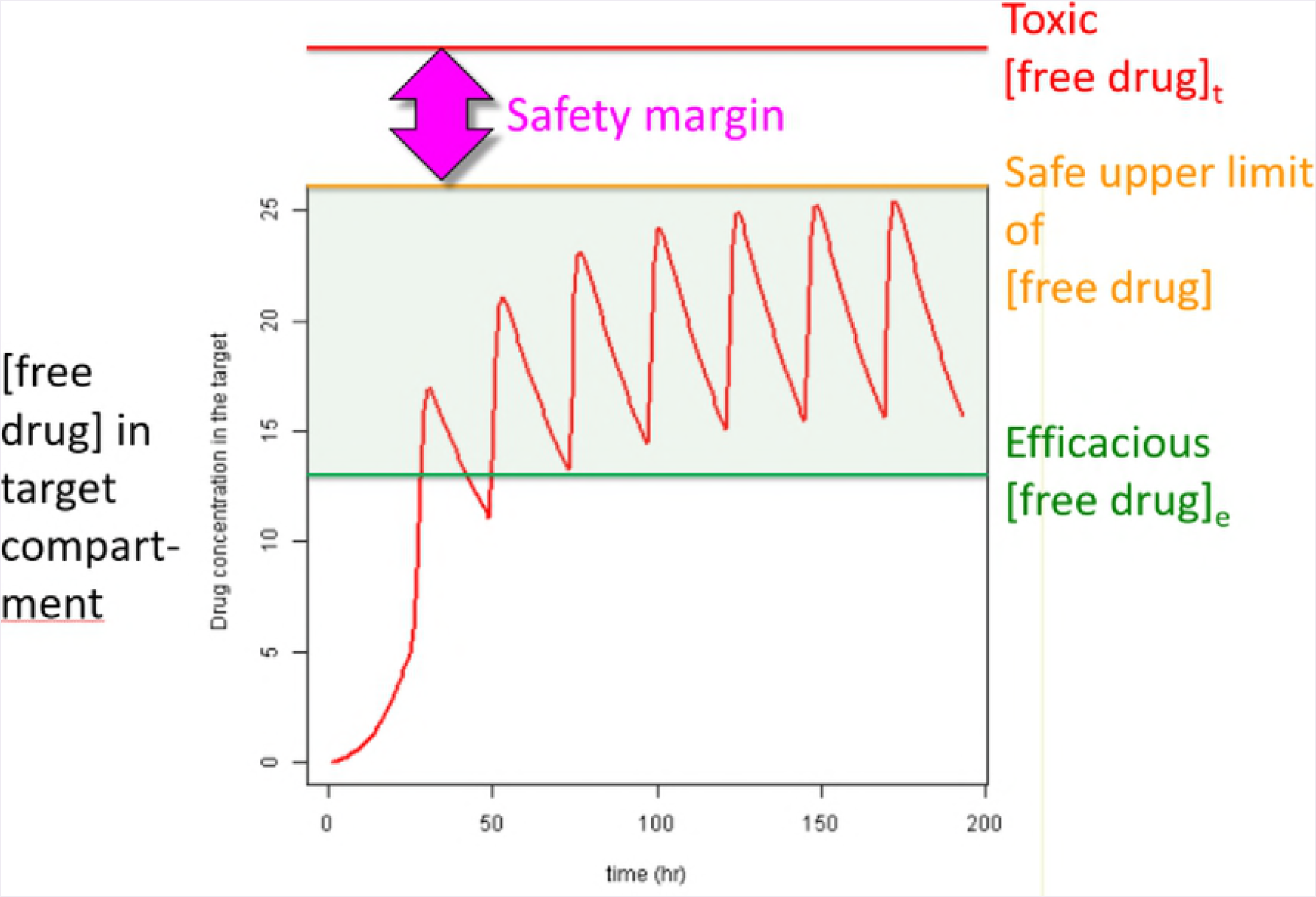
(A) Plot of *B_total_*(*τ*) (gray), *c*(*τ*) at a fixed *L_0_= K_d_* = 1 nM and Λ = 8,333 hr (*k_i_* = *k _− i_* = 1×10^-7^ s^-1^), in which <u>*k_on_* was sampled between 1×10^3^ M^-1^ s^-1^ to 1×10^5^ M^-1^ s^-1^</u> (solid lines color-coded from blue to red as a function of increasing *k_on_*), and *B_free_*(*τ*) (dotted lines color coded the same as *c*(*τ*)). *c*(*τ*) diverges slightly from SSO at *k_on_* = 1×10^3^ M^-1^ s^-1^. (B) Same as A, except Λ = 1,667 hr (*k_i_* = *k _− i_* = 5×10^-7^ s^-1^, *k_onSS_* = 1×10^3^ M^-1^ s^-1^). (C) Same as A, except Λ = 833 hr (*k_i_* = *k _− i_* = 1×10^-6^ s^-1^, *k_onSS_* = 1×10^4^ M^-1^ s^-1^). (D) Same as A, except Λ = 167 hr (*k_i_* = *k _− i_* = 5×10^-6^ s^-1^, *k_onSS_* = 5×10^4^ M^-1^ s^-1^). (E) Same as A, except Λ = 3 hr (*k_i_* = *k _− i_* = 1×10^-5^ s^-1^, *k_onSS_* = 1×10^5^ M^-1^ s^-1^). (F) Same as A, except Λ = 8.3 hr (*k_i_* = *k _− i_* = 1×10^-4^ s^-1^, *k_onSS_* = 1×10^6^ M^-1^ s^-1^). (G) Same as A, except Λ = 50 min (*k_i_* = *k _− i_* = 1×10^-3^ s^-1^, *k_onSS_* = 1×10^7^ M^-1^ s^-1^). (H) Same as A, except Λ = 5 min (*k_i_* = *k _− i_* = 1×10^-2^ s^-1^, *k_onSS_* = 1×10^8^ M^-1^ s^-1^). *k_on_* = 1×10^5^ M^-1^ s^-1^, the fastest *k_on_* sampled at this *k_i_* (1,000-fold < *k_onSS_*), results in nearly zero occupancy.

#### The non-equilibrium regime

*c*(*τ*) enters the non-equilibrium regime in our model at Λ ≌ 833 hr (*k_i_* = *k _− i_* = 1×10^-6^ s^-1^), where the SSO profile is achieved at *k_on_* ≥ *k_onSS_* = 1×10^4^ M^-1^ s^-1^ (Fig 9). Suboptimal *k_on_* results in the nSSO profile and loss of the signature saw-tooth morphology. *c*(*τ*) lags behind, and decays ahead of *B_total_*(*τ*), and c_50_ and c_95_ are no longer achieved.

Higher steady-state occupancies are achieved with decreasing Λ and *k_on_* in the range of 1×10^5^ to 1×10^7^ M^-1^ s^-1^ (Fig 10).

**Fig 10.**
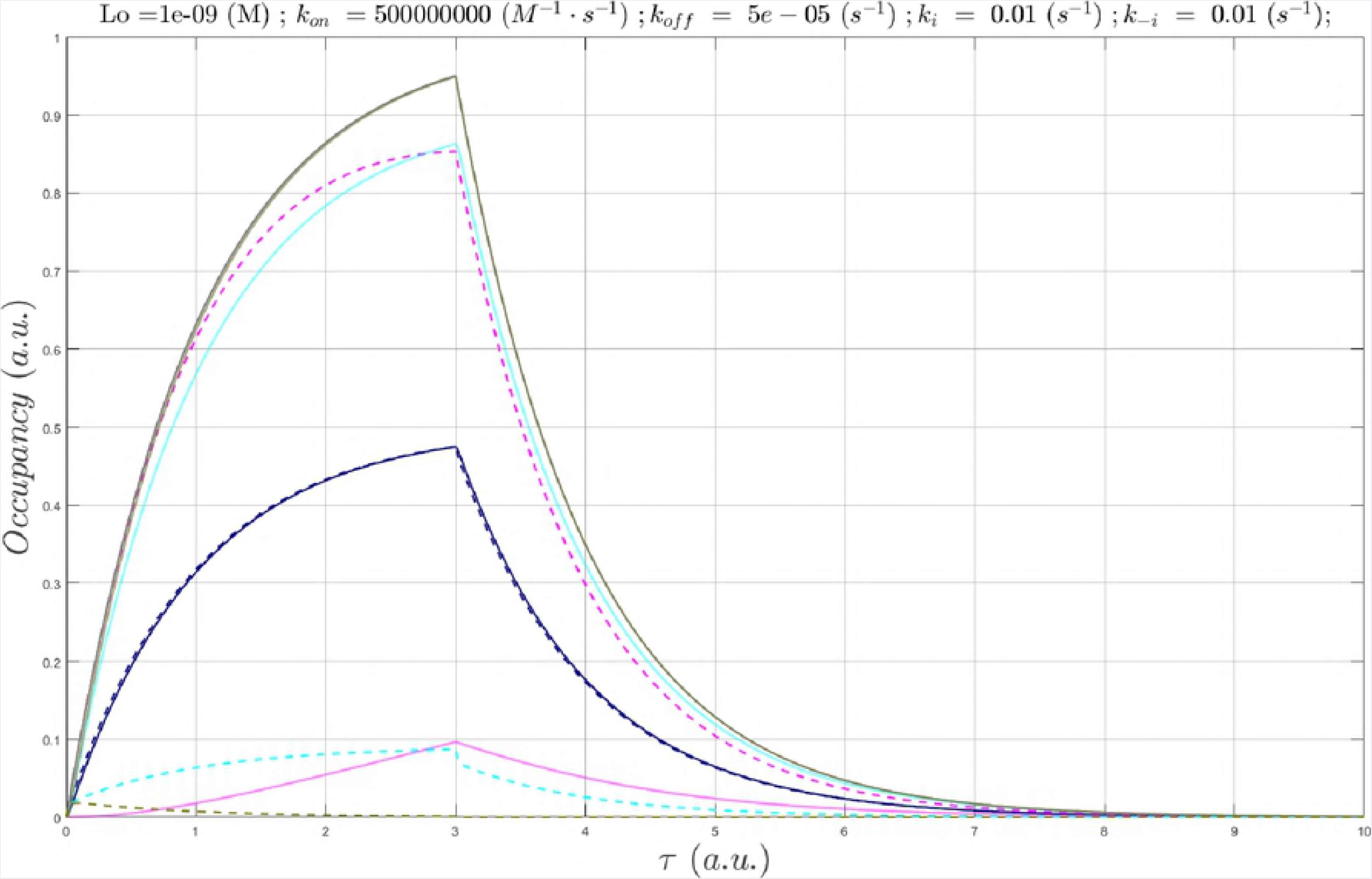
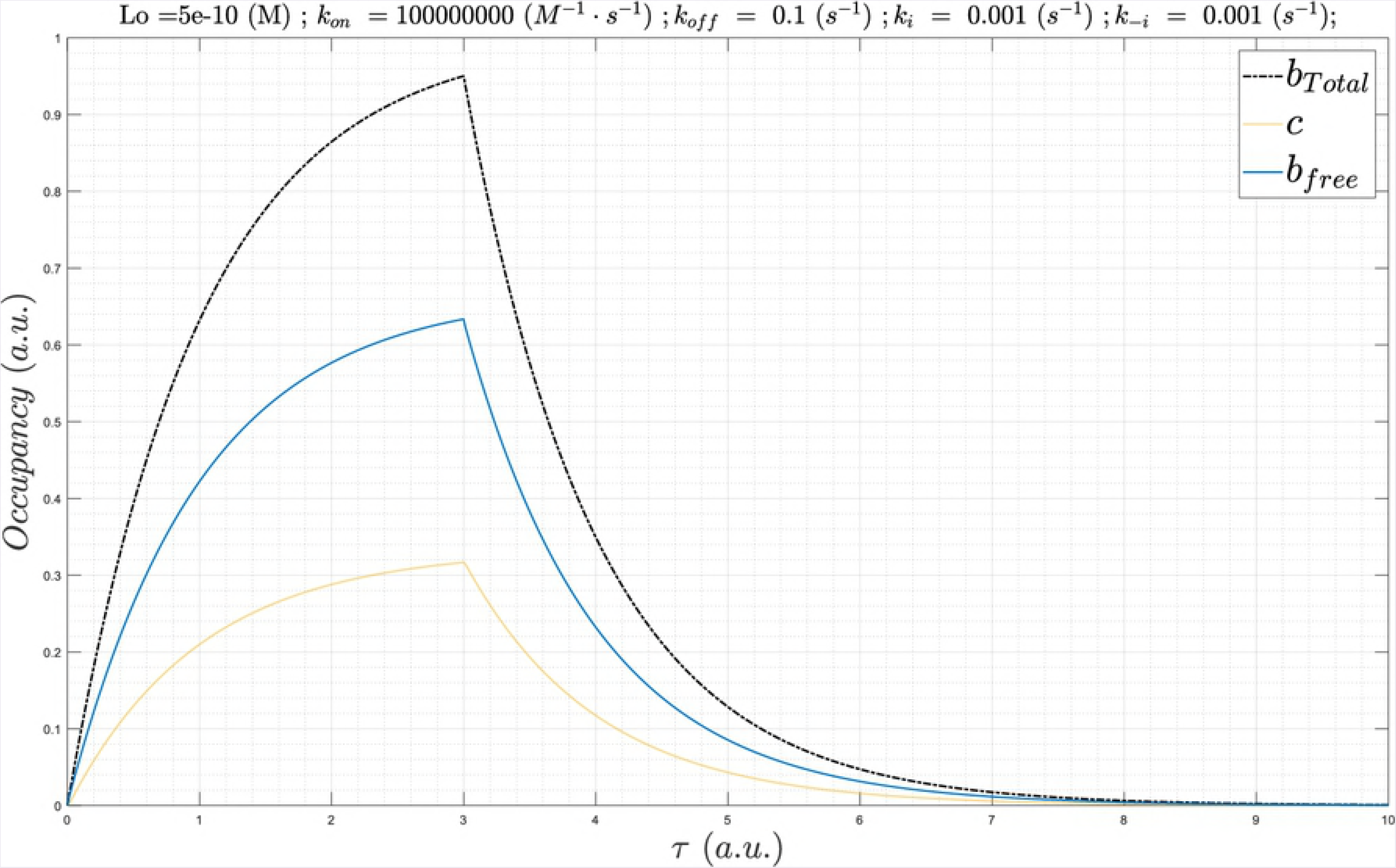
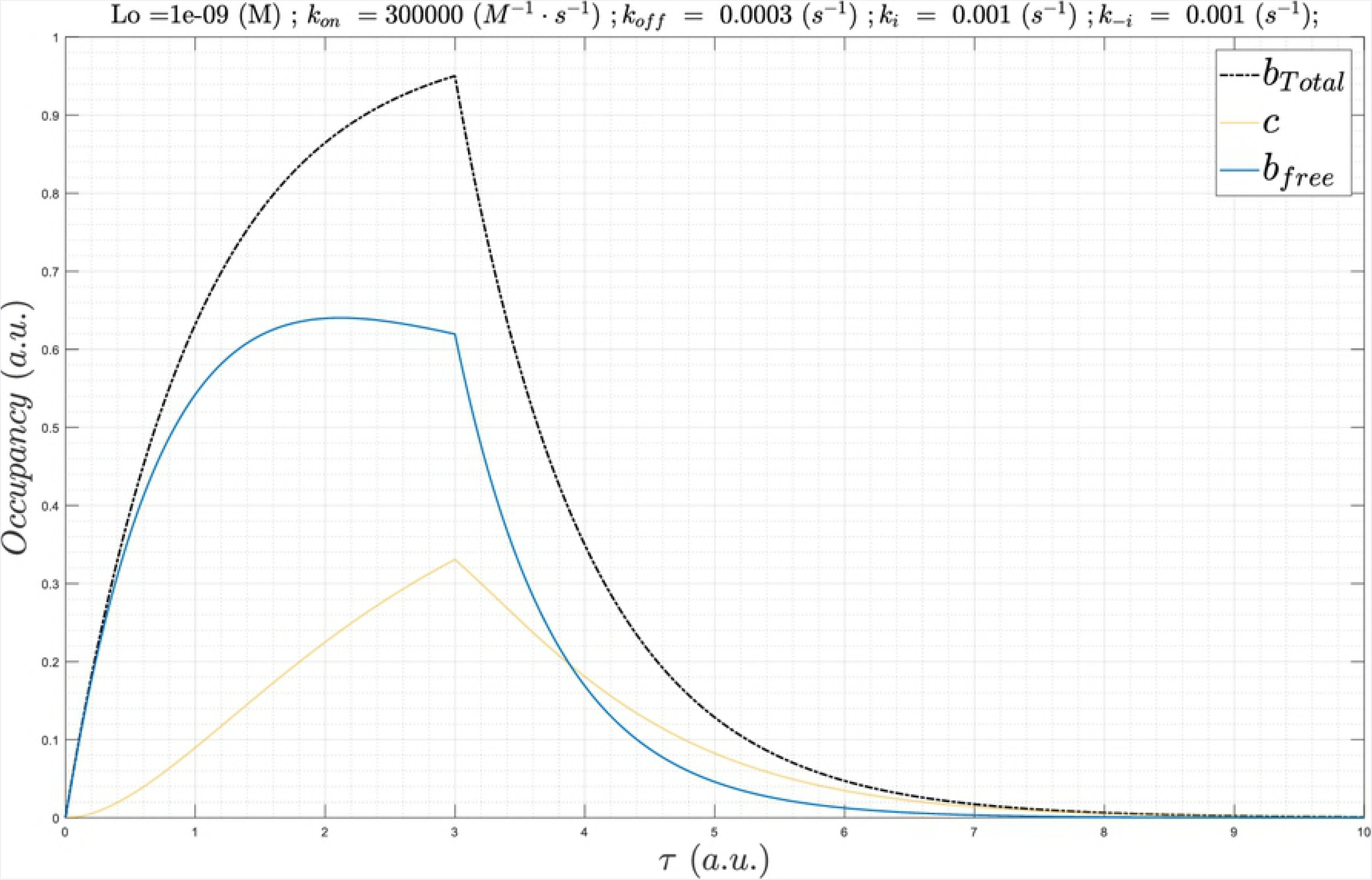
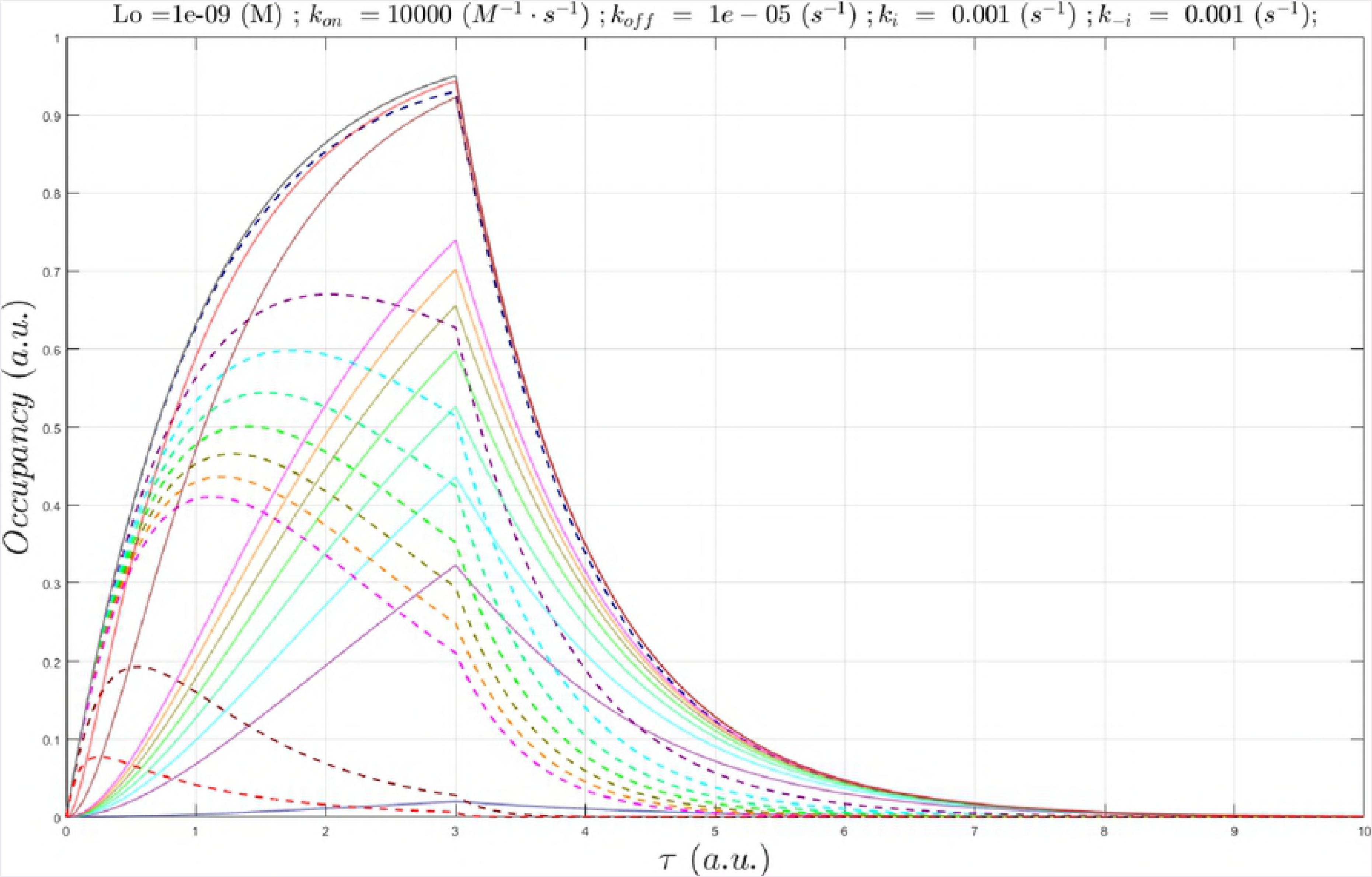
(A) Plot of *B_total_*(*τ*) (gray), *c*(*τ*) at a fixed *L_0_* = *K_d_* = 1 nM and Λ = 8.3 hr (*k_i_* = *k_− i_* = 1×10^-4^ s^-1^, *k_onSS_* = 1×10^6^ M^-1^ s^-1^), in which <u>*k_L_™* was sampled between 1×10^5^ M^-1^ s^-1^ to 1×10^7^ M^-1^ s^-1^</u> (solid lines color-coded from blue to red according to increasing *k_on_*), and *B_free_*(*τ*) (dotted lines color coded the same as *c*(*τ*)). (B) Same as A, except Λ =50 min (*k_i_* = *k_− i_* = 1×10^-3^ s^-1^). (C) Same as A, except Λ = 5 min (*k_i_* = *k_− i_* = 1×10^-2^ s^-1^ *k_onSS_* = 1×10^8^ M^-1^ s^-1^). (D) Same as A, except Λ = 30 s (*k_i_* = *k_− i_* = 1×10^-1^ s^-1^, *k_onSS_* = 1×10^9^ M^-1^ s^-1^). (E) Same as A, except Λ = 3 s (*k_i_* = *k_− i_* = 1×10^0^ s^-1^, *k_onSS_* = 1×10^10^ M^-1^ s^-1^). *k_on_* = 1×10^7^ M^-1^ s^-1^, the fastest *k_on_* sampled at this *k_i_* (1,000-fold < *k_onSS_*), results in nearly zero occupancy.

Additional simulation results as a function of Λ, *k_on_*, and *k_off_* are provided in the Supplementary Information (S1-S4 Figs).

#### Achieving the SSO profile depends on fast k_on_ even for long-lived binding sites

We assume that long lifetimes of cognate partners are achieved via fast buildup and slow decay of the binding-competent structural state or species levels, the latter of which may consist of synthesis and degradation or inter-compartmental shuttling (noting that species dilution caused by cell growth is not necessarily well described by our exponential functions). As is apparent from Fig 11, achieving steady-state occupancy remains dependent on *k_on_* ≥ *k_onSS_*, even at very slow binding site decay rates. At *k_on_* < *k_onSS_*, *c*(*τ*) may converge to *B_total_*(*τ*) far beyond the *B_max_* time point.

**Fig 11.**
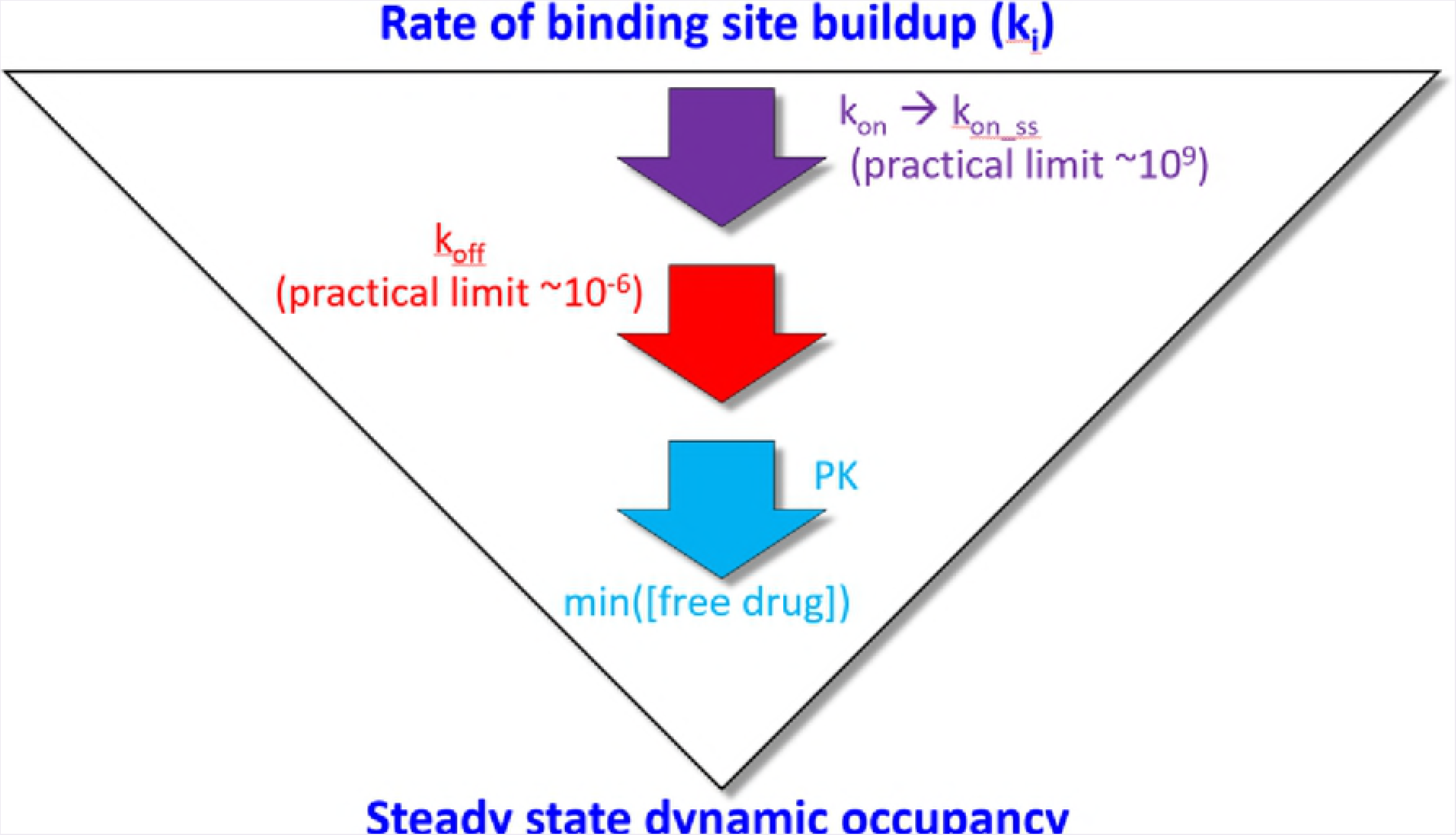
(A) Plot of *B_total_*(*τ*) (black), *B_free_*(*τ*) (blue), and *c*(*τ*) (gold) for Λ = 199 hr (*k_i_* = *k_− i_* = 1×10^-4^ s^-1^ 1×10^-6^ s^-1^, *k_onSS_* = 1×10^6^ M^-1^ s^-1^), *L_0_* = *K_d_* = 1 nM, *k_off_* = 1×10^-5^ s^-1^, and *k_on_* = 1×10^4^ M^-1^ s^-1^. (B) Same as A, except Λ = 199 hr (*k_i_* = 1×10^-4^ s^-1^, *k_− i_* = 1×10^-6^ s^-1^, *k_onSS_* = 1×10^6^ M^-1^ s^-1^), *k_off_* = 1×10^-3^ s^-1^, and *k_on_* = 1×10^6^ M^-1^ s^-1^. In this example, approximate SSO is achieved at this *k_on_*. (C) In this example, *k_on_* is too slow to keep up with *B_total_*(*τ*), but *c*(*τ*) converges to steady-state during the decay phase. Same as A, except Λ = 20 hr (*k_i_* = 8×10^-4^ s^-1^, *k _− i_* = 1×10^-5^ s^-1^, *k_onSS_* = 1×10^6^ M^-1^ s^-1^), *k_off_* = 1×10^-3^ s^-1^, and *k_on_* = 1×10^6^ M^-1^ s^-1^. (D) Same as A, except Λ = 2.1 hr (*k_i_* = 3×10^-3^ s^-1^, *k _− i_* = 1×10^-4^ s^-1^, *k_onSS_* = 1×10^7^ M^-1^ s^-1^), *k_off_* = 1×10^-3^ s^-1^, and *k_on_* = 1×10^6^ M^-1^ s^-1^. Approximate SSO is achieved at this *k_on_*.

### Scenario 2

Binding site buildup and decay in this scenario are assumed to depend on fast conformational dynamics relative to the lifetime of the binding site-containing species, which may range case-by-case from ms (e.g. voltage-gated ion channels) to hr. We assume that scenarios 1 and 2 play different roles in biodynamics (i.e. the generation of non-equilibrium conditions and MDEs, respectively). However, our overall conclusions do not depend on this assumption. As for scenario 1, we set about to characterize the effect of decreasing time-dependent binding site availability on *c*(*τ*).

#### The hypothetical equilibrium regime

As for scenario 1, the SSO profile is achieved in the equilibrium case in a kinetics-agnostic manner (Λ = 3,851 hr and *k_i_* = *k _− i_* = 1×10^-7^ s^-1^) (S5 Fig). Under these conditions, the bound fraction is 50% or 95% (*L_0_* = 1 nM or 19 nM, respectively) at all instants of time, and *c*(*τ*) exhibits the same signature bi-sigmoidal morphology as that of *B_total_*(*τ*) (S5 Fig).

#### The non-equilibrium regime

As for scenario 1, faster *k_on_* is needed to achieve the SSO profile as *k_i_* becomes progressively faster. The nSSO profile, and loss of the signature bi-sigmoidal *c*(*τ*) morphology, result from suboptimal *k_on_*. In such cases, c_max_ may no longer reach c_50_ or c_95_, and *c*(*τ*) lags behind, and decays ahead of *B_total_*(*τ*). *c*(*τ*) enters the non-equilibrium regime at Λ = 833 hr (*k_i_* = *k_− i_* = 1×10^-6^ s^-1^), where a minimum *k_on_* of ~1×10^5^ M^-1^ s^-1^ is required to fully achieve the SSO profile (Fig 12).

**Fig 12.**
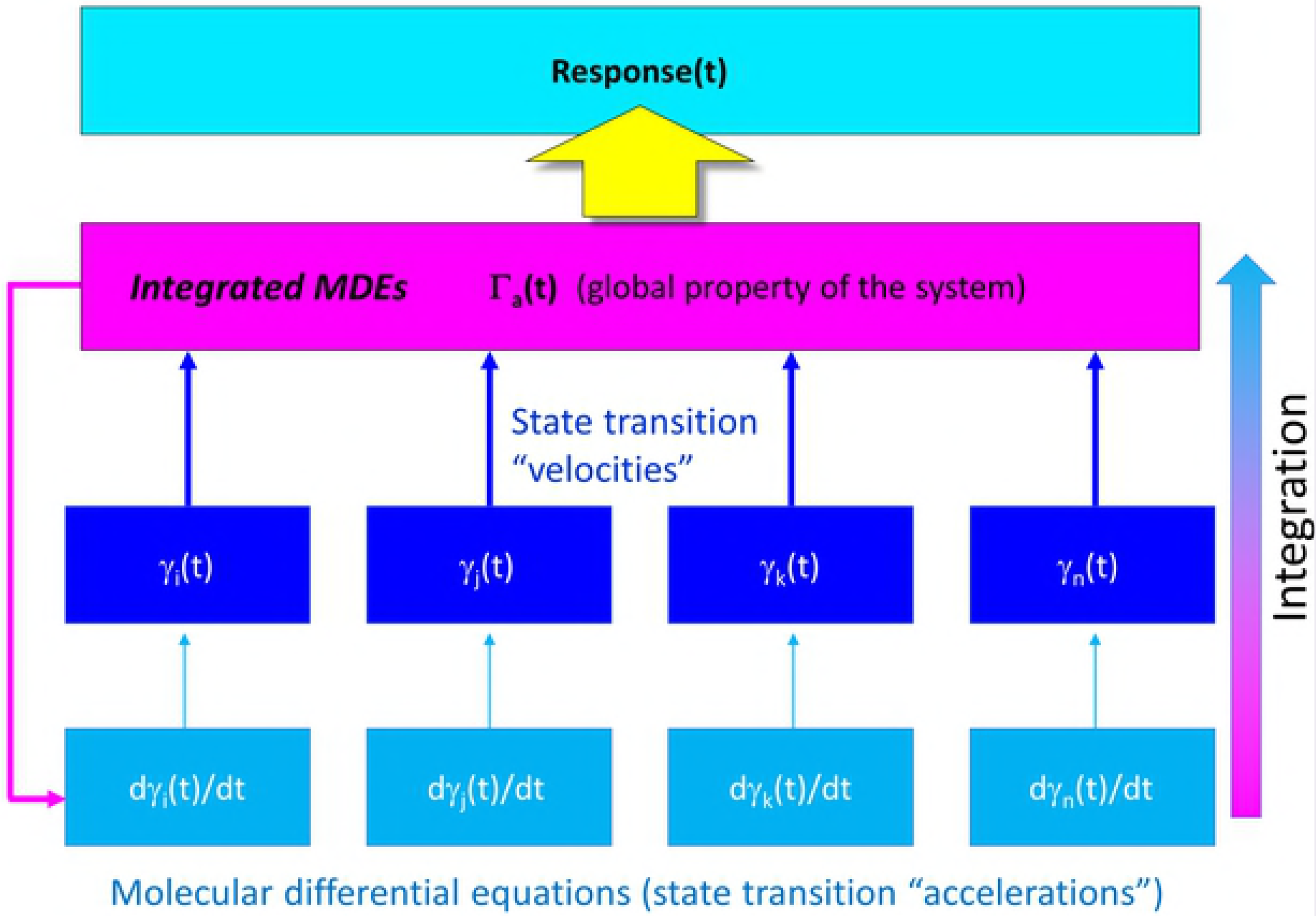
(A) Plot of *B_total_*(*τ*) (gray), *c*(*τ*) at a fixed *L_0_* = 1 nM and Λ = 3,851 hr (*k_i_* = *k_− i_* = 1×10^-7^ s^-1^), in which <u>*k_on_* was sampled between 1×10^3^ M^-1^ s^-1^ to 1×10^5^ M^-1^ s^-1^</u> (solid lines color-coded from blue to red according to increasing *k_on_*), and *B_free_*(*τ*) (dotted lines color coded the same as *c*(*τ*)). The *c*(*τ*) are beginning to diverge from the steady-state at this Λ. (B) Same as A, except Λ = 385 hr (*k_i_* = *k_− i_* = 1×10^-6^ s^-1^). (C) Same as A, except Λ = 3.85 hr (*k_i_* = *k _− i_* = 1×10^-4^ s^-1^). (D) Same as A, except Λ = 23.1 min (*k_i_* = *k _− i_* = 1×10^-3^ s^-1^). *k_on_* = 1×10^5^ M^-1^ s^-1^, the fastest *k_on_* sampled at *k_i_* = *k _− i_* = 1×10^-2^ s^-1^, results in nearly zero occupancy (not shown).

Additional simulation results as a function of Λ, *L*_0_, *k_on_*, and *k_off_* are provided in S6-S7 Figs.

### The implications of binding dynamics for known short-lived binding sites

We next assessed the implications of binding dynamics for hERG channel blockade and LDL receptor (LDL-R) binding to wild type (*wt*) versus the disease-causing gain-of-function D374Y mutant form of PCSK9.

#### PCSK9-LDL receptor

We characterized PCSK9-LDL-R dynamic occupancy based on our scenario 1 model and the pH-dependent binding kinetics data reported in [12] (summarized in Table 2). Rapid buildup of PCSK9 can be inferred from its observed ~5 min half-life [24], which is driven by zymogen activation [25] rather than *de novo* expression (as may be the case for many short Λ species). Negligible PCSK9-LDL-R occupancy at neutral pH follows from the exceptionally high *K_d_* relative to circulating plasma PCSK9 (which ranges between 30-3,000 ng/ml in humans (~405 pM to 40.5 nM) [26]), suggesting that extracellular binding depends on other factors. Binding and LDL-R degradation have indeed been shown to depend on heparin sulfate proteoglycans present on the extracellular surface of hepatocytes [27]. On the other hand, significant nSSO is achieved at the upper end of the concentration range for both *wt* and mutant forms of PCSK9 at lysosomal pH (Fig 13), suggesting that gain-of-function mutations act either through relatively small increases in dynamic occupancy relative to wt, or through some other means (e.g. by slowing PCSK9 degradation).

**Table 2.** Measured binding kinetics data for *wt* and D374Y mutant PCSK9-LDL-R [12]. Fold differences between *k_on_* and *k_off_* for D374Y versus *wt* are shown in parenthesis, noting that the *t*_1/2_ of the bound complex is similar to that of *wt* PCSK9, but considerably longer than that of the D374Y mutant (which is likely subject to *k_off_* hijacking). The ratio of *k_onSS_*/*k_on_* was calculated based on *k_onSS_* = 1×10^8^ M^-1^ s^-1^, corresponding to *k_i_* = 1×10^-2^ s^-1^ (predicted from equation 13 under the assumption that *k_i_* = *k _− i_*).

|  | *wt* |  | D374Y |  |
| --- | --- | --- | --- | --- |
|  | pH 7.4 | pH5.3 | pH 7.4 | pH5.3 |
| $k_{on}$<br>($\text{M}^{-1} \text{ s}^{-1}$) | $1.86 \times 10^3$ | $4.73 \times 10^5$ | $4.57 \times 10^3$<br>(+2.5-fold) | $6.74 \times 10^5$<br>(+1.4-fold) |
| $k_{onSS}/k_{on}$ | 53,763 | 211 | 21,881 | 148 |
| $k_{off}$<br>($\text{s}^{-1}$) | $1.17 \times 10^{-3}$ | $1.97 \times 10^{-3}$ | $4.64 \times 10^{-4}$<br>(-2.5-fold) | $3.64 \times 10^{-4}$<br>(-5.4-fold) |
| $k_{off}'$<br>($\text{s}^{-1}$) | $1.17 \times 10^{-3}$ | $1.97 \times 10^{-3}$ | $2.3 \times 10^{-3\ddagger}$ | $2.3 \times 10^{-3\ddagger}$ |
| $K_d$<br>($\text{M}$) | $628 \times 10^{-9}$ | $4.19 \times 10^{-9}$ | $101 \times 10^{-9}$ | $0.54 \times 10^{-9}$ |
| $K_d'$<br>(M) | 628x10<sup>-9</sup> | 4.19x10<sup>-9</sup> | 503x10<sup>-9</sup> | 3.4x10<sup>-9</sup> |

§Similar clearance rates are assumed for both mutant and *wt* PCSK9.

**Fig 13.**
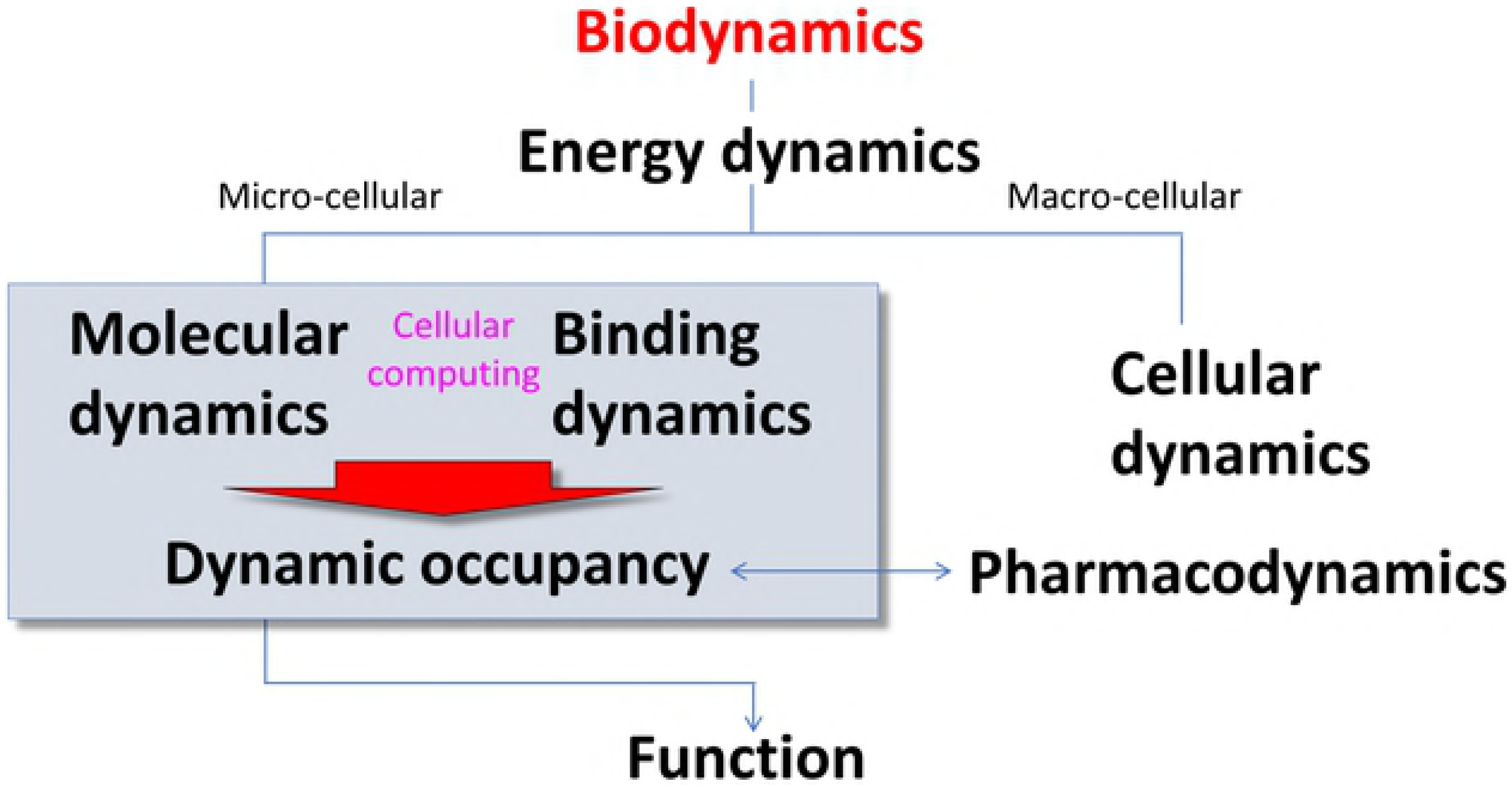
(A) Plot of the hypothetical *B_total_*(*τ*) (gray) and *c*(*τ*) for PCSK9 and PCSK9-LDL-R binding at a fixed PCSK9 *L_0_* = 1 nM, Λ = 5 min (*k_i_* = *k_-i_* = 1×10^-2^ s^-1^), and the *k_on_* and *k_off_* values given in Table 2 (solid lines color-coded as follows: wt/pH 7.4 dark blue (not visible); *wt*/pH 5.3 purple; D374Y/pH 7.4 cyan; D374Y/pH5.3 green), and *B_free_*(*τ*) (dotted lines color coded the same as *c*(*τ*)). Simulated occupancy is essentially zero for both *wt* and mutant PCSK9 at neutral pH, whereas negligible nSSO is achieved at low pH. (B) Same as A, except *L_0_* = 20 nM. Moderate nSSO is achieved at low pH (maximum occupancy = 68% versus 75% for the *wt* and mutant forms, respectively). (C) Same as A, except *L_0_* = 40.5 nM. High nSSO is achieved at low pH, where the gap between wt and mutant forms narrows to only a few percent at all time points (maximum occupancy = 82% versus 85% for the wt and mutant forms, respectively).

#### hERG channel blockade by non-trappable compounds

Certain hERG blockers are trapped within closed channels (which we refer to as “trappable”), whereas others are expelled during channel closing (which we refer to as “non-trappable”) [4,28]. Trappable blocker occupancy builds to *K_d_* (the specific time-dependence of which depends on *k_on_*), whereas that by non-trappable blockers builds and decays during each gating cycle. The ultrashort ~350 ms lifetime of the open and inactivated channel states results in *k_off_* hijacking, in which 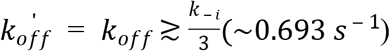 and *k_off_* ≥ *k _− i_* (~2 *s*^-1^) for the trappable and non-trappable cases, respectively [4]. In our previous work, we characterized the dynamic occupancy of a set of hypothetical non-trappable blockers using an alternate model, consisting of the O’Hara-Rudy action potential (AP) simulator, in which we replaced the Hodgkin-Huxley hERG model with a Markov state model incorporating blocker binding [4]. Here, we used the predicted pro-arrhythmic 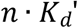 versus *n* · *K_d_* as a function of *k_on_*, *k_off_*, and 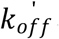 as a metric of dynamic occupancy, and qualitatively compared the results with our binding dynamics models (Fig 14 versus S6(E) Fig and Table 3 versus Table 1).

**Fig 14.**
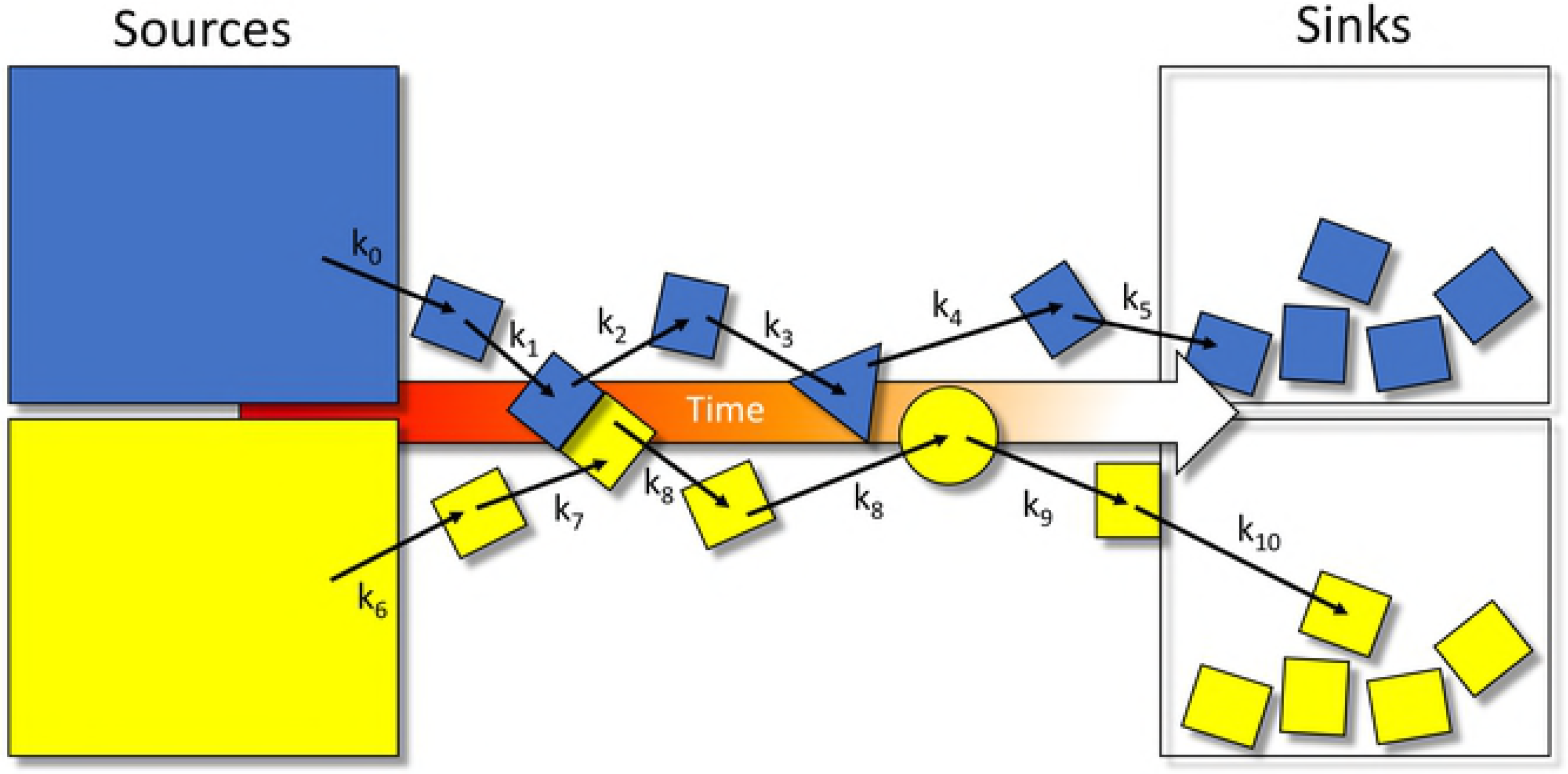
(A) Plot of dynamic fractional hERG occupancy in mid-myocardial (M) cells by a hypothetical non-trappable blocker exhibiting *k_on_* = 1×10^5^ M^-1^ s^-1^ and *k_off_* = 5×10^-1^ s^-1^ (*K_d_* = 5 μM and 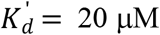) sampled at 10 and 25 μM (green solid and dashed curves, respectively). The total dynamic binding site level consists of the sum of the open/conducting and inactivated channel populations (gray dotted, solid, and dashed curves corresponding to 0, 10, and 25 μM blocker concentrations, respectively). The overall shape of the channel state population curves qualitatively resembles our logistic model (in which *kι* is very fast), noting that recovery from inactivation (the decay region of the curves) is slowed by blockade due to the smaller contribution of the hERG current to the membrane potential, which in turn, alters the response of the channel population (consistent with the recursive effect depicted in Fig 2). The blocker occupancy curves reflect nSSO occupancy at *k_on_* = 1×10^5^ M^-1^ s^-1^ (where *k_on_* ≫ *k_onSS_*, as suggested from Table 1), and the peak occupancy is far below 50% at 5 μM. Nevertheless, even transient fractional occupancy approaching the ~45% level can be highly pro-arrhythmic [4]. (B) Plot of *B_total_*(*τ*) (gray), *B_free_*(*τ*) (blue) and *c*(*τ*) (gold) simulated using our logistic model (scenario 2) for a hERG blocker exhibiting *K_d_* = 5 μM (*k_on_* = 1×10^5^ M^-1^ s^-1^ and *k_off_* = 0.5 s^-1^) and [free blocker] = 10 μM. *k_i_* and *k_-i_* were set to 13 s^-1^ and 2.1 s^-1^, respectively, so as to reproduce *B_max_* at *τ* ≈ 50 ms and *Λ* ≈ 350 ms. *B_total_*(*τ*) and *c*(*τ*) are qualitatively similar to the curves in A.

**Table 3.** *L*_0_ (expressed as *n*·*K_d_* and *n*· *K_d_*) for a set of hypothetical hERG blockers resulting in the pro-arrhythmic fractional channel occupancy levels predicted from our AP simulations. Blocker *K_d_* = 5 μM in all cases, where *k_on_* ranges between 1×10^4^ and 1×10^9^ M^-1^ s^-1^ and *k_off_* between 0.005 and 5,000 (noting that the pro-arrhythmic occupancy level increases as *k_off_* speeds). *n* · *K_d_* was back-calculated from the predicted pro-arrhythmic occupancy using equation 19.

|  |  |  |  |  |  |  |
| --- | --- | --- | --- | --- | --- | --- |
| $k_{on}$ (M<sup>-1</sup> s<sup>-1</sup>) | $1 \times 10^4$ | $1 \times 10^5$ | $1 \times 10^6$ | $1 \times 10^7$ | $1 \times 10^8$ | $1 \times 10^9$ |
| $k_{off}$ (s<sup>-1</sup>) | $5 \times 10^{-2}$ | $5 \times 10^{-1}$ | $5 \times 10^0$ | $5 \times 10^1$ | $5 \times 10^2$ | $5 \times 10^3$ |
| $k_{off}'$ (s<sup>-1</sup>) | 2 | 2 | 2 | 2 | 2 | 2 |
| **Fractional pro-arrhythmic hERG occupancy (%)** | 42 | 45 | 53 | 54 | 55 | 55 |
| **$n \cdot K_d' \rightarrow$ pro-arrhythmic occupancy (M)** | $2.6 \times 10^{-4}$ | $2.9 \times 10^{-5}$ | $7.8 \times 10^{-6}$ | $6.2 \times 10^{-6}$ | $6.0 \times 10^{-6}$ | $6.0 \times 10^{-6}$ |
| **$n \cdot K_d \rightarrow$ pro-arrhythmic occupancy (M)** | $3.6 \times 10^{-6}$ | $4.1 \times 10^{-6}$ | $5.7 \times 10^{-6}$ | $5.9 \times 10^{-6}$ | $6.1 \times 10^{-6}$ | $6.1 \times 10^{-6}$ |

### Implications of binding dynamics for drug discovery

Both drug-target and endogenous time-dependent binding partner occupancy are described by similar biodynamics principles. Namely:

1. Extrinsic rates: the rates of buildup and decay of the binding site and ligand, which may vary with conditions.
2. Intrinsic rates: *k_on_* and *k_off_*, which may also vary with conditions.
3. Fractional occupancy at each instant of time (*c*(*τ*)). It is reasonable to assume that molecular response (both efficacious and toxic) depends on *c*(*τ*) ≥ a fractional occupancy threshold during each binding site buildup/decay cycle (in general, ranging from a small fraction to nearly 100%, case-by-case).

Our results suggest the paramount importance of tuning *k_on_* and *k_off_* to the rates of binding site buildup and decay, respectively, for achieving the Goldilocks zone of efficacious target occupancy and target/off-target selectivity (Fig 15). Constant maximal <u>fractional</u> occupancy is maintained under the SSO profile <u>at the lowest possible</u> *L* (where *c*(*τ*) depends solely on *L* = *n*· *K_d_*). On the other hand, fractional occupancy in the qSSO regime approaches a constant level only under saturating conditions (i.e. at n ≫ 1) (S3 Fig). Target/off-target selectivity may be compromised with the nSSO profile, depending on the ratio of *K_d_*(*_target_*)/*K*_*d*(*0ff-target*)_, keeping in mind that nSSO may exist for the target, and SSO for one or more off-targets and/or competing endogenous substrates.

**Fig 15.**
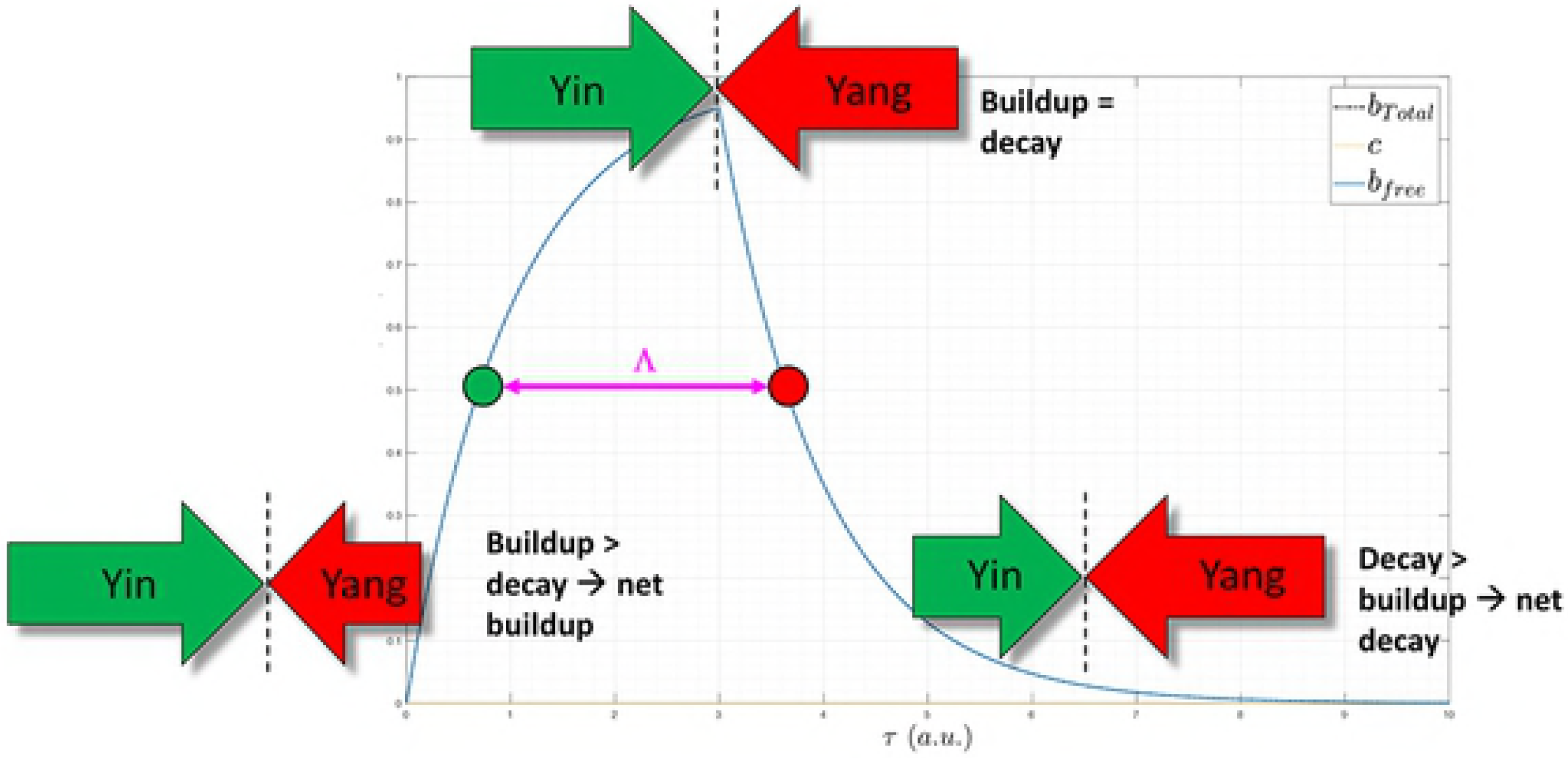
Hypothetical plot of drug *L* versus time (red), showing a series of buildup-decay cycles (one per dose) culminating in a steady-state condition, together with the efficacious threshold of *L* (green), the safe upper limit of *L* (orange), and the toxic level of *L* (red). *L* is maintained in the efficacious/sub-toxic “green zone” by overshooting and decaying back to the efficacious level. Optimizing to the SSO regime affords the greatest chance of achieving a therapeutic index in humans. *Tuning kinetics to the qSSO profile*

#### Tuning kinetics to the qSSO profile

Optimization of drug *k_off_* or “residence time” (i.e. *t*_1/2_ = ln (2)/*k_off_*) does not translate to increased occupancy when *k_off_* < *k_i_* (unless *k_-i_* is slowed in the bound state). We used equation 10 to test the effect of slowing *k_off_* under qSSO versus SSO conditions (i.e. *k_on_ < k_on_ss_* versus *k_on_* ≥ *k_on_ss_*) at constant *K_d_* (Fig 16).

**Fig 16.**
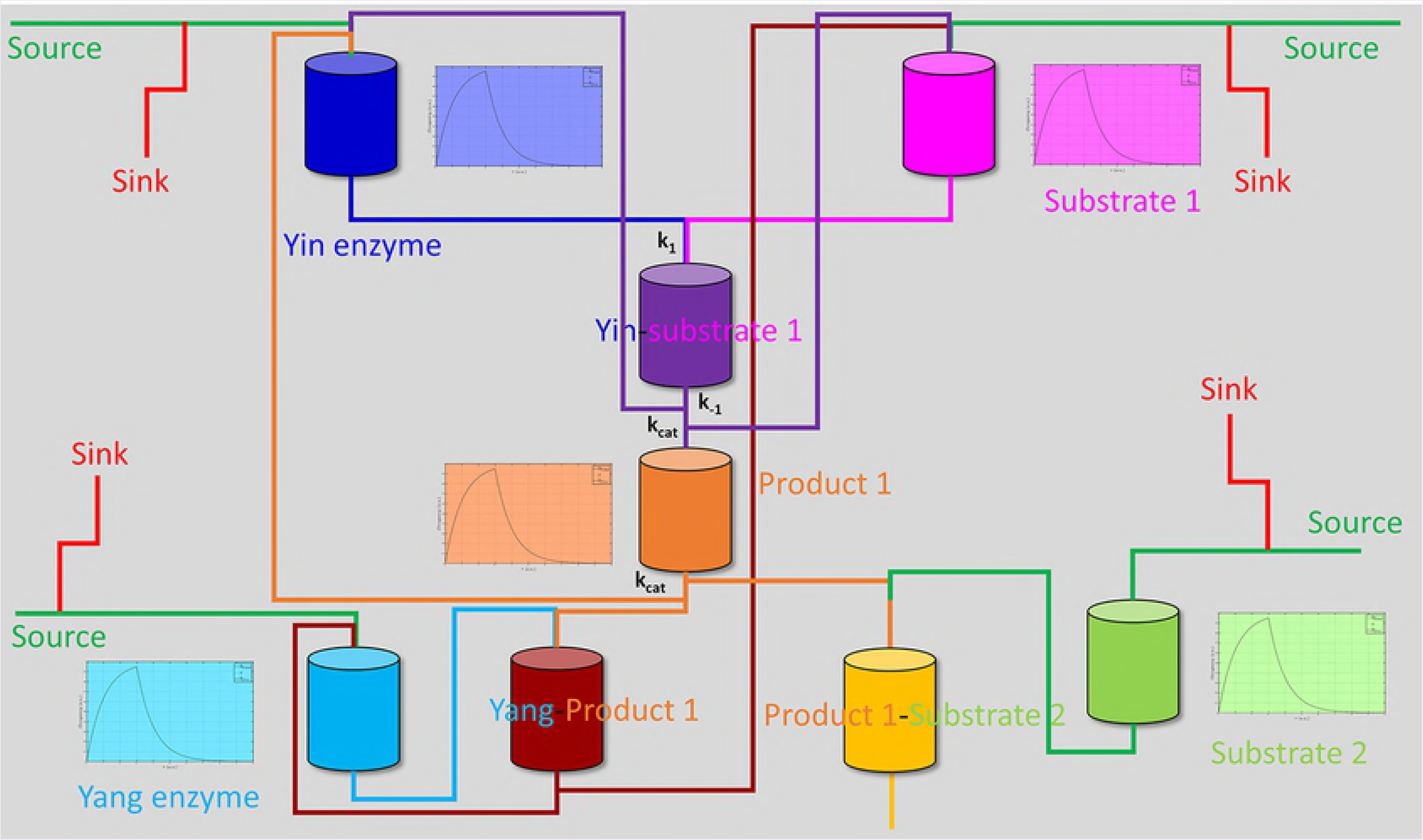
Plot of *B_total_*(*τ*) (gray), *c*(*τ*) at *L*_0_ = *k_off_*/*k_on_* = 1 nM and Λ = 5 min (*k_i_* = *k_-i_* = 1×10^-2^ s^-1^), <u>in which *k_off_* and *k_on_* were sampled at 5×10^-1^ s^-1^/5×10^8^ M^-1^ s^-1^ (blue), 5×10^-7^ s^-1^/5×10^5^ M^-1^ s^-1^ (magenta), 5×10^-2^ s^-1^/5×10^8^ M^-1^ s^-1^ (cyan), and 5×10^-5^ s^-1^/5×10^8^ M^-1^ s^-1^</u> (olive) (solid lines), and *B_free_*(*τ*) (dotted lines correspond to the *c*(*τ*) color scheme). *c*(*τ*) is unaffected by slowing *k_off_* (magenta curve) when *k_on_* < *k_onSS_*. However, when *k_on_* ≥ *k_0nss_*, *c*(*τ*) increases when *k_off_* is further slowed (e.g. 5×10^-1^ s^-1^ → 5×10^-2^ s^-1^ (cyan curve) and 5×10^-1^ s^-1^ → 5×10^-5^ s^-1^ (olive curve)).

Lastly, we used equation 10 to test the effect of increasing *L*_0_ toward the qSSO profile at *k_on_* < *k_on_ss_* (Fig 17 and S2 Fig).

**Fig 17.**
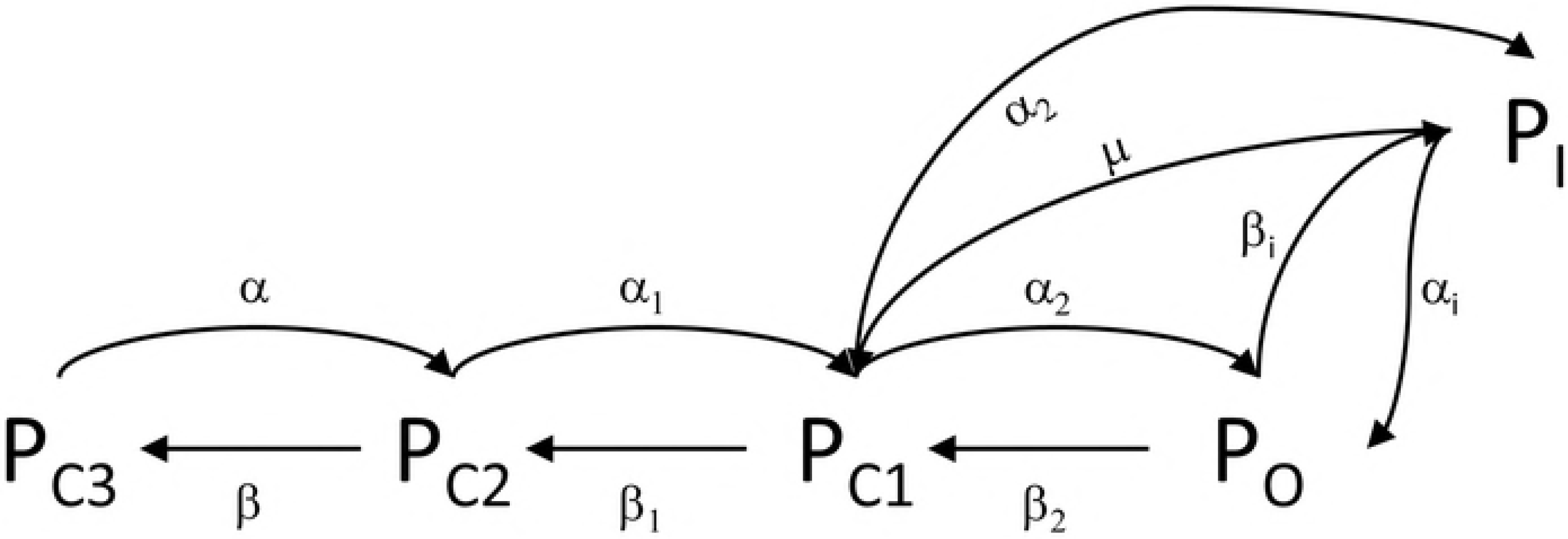
(A) Plot of *B_total_*(*τ*) (black), *B_free_*(*τ*) (blue), and *c*(*τ*) at *L*_0_ = 0.5 · *K_d_* = 0.5 nM and Λ = 5 min (*k_i_* = *k_-i_* = 1×10^-3^ s^-1^, *k_onSS_* = 1×10^8^ M^-1^ s^-1^), where *k_off_* = 1×10^-1^ *s^Λ^/k_on_* = 1×10^8^ M^-1^ s^-1^ (gold). (B) Same as A, except at *L*_0_ = *K_d_* = 1 nM, and *k_off_* = 3×10^-4^ s^-1^/*k_on_* = 3×10^5^ M^-1^ s^-1^. Although *c*(*τ*) reaches ~c_31_ in both cases, the steady-state scenario in A reaches ~c_31_ at all instants of time (an example of kinetically tuned binding), whereas the nSSO scenario results in considerably greater *B_free_*(*τ*) prior to c_max_ (an example of non-kinetically tuned binding). (C) Plot of *B_total_*(*τ*) (gray), *c*(*τ*) at *K_d_* = 1 nM, *Λ =5* min (*k_i_* = *k_− i_* = 1×10^-3^ s^-1^, *k_onSS_* = 1×10^8^ M^-1^ s^-1^), *k_off_* = 1×10^-5^ s^-1^, and *k_on_* = 1×10^4^ M^-1^ s^-1^, in which *L*_0_ was sampled between 1 nM and 2 μM (solid lines), and *B_free_*(*τ*) (dotted lines correspond to the *c*(*τ*) color scheme). Slow *k_on_* is only compensated at sufficiently high *L*_0_, which in this case, approaches SSO conditions at *L_0_ =* 1 μM.

#### The putative effects of competition and drug pharmacokinetics (PK) on drug-target occupancy

Although we have not considered the effects of endogenous ligand competition and time-dependent drug concentration (L(t)) on dynamic drug-target occupancy, these factors can only serve to further increase *k_on_ss_*. The rates of binding site and drug buildup within the target-containing compartment (which may or may not be slower than the rate of drug buildup within the central compartment) should ideally be synchronized to ensure that *L* is always ≥ *K_d_*. Fractional drug-target occupancy is decreased by *L_e_* · *K_i_*/*K_e_* in the presence of endogenous ligand competition, as is apparent from the following equation [23]:

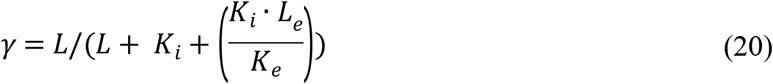

where *L*_e_, *K_e_*, and *K_i_* are the free endogenous ligand concentration and *K_d_*, and inhibitory drug *K_d_*, respectively. For example, drug SSO = 50% is achieved at *L*_50_ = *K_i_* + *L_e_* when *K_i_* = *K_e_*, and *L*_50_ further increases with increasing *L_e_* (or synchronization with binding site buildup) and/or decreasing *K_e_*. It is further apparent that qSSO is achieved at higher *L* than in the non-competitive case, and that competition between endogenous ligand binding via the SSO profile would be greatly favored over inhibitory drug binding via the nSSO profile (more the reason for optimizing to *k_on_* ≥ *k_on_ss_*).

#### The binding kinetics profiles of marketed drugs are consistent with non-equilibrium drug-target binding

We set about to test the relative importance of *k_on_ss_* versus *k_off_* in the human setting using measured *k_on_* and *K_d_* data curated by Dahl and Akrud for 32 marketed drugs [29] (reproduced in Table 3). We assume that marketed drugs observe steady-state or quasi-steady-state occupancy for targets with fast *k_i_* and *k_-i_*, or bind to targets that build and decay slowly (i.e. kinetics-agnostic occupancy). The data can be summarized as follows:

1. *k_on_* ranges between 1.4×10^1^ and 9.2×10^7^ M^-1^ s^-1^, with *k_on_* ≥ 1×10^5^ M^-1^ s^-1^ occurring in ~72% of the cases, sufficient for achieving the SSO profile for binding site buildup times between 1 m ≤ *t*_1/2_ ≤ 19 h (blue text in Table 3).
2. *k_off_* ranges between 4.8×10^-6^ and 2.8 s^-1^, with *k_off_* < 3.9×10^-4^ s^-1^ (t_1/2_ ≿ 30 min) occurring in ~47% of the cases (green text in Table 3).
3. *K_d_* < 1 nM was achieved in ~44% of the cases (red text in Table 3), which is due to *k_off_* < 3.9×10^-4^ s^-1^ in 57% of those cases (i.e. extreme potency is not driven by slow *k_off_*).
4. *k_off_* > 3.9×10^-4^ s^-1^ and *k_on_* > 1×10^5^ M^-1^ s^-1^ (i.e. *k_on_-d?ven* occupancy) occurs in ~44% of the cases.
5. *k_off_* < 3.9×10^-4^ s^-1^ and *k_on_* < 1×10^5^ M^-1^ s^-1^ (i.e. *k_0_ff-driven* occupancy) occurs in only ~19% of the cases, of which 4 are receptors, 2 are NS3 protease, and 1 is HIV-1 RT (noting that *k_on_* is near our cutoff in the receptor ligand telmisartan, and the NS3 inhibitor ciluprevir exhibits fast *k_on_)*.

**Table 3.** Published *k_on_*, *k_off_*, and *K_d_* values for a set of marketed drugs [29]. The drugs *are* grouped and color-coded *by target. Cases* of *K_d_ < 1* nM *are* color-coded with red text, *k_on_ >* 1×10^5^ M^-1^ s^-1^ *are* color-coded with blue text, and *t_1_/_2_ >* 30 min *are* color-coded with green *text*.

| Drug | Target | $k_{on}$<br>($\text{M}^{-1} \text{ s}^{-1}$) | $k_{off}$<br>($\text{s}^{-1}$) | $t_{1/2}$<br>(min) | $K_d$<br>(M) |
| --- | --- | --- | --- | --- | --- |
| Aliskiren | Renin | 4.00E+05 | 1.1E-04 | 105 | 3.00E-10 |
| Aclidinium | Muscarinic M3 receptor | 1.10E+06 | 2.0E-05 | 578 | 1.80E-11 |
| Ipratropium | Muscarinic M3 receptor | 4.10E+06 | 1.1E-03 | 11 | 2.60E-10 |
| Tiotropium | Muscarinic M3 receptor | 7.60E+05 | 7.2E-06 | 1604 | 9.50E-12 |
| Candesartan | Human angiotensin II type 1 receptor | 2.60E+04 | 1.9E-04 | 61 | 7.40E-09 |
| Telmisartan | Human angiotensin II type 1 receptor | 9.10E+04 | 1.5E-04 | 77 | 1.70E-09 |
| Granisetron | 5-HT3 | 7.50E+05 | 7.5e-04 | 15 | 1.00E-09 |
| Lapatinib | ERBB2/EGFR | 1.30E+04 | 3.9e-05 | 296 | 3.00E-09 |
| Desloratadine | Histamine H1 receptor | 3.20E+04 | 3.2e-05 | 361 | 1.00E-09 |
| Amlodipine | L-type calcium channel | 1.50E+05 | 3.0E-04 | 39 | 2.00E-09 |
| Verapamil | L-type calcium channel | 9.20E+07 | 2.8E0 | 0.004 | 3.00E-08 |
| Phenytoin | Na<sup>+</sup> channel | 2.50E+04 | 5.0E-01 | 0.02 | 2.00E-05 |
| Amprenavir | HIV-1 protease | 4.40E+06 | 4.9E-03 | 2 | 1.10E-09 |
| Atazanavir | HIV-1 protease | 1.70E+06 | 6.9E-04 | 17 | 4.00E-10 |
| Indinavir | HIV-1 protease | 1.50E+06 | 1.6E-03 | 7 | 1.10E-09 |
| Lopinavir | HIV-1 protease | 6.60E+06 | 6.5E-04 | 18 | 1.00E-10 |
| Ritonavir | HIV-1 protease | 3.90E+06 | 2.2E-03 | 5 | 6.10E-10 |
| Saquinavir | HIV-1 protease | 8.20E+05 | 2.3E-04 | 50 | 3.20E-10 |
| Nelfinavir | HIV-1 protease | 6.60E+05 | 6.7E-04 | 17 | 1.60E-09 |
| Boceprevir | Hepatitis C virus nonstructural protease (NS3) | 2.40E+03 | 4.9E-05 | 236 | 2.00E-08 |
| Ciluprevir | Hepatitis C virus nonstructural protease (NS3) | 9.10E+05 | 7.3E-04 | 16 | 6.20E-09 |
| Telaprevir | Hepatitis C virus nonstructural protease (NS3) | 2.20E+03 | 9.6E-05 | 120 | 4.30E-08 |
| Deoxy-conformycin | Adenosine deaminase | 2.40E+06 | 4.8E-06 | 2,406 | 2.00E-12 |
| Efavirenz | HIV-1 reverse transcriptase | 1.40E+01 | 6.8E-05 | 170 | 5.00E-06 |
| Elvitegravir | HIV integrase | 2.10E+06 | 8.3E-03 | 1 | 4.00E-09 |
| Raltegravir | HIV integrase | 2.50E+05 | 2.5E-03 | 5 | 1.00E-08 |
| Methotrexate | Chicken dihydrofolate reductase | 3.70E+07 | 3.3E-04 | 35 | 9.00E-12 |
| Trimethoprim | Escherichia coli dihydrofolate reductase | 7.20E+07 | 1.4E-03 | 8 | 2.00E-11 |
| Oseltamivir | Viral neuraminidase | 7.90E+05 | 2.5E-04 | 46 | 3.10E-10 |
| Saxagliptin | Human dipeptidyl-peptidase IV | 1.60E+05 | 5.4E-05 | 214 | 3.50E-10 |
| Vildagliptin | Human dipeptidyl-peptidase IV | 8.70E+04 | 9.8E-04 | 12 | 1.10E-08 |
| Triclosan | Enoyl-ACP reductase and Francisella tularensis | 8.20E+06 | 4.2E-04 | 28 | 5.10E-11 |

These observations (and numbers 4 and 5 in particular) are consistent with our claim that high dynamic occupancy under non-equilibrium conditions found *in vivo* depends first and foremost on fast *k_on_*, and further suggest that the rates of buildup of many of the targeted binding sites are ≥ 1×10^-5^ s^-1^ (based on Table 1), assuming that *k_on_* was, in fact, wittingly or unwittingly optimized to *k_on_ss_* This contrasts with the claim of Dahl and Akrud [29] and Folmer [30] that the drugs were optimized on the basis of *K_d_* (noting that Folmer acknowledges the greater contribution of fast *k_on_* over slow *k_off_* to drug success [30]).

## Discussion

The fundamental mechanisms by which molecular properties are transduced into cellular structure-function are poorly understood. Molecules are an unruly lot, whose states are distributed randomly in the absence of transducible energy inputs to achieve maximal configurational entropy. The relevant energy inputs consist largely of: 1) H-bond losses and gains among solvating water molecules (versus other liquid solvents); and 2) continual/continuous production and decay of the participating species needed to maintain non-equilibrium conditions. Furthermore, dynamic counter-balancing is essential for overcoming the inherent susceptibility of exponential molecular state transitions to under- and overshooting. Our biodynamics theory, in which cellular structure-function is predicated on a molecular form of analog computing, is well-aligned with these principles. Multiple levels of integrated dynamic contributions underlying cellular function, dysfunction, pharmacodynamics, and drug PK are addressed by our theory, the implications of which are summarized in the following sections.

### The rates of change in binding partner levels/states are sensed via binding dynamics

The bound fraction of binding or catalytic sites can be calculated under strictly equilibrium conditions using the Hill and Michaelis-Menten equations, respectively. However, binding sites undergo transient buildup and decay driven by synthesis/degradation, translocation, or conformational state transitions to/from their binding-competent states. The lifetimes of many proteins range from single-digit minutes to hours [21] (e.g. PCSK9 is on the order of 5 min [24], and tumor cell line-derived NIK ranges from ~5-30 min [31]). The lifetimes of binding competent states can range from μs (e.g. mRNA) or ms (e.g. voltage-gated ion channels) to the lifetime of the molecule itself. Here, we have shown that dynamic occupancy is greatly influenced by the time window of binding site availability, where the SSO profile is only achieved when *k_on_* is on the order of *k_i_*. We assume that the association and dissociation rate constants for endogenous partners are kinetically tuned to the dynamic ranges of *B_total_*(*t*) and *L*(*t*). Perturbations to partner states/levels can be sensed through binding dynamics, and specifically, kinetic tuning/de-tuning of *C*(*t*) via:

1. Modulation of the maximum time-dependent binding partner levels (*B_total_*(*t*)) (the “extrinsic” rates [1]).
2. Slowing/speeding of *k_i_* and *k_-i_* thereby shifting the *k_on_* threshold for achieving the SSO profile (i.e. *k_0nss_*) or the qSSO profile.
3. Modulation of partner-specific association and dissociation rate constants (*k_on_* and *k_off_*) (the “intrinsic” rates [1]).

Sensitivity of cellular function to the bound state depends on:

1. The dynamic range in *B_total_*(*t*):

A. Constitutively low *B_total_*(*t*) levels may result in undetectable levels of the bound state.
B. Constitutively high *B_total_*(*t*) levels may result in reduced sensitivity of the bound state to changes in concentration of the other partner(s).
2. *k_on_* (subject to allosteric modulation, case-by-case).

A. Very fast *k_on_* (≫ *k_on_ss_*) diminishes the sensitivity of the bound state to changes in partner concentration.
B. Very slow *k_on_* (≫ *k_on_ss_*) may result in undetectable levels of the bound state at lower partner concentrations.

We assume that cellular systems operate within the binding sensitive regime, such that the buildup and decay of binding partners and the bound states thereof remain in phase over time. Although we have assumed a constant free ligand concentration in constructing our analytical dynamic occupancy expressions, it is apparent that this condition rarely occurs *in vivo* (noting that buildup and decay of the binding partners may proceed at different rates, and in an in- or out-of-phase relationship). As such, variable ligand concentration introduces potentially far greater kinetic demands on dynamic occupancy, which is likely underestimated in our approach.

### The implications of biodynamics for pharmacodynamics and drug discovery

Efficacious and toxic drug levels are conventionally defined in terms of the equilibrium scenario (equation 20), which rests on the assumptions that drug-target occupancy, drug exposure within the target compartment (typically intra- or extracellular), and target level are quasi-time-independent quantities, when in fact, they are not:

1. Biomolecular species underlying cellular function undergo continuous production/degradation and state transitions (including drug targets).
2. Drug levels build and decay based on PK principles (S7 Fig).
3. Drug-target binding builds and decays based on binding dynamics principles.
4. Drug-target binding alters the rates of dysfunction promoting processes (i.e. pharmacodynamics).

Pharmacodynamics may be taken as the mitigation of Yin-Yang imbalances [1] via the insertion of one or more exogenous MDEs into afflicted system(s) (where inhibition and stimulation effectively slow and speed target buildup, respectively). As for endogenous complexes:

1. Buildup of drug-target occupancy is on-rate driven, and the bound state must rebuild with each target buildup/decay cycle due to loss of the bound drug-target population.
2. On-rate may be driven by *k_on_* and/or free drug concentration.
3. The SSO profile is achievable, in practice, for kinetically tuned binding, in which *k_on_* is optimized to ≥ *k_0nss_*, as dictated by the lifetime of the target or binding site (Fig 18). Under these conditions, efficacious or toxic occupancy, which can conceivably vary from ≫ 50% to 95+% is expected at a free drug concentration = *n*·*K_d_* (e.g. n = 19 yields 95% occupancy), noting that the putative safe range of trappable hERG blocker occupancy ranges between 0-3% [4]. Conversely, achieving the qSSO profile (converging to ~95% occupancy) may require considerably larger n than the SSO profile.
4. In the absence of kinetic tuning, the SSO profile is unachievable via escalation of drug concentration, although the qSSO profile is conceivable at sub-toxic exposures. This constitutes a potential source of *in vitro-in vivo* disconnects and clinical failures due to loss of the therapeutic index. The worst-case scenario consists of a drug that is kinetically mistuned to its target, while being kinetically tuned to one or more off-targets, and that competes for the target with an endogenous kinetically tuned partner.
5. Residence time (i.e. In (2)/*k_off_* below the rate of binding site decay adds no benefit to occupancy (target dynamics are not considered by other workers advocating *k_off_* optimization [32,33]).

**Fig 18.**
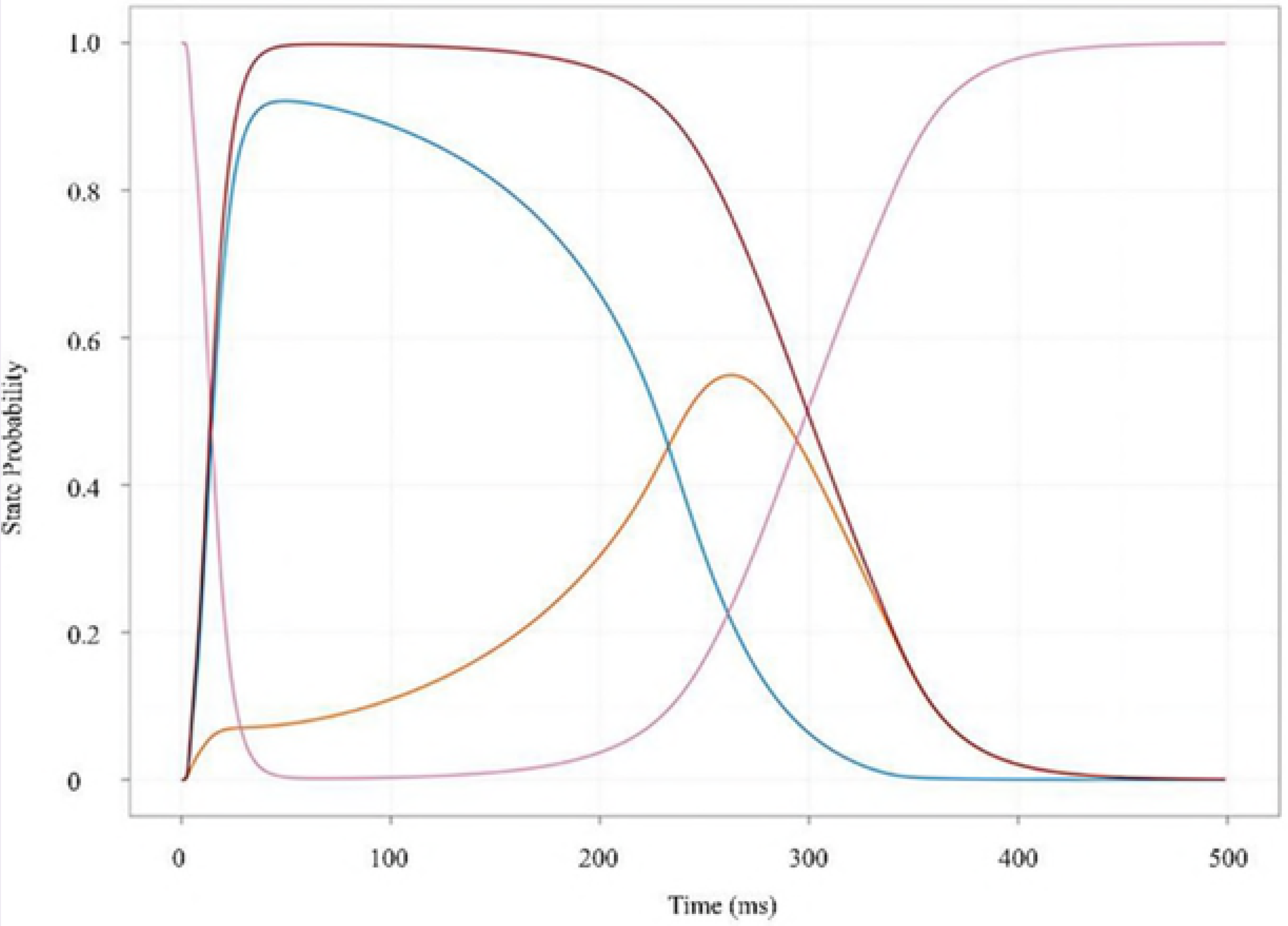
Optimization of dynamic occupancy is achieved by first speeding *k_on_* to ≥ *k_0nss_* (based on the binding site lifetime), followed by slowing *k_off_* to *k_-i_*, a process we refer to as “kinetic tuning.” <u>The ultimate objective is to achieve the highest occupancy at the lowest free drug concentration</u> (i.e. the SSO profile), affording the greatest chance for achieving a therapeutic index in humans.

## Conclusion

In our previous work [1], we outlined a first principles theoretical framework (referred to herein as biodynamics) that explains the general mechanisms of cellular analog computing on the basis of non-equilibrium biomolecular state transitions, as well as the key implications of our theory for:

1. The transformation of chemical systems into living cells capable of maintaining non-equilibrium conditions, harnessing exponential behavior, and exploiting water H-bond energy to generate state transition barriers.
2. The origin of disease-causing cellular dysfunction (Yin-Yang imbalances), and the pharmacological mitigation thereof (insertion of exogenous MDEs capable of fully or partially restoring the normal Yin-Yang balance).

The current work is focused on the interplay between two key biodynamics contributions consisting of molecular dynamics and binding dynamics. We used a simplified analytical model that qualitatively captures these contributions to explore the possible effects of non-equilibrium conditions (namely, binding site buildup and decay) on dynamic occupancy. Our results suggest that:

1. Achieving the SSO profile as a function of diminishing binding site lifetime depends on increasingly faster *k_on_* (noting that nSSO *>* the qSSO profile is achievable only at high *L* compared with SSO).
2. Nature has likely tuned *k_on_* to achieve SSO or qSSO binding for endogenous partners (i.e. maximal binding efficiency).

Our findings challenge the conventional equilibrium binding paradigm on which much of modern drug discovery is based, as follows:

1. Knowledge of target and binding site buildup and decay rates is essential for achieving the SSO profile on a non-trial-and-error basis.
2. Binding site lifetimes ranging from single-digit hours to milliseconds require increasingly faster *k_on_* to achieve the SSO profile (i.e. kinetic tuning versus potency optimization).
3. Increased drug exposure (i.e. PK) is a poor substitute for kinetically tuned binding, in which the qSSO profile is only asymptotically approached. As such, dose escalation during clinical trials is an extremely risky proposition from the safety standpoint.

## Acknowledgements

We gratefully acknowledge Xin Chen and Andrei Golosov for helpful discussions and comments on this manuscript.

## References

1. Pearlstein RARA, McKay DJJDJJ, Hornak V, Dickson C, Golosov A, Harrison T, et al. Building New Bridges between In Vitro and In Vivo in Early Drug Discovery: Where Molecular Modeling Meets Systems Biology. Curr Top Med Chem. 2017;17: 1–1. doi:10.2174/1568026617666170414152311

2. Daniel R, Rubens JR, Sarpeshkar R, Lu TK. Synthetic analog computation in living cells. Nature. Nature Publishing Group; 2013;497: 619–623. doi:10.1038/nature12148

3. Howe RM. Fundamentals of the Analog Computer. IEEE Control Syst Mag. 2005; 29–36.

4. Pearlstein RA, MacCannell KA, Erdemli G, Yeola S, Helmlinger G, Hu Q-Y, et al. Implications of dynamic occupancy, binding kinetics, and channel gating kinetics for hERG blocker safety assessment and mitigation. Curr Top Med Chem. 2016;16.

5. O’Hara T, Virág L, Varró A, Rudy Y. Simulation of the undiseased human cardiac ventricular action potential: model formulation and experimental validation. PLoS Comput Biol. 2011;7: e1002061. doi:10.1371/journal.pcbi.1002061

6. Willis WT, Jackman MR, Messer JI, Kuzmiak-Glancy S, Glancy B. A simple hydraulic analog model of oxidative phosphorylation. Med Sci Sports Exerc. 2016;48: 990–1000. doi:10.1249/MSS.0000000000000884

7. Alon U. An introduction to systems biology : design principles of biological circuits. Chapman & ; Hall/CRC; 2007.

8. Keener J, Sneyd J, editors. Mathematical Physiology [Internet]. New York, NY: Springer New York; 2009. doi:10.1007/978-0-387-75847-3

9. Keener JP. Mathematical physiology 2009 : systems physiology ii. 2nd revise. Springer; 2008.

10. Beard DA. Biosimulation : simulation of living systems [Internet]. Cambridge University Press; 2012. Available: https://books.google.com/books?id=PPl3yNpij9gC&pg=PR4&lpg=PR4&dq=Daniel+A.+Beard.+Biosimulation:+Simulation+of+Living+Systems.+Cambridge+University+Press,+Cambridge,+UK.,+2012.+ISBN+978-0-521-76823-8.&source=bl&ots=RKhaR4C26H&sig=WXszESobIMh3VrWyH9lDX8nuZ

11. Voit EO. A first course in systems biology. Garland Science; 2013.

12. Pearlstein RA, Hu Q-Y, Zhou J, Yowe D, Levell J, Dale B, et al. New hypotheses about the structure-function of proprotein convertase subtilisin/kexin type 9: Analysis of the epidermal growth factor-like repeat A docking site using WaterMap. Proteins Struct Funct Bioinforma. Wiley-Blackwell; 2010;78: 2571–2586. doi:10.1002/prot.22767

13. Pearlstein RA, Sherman W, Abel R. Contributions of water transfer energy to proteinligand association and dissociation barriers: Watermap analysis of a series of p38α MAP kinase inhibitors. Proteins Struct Funct Bioinforma. 2013;81. doi:10.1002/prot.24276

14. Tran Q-T, Williams S, Farid R, Erdemli G, Pearlstein R. The translocation kinetics of antibiotics through porin OmpC: Insights from structure-based solvation mapping using WaterMap. Proteins Struct Funct Bioinforma. 2013;81. doi:10.1002/prot.24185

15. Velez-Vega C, McKay DJJ, Kurtzman T, Aravamuthan V, Pearlstein RA, Duca JS. Estimation of solvation entropy and enthalpy via analysis of water oxygen-hydrogen correlations. J Chem Theory Comput. 2015;11. doi:10.1021/acs.jctc.5b00439

16. Pratt JM, Petty J, Riba-Garcia I, Robertson DHL, Gaskell SJ, Oliver SG, et al. Dynamics of Protein Turnover, a Missing Dimension in Proteomics. 2002; 579–591. doi:10.1074/mcp.M200046-MCP200

17. Hopper AK, Patel M, Furia BS, Peltz SW, Trotta CR, Trotta CR, et al. Proteome Half-Life Dynamics in Living Human Cells. 2011; 764–769.

18. Yen HS, Xu Q, Chou DM, Zhao Z, Elledge SJ. Global Protein Stability Profiling in Mammalian Cells. 2008;322: 918–924.

19. Rothman S. How is the balance between protein synthesis and degradation achieved ? 2010; 1–11.

20. Jiang X, Coffino P, Li X. Development of a method for screening short-lived proteins using green fluorescent protein. 2004;

21. Loriaux PM, Hoffmann A, Haugh JM. A Protein Turnover Signaling Motif Controls the Stimulus-Sensitivity of Stress Response Pathways. PLoS Comput Biol. 2013;9. doi:10.1371/journal.pcbi.1002932

22. Hoor M ten. “Are we there yet?” … Can Equilibrium ever be Achieved ? ChemEd NZ. 2009x; 7–8. Available: http://nzic.org.nz/chemed-nz/issue-archive.html

23. Invitrogen Corporation. Theory of Binding Data Analysis. Fluoresc Polariz Tech Resour Guid Chapter 7. 2008; 1–18. Available: www.invitrogen.com/drugdiscovery

24. Grefhorst A, Mcnutt MC, Lagace TA, Horton JD. Plasma PCSK9 preferentially reduces liver LDL receptors in mice. 2008;49: 1303–1311. doi:10.1194/jlr.M800027-JLR200

25. Benjannet S, Rhainds D, Essalmani R, Mayne J, Wickham L, Jin W, et al. NARC-1/PCSK9 and Its Natural Mutants ZYMOGEN CLEAVAGE AND EFFECTS ON THE LOW DENSITY LIPOPROTEIN (LDL) RECEPTOR AND LDL CHOLESTEROL*. 2004; doi:10.1074/jbc.M409699200

26. Schulz R, Schluter KD, Laufs U. Molecular and cellular function of the proprotein convertase subtilisin/kexin type 9 (PCSK9). Basic Res Cardiol. 2015;110. doi:10.1007/s00395-015-0463-z

27. Gustafsen C, Olsen D, Vilstrup J, Lund S, Reinhardt A, Wellner N, et al. Heparan sulfate proteoglycans present PCSK9 to the LDL receptor. Nat Commun. Springer US; 2017;8: 1–14. doi:10.1038/s41467-017-00568-7

28. Stork D, Timin EN, Berjukow S, Huber C, Hohaus a, Auer M, et al. State dependent dissociation of HERG channel inhibitors. Br J Pharmacol. 2007;151: 1368–1376. doi:10.1038/sj.bjp.0707356

29. Dahl G, Akerud T. Pharmacokinetics and the drug-target residence time concept. Drug Discov Today. 2013;18: 697–707. doi:10.1016/j.drudis.2013.02.010

30. Folmer RHAA. Drug target residence time: a misleading concept. Drug Discov Today. 2017;00: 1–5. doi:10.1016/j.drudis.2017.07.016

31. Annunziata CM, Davis RE, Demchenko Y, Bellamy W, Zhan F, Lenz G, et al. Frequent engagement of the classical and alternative NF-κB pathways by diverse genetic abnormalities in multiple myeloma. Cancer Cell. 2007;12: 115–130. doi:10.1016/j.ccr.2007.07.004.Frequent

32. Copeland RA, Pompliano DL, Meek TD. Drug-target residence time and its implications for lead optimization. Nat Rev Drug Discov. 2006;5: 730–739. doi:10.1038/nrd2082

33. Schoop A, Dey F. On-rate based optimization of structure-kinetic relationship - Surfing the kinetic map. Drug Discov Today Technol. Elsevier Ltd; 2015;17: 9–15. doi:10.1016/j.ddtec.2015.08.003

